# Paralemmin 2 is a shared, alternatively-spliced regulator of axon initial segments and paranodal junctions in oligodendrocytes

**DOI:** 10.64898/2026.08.18.745582

**Authors:** Xiaoyun Ding, Yuki Ogawa, Victoria L. Palfini, Wei Zhang, Andrew P. Anderson, Lucia Gancedo-López, Zian Liao, Zihan Yang, Seth G. Haddix, Yueheng Xing, Duc V. M. Nguyen, Jackson R. Curtis, Davis J. Church, Yu Wu, Antrix Jain, Alexander B. Saltzman, Juan A. Oses-Prieto, Matthew Torre, Trinh T. Tat, Daniel C. Kraushaar, Alma L. Burlingame, Anna Malovannaya, Lulu Shang, Pengfei Liu, Jiaxing Li, Yudong Gao, Matthew N. Rasband

**Author notes:** Lead contact: Matthew Rasband.

## Abstract

Nervous system function depends on highly specialized axonal membrane domains. For example, axon initial segments (AIS), nodes of Ranvier, and paranodal junctions are essential for action potential initiation and saltatory conduction. These domains use shared molecular machinery converging on the master scaffolding protein AnkyrinG (AnkG). Using endogenous AnkG-TurboID proximity proteomics, we identify Paralemmin 2 (Palm2), a previously uncharacterized CNS protein, as a shared component of the AIS and nodes of Ranvier in neurons, and paranodes in oligodendrocytes. Palm2 localizes transiently to the AIS and nodes during development, but is found at paranodes throughout life, reflecting cell type-specific splicing of a lipid-modified domain both necessary and sufficient for AnkG interaction. Mice lacking oligodendroglial Palm2 assemble normal paranodes during development, but aged mice lose paranode integrity due to loss of paranodal AnkG and Neurofascin-155. Our findings identify Palm2 as a lipid raft-associated regulator of axonal and oligodendroglial membrane domains and reveal cell type-specific protein variants as a recurring principle for specializing axonal and glial membrane architectures.

## Introduction

Axons and myelinating oligodendrocytes enable efficient and rapid action potential (AP) propagation through the assembly and maintenance of specialized excitable membrane domains. For example, the axon initial segment (AIS) is highly enriched with ion channels, cell adhesion molecules (CAMs), and cytoskeletal proteins that together initiate APs in response to synaptic input ^1,2^. In myelinated axons, APs are regenerated and propagated by the high density of ion channels found at nodes of Ranvier. Node assembly and maintenance, and AP propagation also depend on paranodal junctions that flank each node ^3^. Paranodal junctions, formed between the axon and myelinating oligodendrocyte, are the largest adhesion junction in vertebrates and function as the direct point of contact for axon-glia communication. Disruption of these domains contributes to a broad range of neurological disorders and injuries ^4,5^.

Remarkably, many of the core proteins that comprise these domains are shared between neurons and oligodendrocytes. For example, splice variants of the scaffolding protein AnkyrinG (AnkG) and the CAM Neurofascin function in both neurons and oligodendrocytes, yet occupy structurally and functionally distinct domains. AnkG, the master organizer of the AIS, is encoded by the gene *Ank3*. Its large neuronal isoforms organize and cluster AIS and nodal ion channels, while a smaller oligodendroglial variant facilitates assembly and maintenance of the paranodal junction through binding to the CAM Neurofascin, encoded by the gene *Nfasc* ^6,7^. At AIS and nodes, AnkG binds to the larger 186 kDa neuronal splice variant (NF186); and at paranodes, AnkG binds a smaller 155 kDa oligodendroglial variant (NF155). NF155 is an essential component of paranodal junctions, and its loss prevents paranodal junction assembly ^8-10^. These observations suggest that diversifying shared molecular machineries through alternative splicing may be a fundamental strategy used by neurons and glia to generate distinct membrane domains.

Recently, the development and application of proteomic tools – especially proximity biotinylation – has begun to reveal the AIS and nodal proteomes. The discovery of new AIS and nodal proteins has fueled mechanistic insights into the structure, function, assembly, and maintenance of these important domains ^11-13^. For example, proximity biotinylation revealed that AIS Kv1 channels can be clustered and dephosphorylated through the AIS-enriched proteins Scrib and Ppp2r2c, respectively ^14,15^. In contrast, and despite the importance of paranodes for nervous system function, no new paranodal protein has been identified in more than two decades.

To further capitalize on the power of *in vivo* proximity proteomics, we generated a Cre-dependent endogenous *Ank3*^*LSL-TurboID*^ knock-in mouse, which enables cell-type and domain specific proximity proteomic analyses. Using this mouse, we profiled four distinct AnkG-enriched domains and identified Paralemmin 2 (Palm2) as a shared component of the AIS, nodes, and paranodes. We observed both temporal and cell-type specific regulation of its expression in neurons and oligodendrocytes. Neuronal Palm2 is transiently found at the AIS where it participates in the assembly of the AIS, but subsequently undergoes alternative splicing and transcriptional readthrough that prevents Palm2’s AIS localization later in development. In contrast, oligodendroglial Palm2 is constitutively expressed and accumulates at paranodes. We find that Palm2 is a lipid raft-associated protein that regulates the maintenance and stability of paranodal AnkG and NF155. Although paranodes form normally in mice lacking paranodal Palm2, aged mice have disrupted paranodal junctions due to the loss of paranodal NF155 and AnkG. Thus, Palm2 joins Neurofascin and AnkG as core, alternatively spliced proteins at AIS, nodes of Ranvier, and paranodal junctions that are necessary for the structural and functional organization of myelinated axons throughout life.

## Results

### AnkG proximity proteomes

Neurons and glia are highly polarized cells with specialized membrane domains that perform distinct physiological functions. These domains can exhibit strikingly different molecular architectures but share some core protein components ^3^. How common molecular machinery is repurposed to assemble and maintain distinct membrane domains is poorly understood. To begin to address this question, we focused on AnkG, a master regulator and scaffolding protein that is enriched at and plays key roles in the assembly and maintenance of several specialized membrane domains including: AIS, nodes of Ranvier, paranodal junctions in oligodendrocytes, and the peri-junctional zone (PJZ) of the neuromuscular junction (NMJ) ^7,16-18^. We reasoned that AnkG could serve as a common molecular entry point to systematically compare the proteomes of these distinct membrane domains.

To this end, we generated a Cre-dependent TurboID knock-in mouse line at the endogenous *Ank3* locus (*Ank3*^*LSL-TurboID*^), with the goal of cell-type-specific profiling of AnkG-proximity proteomes (Fig. 1A). To determine the efficiency and specificity of biotinylation in *Ank3*^*LSL-TurboID*^ mice, we crossed *Ank3*^*LSL-TurboID*^ mice with lineage-specific Cre lines, and injected them with biotin, collected nervous system tissues and labeled AIS, nodes, and the PJZ of the NMJ using streptavidin and domain specific markers; we used AnkG for AIS (Fig. 1B), Caspr for paranodes (Fig. 1C), NF186 for nodes of Ranvier (Fig. 1D), and α-Bungarotoxin (α-BTX) for the NMJ (Fig. 1E). Notably, in the absence of Cre-mediated recombination, streptavidin labeling was detectable and enriched at cortical AIS and optic nerve nodes of Ranvier (Fig. S1A), indicating leakiness of the conditional allele due to direct splicing onto the duplicated splice acceptor downstream of the STOP cassette^19^. However, little streptavidin signal was observed outside neurons under baseline conditions, and *Vglut2*^*Cre*^*;Ank3*^*LSL-TurboID*^ mice markedly increased biotinylation at AIS and nodes of Ranvier compared with non-Cre controls (Fig. 1B, C and Fig. S1A). This suggests that despite neuronal leakiness of the system, Cre-mediated recombination could yield stronger cell-type specific labeling in other cell types.

**Figure 1.**
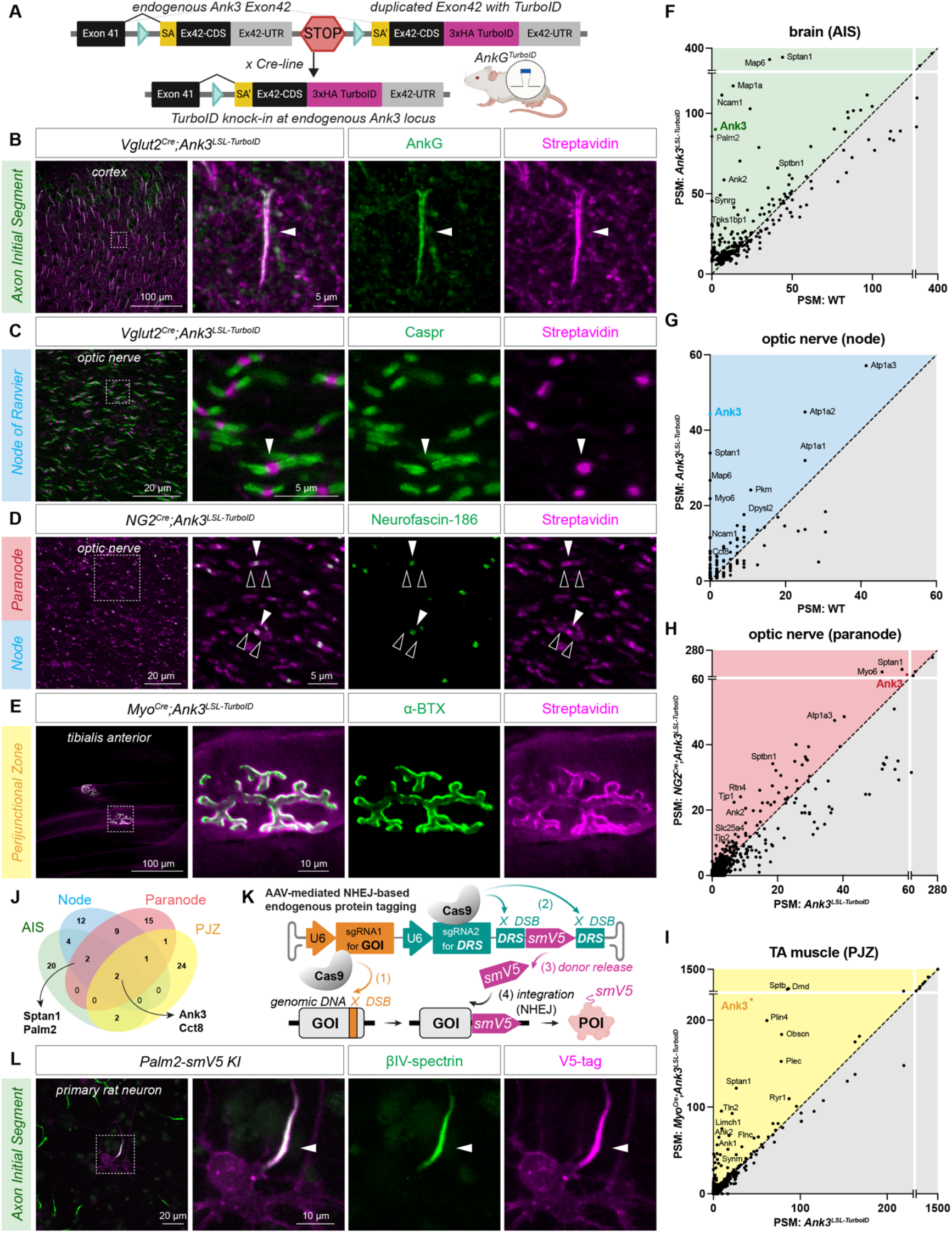
Endogenous AnkG-proximity proteomics identifies Palm2 as a shared component of axon initial segments, nodes of Ranvier, and paranodes. (A) Schematic of the *Ank3*^*LSL-TurboID*^ conditional knock-in allele. (B) Immunostaining of adult cortex showing efficient and specific biotinylation at the AIS (streptavidin, magenta), colocalizing with AnkG (green) in *Vglut2*^*Cre*^*;Ank3*^*LSL-TurboID*^ mice. (C) Immunostaining of adult optic nerve showing efficient and specific biotinylation of the nodes (streptavidin, magenta), flanked by paranodal Caspr (green) in *Vglut2*^*Cre*^*;Ank3*^*LSL-TurboID*^ mice. (D) Immunostaining of adult optic nerve showing additional biotinylation at paranodes (magenta; black arrowheads) flanking nodes of Ranvier (NF186, green; white arrowheads) in *NG2*^*Cre*^*;Ank3*^*LSL-TurboID*^ mice. (E) Immunostaining of adult tibialis anterior muscle showing efficient and specific biotinylation of the perijunctional zone (PJZ) at the neuromuscular junction (NMJ) in *Myo*^*Cre*^*;Ank3*^*LSL-TurboID*^ mice. (F-I) Peptide spectral matches (PSMs) for candidate AIS proteins (F), nodal proteins (G), paranodal proteins (H), and PJZ proteins (I). (J) Venn diagram of the top 30 candidate proteins identified at the AIS (green), node (blue), paranode (red), and PJZ (yellow). (K) Schematic of AAV-mediated endogenous protein tagging with spaghetti monster-V5 (smV5) for antibody-independent validation of protein localization. (L) Endogenous Palm2-smV5 knock-in (KI) signals are enriched at the AIS of cultured neurons. Arrowheads indicate colocalization of Palm2-smV5 (magenta) with βIV-spectrin (green). Scale bars are indicated in each panel.

To define neuronal AnkG-proximity proteomes at AIS and nodes of Ranvier, we captured biotinylated proteins and performed mass spectrometry on both cortex and optic nerve samples from WT, *Ank3*^*LSL-TurboID*^, and *Vglut2*^*Cre*^*;Ank3*^*LSL-TurboID*^ mice (Figs. 1F, G and Figs. S1B, C; Supplemental Data 1). We observed enrichment of many known AIS and nodal proteins, including AnkG (*Ank3*), αII-spectrin (*Sptan1)* and Map6 (*Map6*) (Figs. 1F, G and Figs. S1B, C, E, F, and S1I), validating the specificity of the approach ^11^. Interestingly, Na^+^ channels, K^+^ channels, and NF186, known AnkG-binding proteins, were not among the most enriched biotinylated proteins. TurboID has a biotinylation range of ∼10 nm ^20^, and *Ank3*^*LSL-TurboID*^ has TurboID fused to the C-terminus of AnkG. Thus, canonical AIS and nodal proteins that bind to the N-terminal ankyrin repeats (e.g. Na^+^ channels and NF186) are located far from the biotin ligase and may not be heavily biotinylated.

To define glial AnkG-proximity proteomes at the paranodal junction, we crossed *Ank3*^*LSL-TurboID*^ mice with the oligodendrocyte-lineage specific Cre line *NG2*^*Cre*^ (Fig. 1D). We observed robust paranodal biotinylation (Fig. 1D, black arrowheads); in addition, we observed leaky expression in some neurons as indicated by nodal streptavidin (Fig. 1D, white arrowheads). To differentiate oligodendrocyte-enriched proteins from neuronal proteins, we compared *NG2*^*Cre*^*;Ank3*^*LSL-TurboID*^ mice with both *Ank3*^*LSL-TurboID*^ controls (Figs. 1H, S1G, J, and Supplemental Data 1) and *Vglut2*^*Cre*^*;Ank3*^*LSL-TurboID*^ mice (Fig. S1D). We identified proteins more enriched within the oligodendrocyte proteome than the neuronal proteome. For example, in addition to AnkG, we found enrichment for known oligodendroglial paranodal proteins including αII-spectrin (*Sptan1)*, βII-spectrin (*Sptbn1*), and AnkyrinB (*Ank2*) ^3^.

As one last AnkG-enriched domain not found in neurons or glia, we also investigated the PJZ. We crossed *Ank3*^*LSL-TurboID*^ mice with the skeletal muscle-specific Cre line *Myo*^*Cre*^ to biotinylate proteins in the vicinity of muscle AnkG (Fig. 1E). We then purified biotinylated proteins and performed mass spectrometry (Figs. 1I, S1H, K, and Supplemental Data 1). We found known PJZ enriched proteins, including βI-spectrin (*Sptb*), Dystrophin (*Dmd*) and Perilipin 4 (*Plin4*) ^21^.

To identify common molecular machineries used at these different AnkG-enriched membrane domains, we cross compared the top 30 candidates from the four membrane domains (AIS, node, paranode, and PJZ; Fig. 1J; see also source data), and found AnkG (*Ank3)* and the chaperonin subunit protein Cct8 (*Cct8)* enriched in all four groups, while the cytoskeletal protein αII-spectrin (*Sptan1*) and a poorly characterized membrane-associated protein, Paralemmin2 (*Palm2*), are found enriched in the CNS datasets but not at the PJZ. Together our results show that *Ank3*^*LSL-TurboID*^ mice, combined with a variety of Cre-lines, can reveal cell-type and membrane-domain specific proximity proteomes including proteins that may be common among these domains.

### Palm2 is a bona fide AIS protein

Palm2 is a member of the paralemmin family of proteins ^22^. Although it is highly expressed in the brain, its role is unknown. Palm2 undergoes extensive lipid modification including prenylation and palmitoylation ^23^, suggesting it is anchored to the membrane and may participate in membrane organization and remodeling. Palm1 was previously reported to bind to βII-spectrin and contribute to the organization of the axonal membrane-associated periodic skeleton (MPS) ^24^. Among the candidates identified in the intersecting proteomes, we focused on Palm2 since the AIS has a robust MPS and is enriched with βIV-spectrin and palmitoylated proteins ^25,26^. To confirm that Palm2 is an AIS protein we performed three separate experiments. First, we used AAV, CRISPR/Cas9, and non-homologous end joining (NHEJ)-based genome editing (Fig. 1K) ^27^ to add a spaghetti monster-V5 tag (smV5) to endogenous neuronal Palm2. After transduction of primary rat neurons with viruses containing Cas9 and Palm2-specific gRNAs, we found smV5-tagged Palm2 highly enriched at the AIS and colocalizing with AIS βIV-spectrin (Fig. 1L). Second, we transduced primary neurons with AAVs expressing a V5-tagged Palm2 under the CamKII promoter. Like the tagged endogenous Palm2, exogenous Palm2 is enriched at the AIS where it colocalizes with βIV-spectrin (Fig. 2A). Third, we obtained three commercially available antibodies recognizing different regions of Palm2 and performed immunofluorescence labeling of primary neurons at 4, 7, and 21 days *in vitro* (DIV; Figs. 2B and S2A). All three antibodies showed enrichment of Palm2 at the AIS and colocalized with AnkG (Figs. 2B and S2B). We quantified the polarity index of Palm2 and found a consistent enrichment at the AIS from DIV4 to DIV21, although the polarity index showed a significant decrease at DIV21 (Fig. 2C).

**Figure 2.**
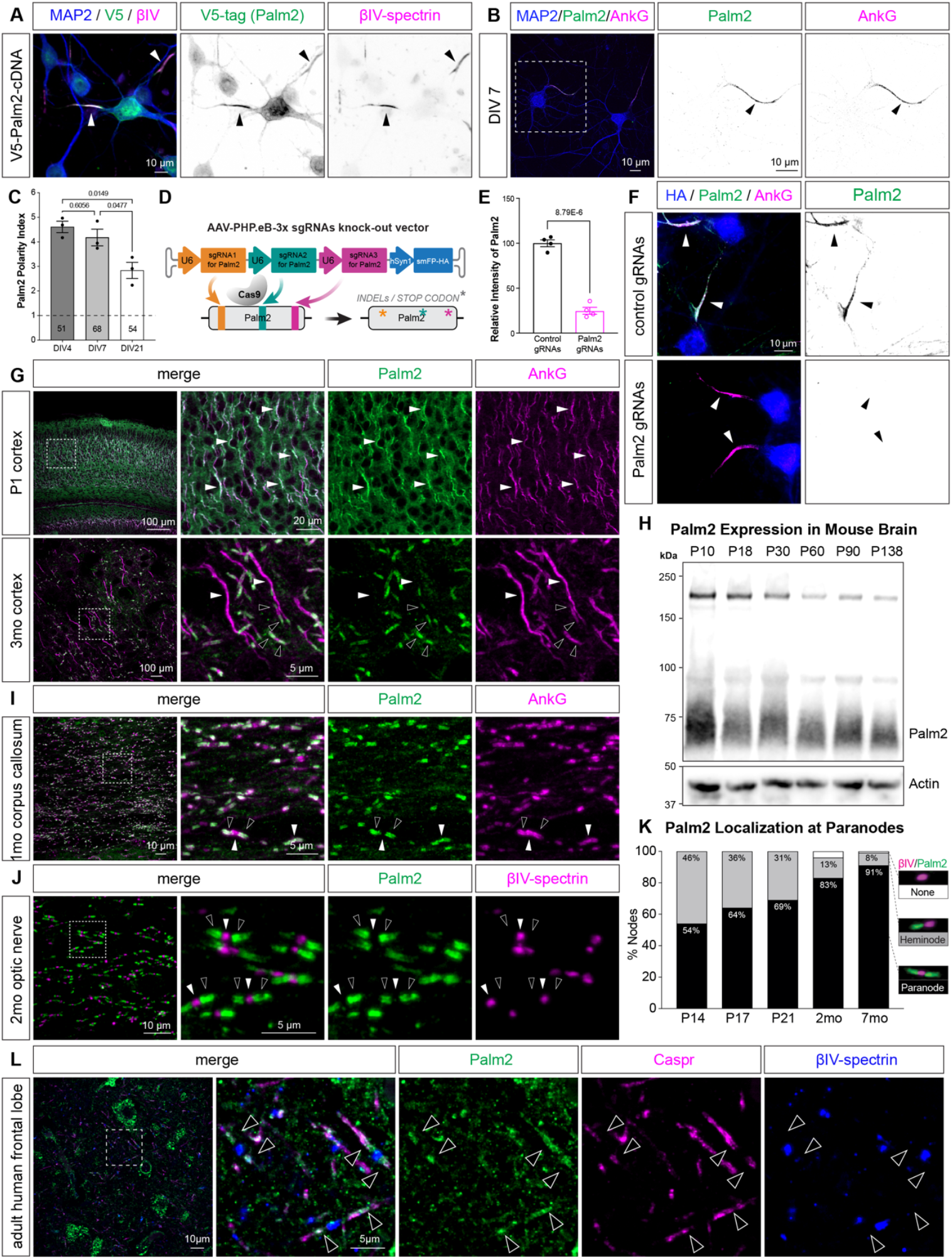
Palm2 is transiently localized to the AIS but constitutively found at paranodes. (A) V5-tagged exogenous Palm2 localizes to the AIS. Arrowheads indicate colocalization of V5-Palm2 (green) with βIV-spectrin (magenta). (B) Immunostaining of endogenous Palm2 in cultured cortical neurons shows colocalization of Palm2 (green) with AnkG (magenta). Arrowheads indicate the AIS. (C) Palm2 polarity index (AIS/dendrite fluorescence intensity ratio) at 4, 7, and 21 days in vitro (DIV). A polarity index >1 indicates AIS enrichment. N = 3 independent cultures, with the total number of neurons indicated within each bar. Error bars represent mean ± SEM; One-way ANOVA with multiple comparisons. (D) Schematic of AAV-mediated knockout (KO) by CRISPR/Cas9 using triple gRNAs. (E) Palm2 fluorescence intensity following AAV/CRISPR-mediated KO in primary mouse neurons. N = 4 independent cultures. Error bars represent mean ± SEM. Statistical comparisons were performed using Welch’s t-test; p value is shown on the figure. (F) Immunostaining of Palm2 (green) and AnkG (magenta) in HA-positive (blue) transduced primary neurons. Arrowheads indicate the AIS. (G) Immunostaining of Palm2 (green) and AnkG (magenta) in P1 and 3-month-old mouse cortex. White arrowheads indicate AIS. Black arrowheads indicate paranodal Palm2. (H) Immunoblotting of Palm2 in developing and adult mouse brain homogenates. (I) Immunostaining of Palm2 (green) and AnkG (magenta) in 1-month-old mouse corpus callosum. White arrowheads indicate nodes of Ranvier. Black arrowheads indicate paranodes. (J) Immunostaining of Palm2 (green) and βIV-spectrin (magenta) in 2-month-old mouse optic nerve. White arrowheads indicate nodes of Ranvier. Black arrowheads indicate paranodes. (K) Palm2 localization relative to βIV-spectrin during node and paranode development. Mature paranodes are defined by nodal βIV-spectrin (magenta) flanked by Palm2 (green) on both sides of the node (black bars), heminodes are defined by βIV-spectrin flanked by Palm2 only on one side of the node (gray bars), and some nodes only had βIV-spectrin without any flanking Palm2 (white bars). (L) Immunostaining of Palm2 (green), Caspr (magenta), and βIV-spectrin (blue) in postmortem adult human frontal lobe. Arrowheads indicate paranodes. Scale bars are indicated in each panel.

Finally, to validate the specificity of the Palm2 antibodies, we performed AAV and CRISPR/Cas9-mediated knockout of Palm2 in cultured neurons using a combination of three independent gRNAs (3x gRNA; Fig. 2D) ^28^. We observed loss of Palm2 immunofluorescence labeling in neurons transduced with Palm2 gRNAs compared to control gRNAs (Figs. 2E, F). Together, these results confirm that Palm2 is an AIS-enriched protein.

### Palm2 localization at the AIS is transient during development

To determine if Palm2 is found at the AIS *in vivo*, we immunolabeled cortical brain sections at different ages. At postnatal day 1 (P1), Palm2 strongly colocalized with AnkG in the cortex, indicating it is present at the developing AIS *in vivo* (Fig. 2G, white arrowheads). However, with increasing age the AIS Palm2 immunofluorescence became gradually less detectable and was completely absent from cortical AIS in 3-month-old mice (Fig. 2G, white arrowheads). Despite this apparent reduction in AIS labeling, immunoblots of brain membrane fractions across postnatal development showed a continuous increase in the total Palm2 protein in the brain, with only a slight decrease at P30 (Figs. 2H and S2C). The increase in Palm2 protein paralleled an increase in Neurofascin, the latter reflecting myelination and the accompanying maturation of nodes of Ranvier and paranodal junctions (Fig. S2C). Together, these results show that despite increasing Palm2 protein expression during development, it is found only transiently at the AIS.

### Palm2 is enriched at paranodes throughout life

Although Palm2 was absent from the AIS in adult cortex, we observed abundant Palm2-immunolabeling that overlapped with paranodal and oligodendroglial AnkG (Fig. 2G, black arrowheads). To determine if Palm2 is also a paranodal protein, we examined different white matter tracts both during development and in adults. Consistent with the staining observed in cortex, Palm2 robustly colocalized with paranodal AnkG in the corpus callosum (Fig. 2I), and flanked nodal βIV-spectrin in the optic nerve (Fig. 2J). Immunostaining using three different Palm2 antibodies generated against different regions of Palm2 (Fig. S2A) showed differing sensitivities to AIS, nodes, and paranodal junctions (Fig. S2I). Importantly, the Palm2 mouse monoclonal antibodies showed very high specificity for paranodes with very little or no labeling of nodal Palm2 (Figs. 2J and S2D-F). Immunostaining of paranodal Palm2 during developmental myelination and in adults showed that Palm2 is enriched at heminodes and remains at mature paranodes from P14 through 7 months of age (Fig. 2K and S2D-I). Like NF155, Palm2 was detected at over 80% of paranodes examined during both development and in adulthood. However, immunostaining of peripheral nerves using multiple Palm2 antibodies showed only weak nodal immunoreactivity at one month of age, and no paranodal Palm2 (Fig. S2E). Immunostaining of peripheral nodes of Ranvier from 4-month-old mice showed even less nodal enrichment for Palm2; the antibody with the strongest CNS paranodal Palm2 signal (Abnova) showed no PNS paranodal immunoreactivity (Fig. S2E). Together, these findings suggest that Palm2 is a core component of CNS, but not PNS, paranodes. Finally, to determine if Palm2 is also found at paranodes in human brain, we immunolabeled postmortem adult human frontal cortex. Nodes and paranodes were labeled for βIV-spectrin and Caspr, respectively. Similar to mouse CNS tissue, Palm2 was enriched at paranodes, and partially colocalized with Caspr (Fig. 2L). These findings show that Palm2 is a paranodal protein in both mouse and human, suggesting it has an evolutionarily conserved role.

### Alternative splicing of Palm2 regulates its localization

Consistent with the immunostaining results, RNA sequencing data show that *Palm2* is expressed in neurons and oligodendrocytes in both mice and humans (Figs. S3A-C) ^29,30^. However, it is unclear why Palm2 is lost from the AIS with increasing age but remains at paranodes throughout life. To answer this question, we examined *Palm2’s* gene structure (Fig. 3A) and potential regulation. In addition to the Palm2 protein, *Palm2* can form a naturally occurring readthrough product with its downstream gene *Akap2*, which is annotated as *Pakap* (Fig. 3B) ^31^. The two genes consist of a total of 12 exons, with the first 7 exons encoding *Palm2* and exons 9-12 encoding *Akap2*. In the event of gene readthrough, the fusion product *Pakap* lacks the last exon of *Palm2* (exon 7) and incorporates the downstream exon 8 in addition to *Akap2*. To determine if the protein products from *Palm2, Akap2*, or *Pakap* can localize to the AIS, we transduced primary neurons with AAVs expressing V5 tagged Palm2, Akap2, and Pakap (Fig. 3C). We found that V5-tagged Palm2 was highly enriched at the AIS (Fig. 2A), but neither V5-tagged Akap2 nor Pakap were found at the AIS, and their polarity index showed no AIS enrichment (Figs. 3D, arrowheads and 3E). To further define the domain of Palm2 responsible for its AIS localization, we generated internal deletion constructs lacking individual exons (exons 2-6) of Palm2 (Fig. 3F). None of these deletions affected AIS localization (Fig. 3G) or enrichment (Fig. 3H). However, deletion of the last 50 amino acids from exon 7 of Palm2 (Fig. 3F; Palm2 Δex7C) blocked the AIS clustering of Palm2 (Figs. 3G, H). Finally, we used CRISPR/Cas9 and gRNAs to target only exon 7. Targeting exon 7 alone is sufficient to disrupt Palm2’s AIS localization (Figs. S3D, E). Together, these results suggest that neither Akap2 nor Pakap are enriched at the AIS, and that inclusion of exon 7 in *Palm2* is required for the AIS localization of Palm2 protein.

**Figure 3.**
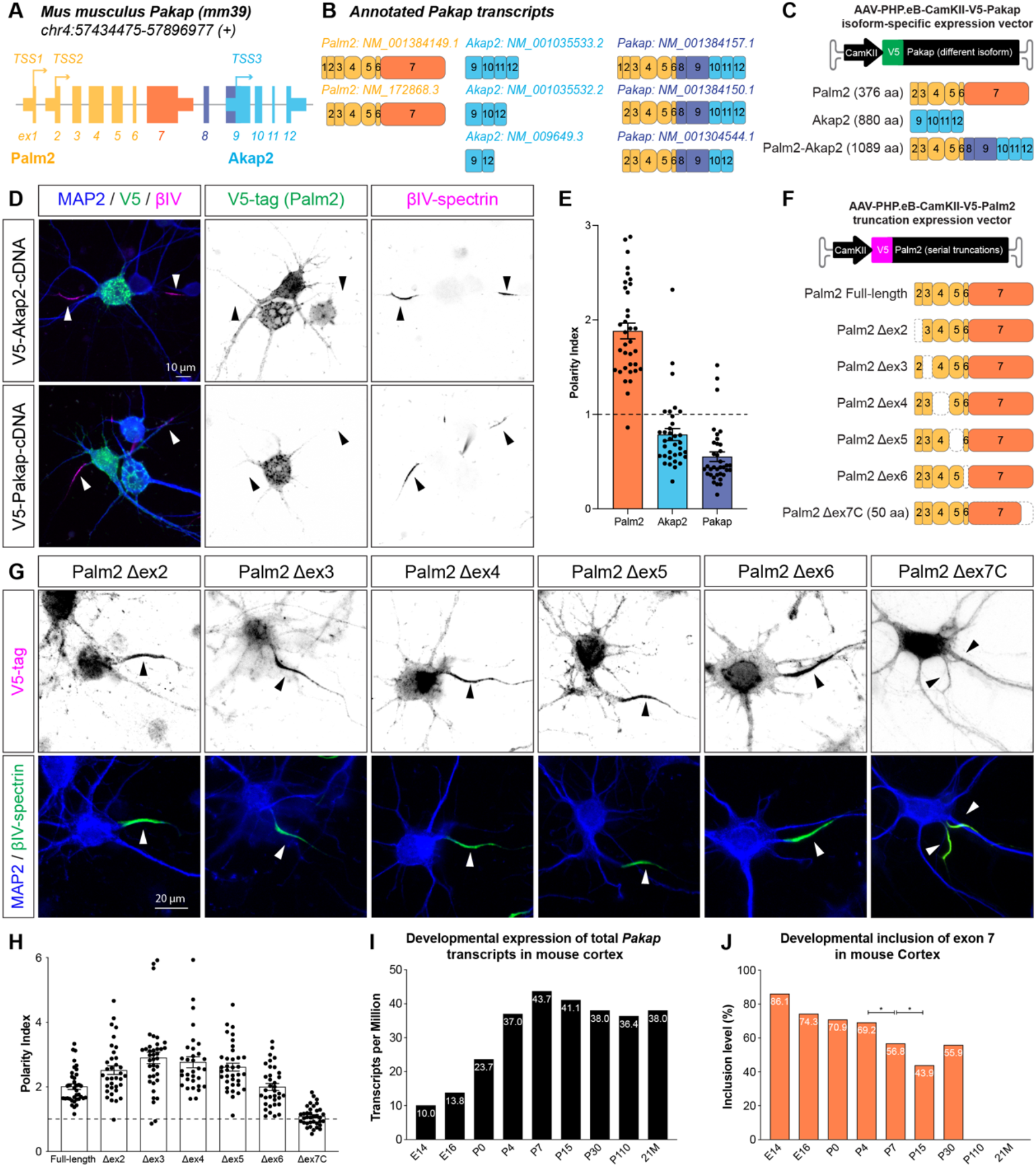
AIS localization of Palm2 is regulated by alternative splicing. (A) Illustration of the genomic structure of the Pakap gene including 12 exons, with two transcription start sites (TSSs) in the upstream Palm2 gene (exon 1-7; yellow), and one TSS in the downstream Akap2 gene (exon 9-12; light blue). (B) Illustration of all annotated Pakap transcripts and their exon usage, with two Palm2 transcripts in yellow (Palm2-exclusive exon in orange), three Akap2 transcripts in light blue, and three Pakap fusion transcripts (Pakap-specific exon in dark blue). (C) Schematic of AAV-mediated expression of different Pakap isoforms (Palm2, Akap2, and Pakap). (D) Immunostaining of V5-tagged Akap2 or Pakap (green), βIV-spectrin (magenta), and MAP2 (blue) in cultured neurons. Arrowheads indicate the AIS. (E) Polarity index of exogenously expressed Palm2, Akap2, and Pakap in cultured neurons. A polarity index >1 indicates AIS enrichment. Error bars represent mean ± SEM. Each dot represents an individual neuron. (F) Schematic of AAV-mediated expression of truncated Palm2 constructs lacking individual exons (Δex2–Δex6) or the C-terminal 50 amino acids encoded by exon 7 (Δex7C). (G) Immunostaining of V5-Palm2 variants (magenta), βIV-spectrin (green), and MAP2 (blue) in cultured neurons. Arrowheads indicate the AIS. (H) Polarity index of truncated Palm2 in cultured neurons. A polarity index >1 indicates AIS enrichment. Error bars represent mean ± SEM. Each dot represents an individual neuron. (I) Expression level of total Pakap transcript in developing and adult mouse cortex, obtained from Cortexa. Read counts per million are indicated within each bar. (J) Inclusion level of exon 7 during alternative splicing in developing and adult mouse cortex, obtained from Cortexa. Percentages of inclusion are indicated within each bar. One-way ANOVA with multiple comparisons. Asterisks indicate statistical significance. Scale bars are indicated in each panel.

Analysis of *Pakap’s* alternative splicing in mouse cortical neurons showed the expression of total *Pakap* transcripts increases during development and remains stable in adults (Fig. 3I) ^32,33^. Strikingly however, inclusion of exon 7 progressively declines during development and becomes undetectable by P110 (Fig. 3J). The loss of exon 7 from *Pakap* transcripts matches the loss of AIS-localized Palm2 (Fig. 2G). Thus, we conclude that Palm2 is lost from the AIS because developmental splicing of *Palm2* results in production of *Pakap* lacking exon 7, which is required for its AIS localization.

### Ankyrins are required for Palm2 localization to AIS and paranodes

How is Palm2 recruited to AIS, nodes, and paranodes? Since Palm2 was identified using AnkG-proximity proteomics (Fig. 1), we tested if Palm2’s localization depends on AnkG. To test this for AIS, we performed CRISPR-Cas9 3x gRNA knockout of AnkG from primary cultured neurons and confirmed the loss of Palm2 from AIS in AnkG-negative cells compared to AnkG-positive control neurons (Figs. 4A, B). The loss of AnkG did not eliminate Palm2 protein, but rather caused the loss of clustering and redistribution of Palm2 throughout the neuron. To test if paranodal clustering of Palm2 requires AnkG, and to confirm that paranodal Palm2 is in myelinating oligodendrocytes rather than the axon, we ablated AnkG from oligodendrocytes using *NG2*^*Cre*^*;Ank3*^*F/F*^ mice. Although these mice showed a significant decrease in paranodal Palm2 (Fig. S3F), it was not completely absent from the paranode. Since paranodal AnkB can partially compensate for the loss of paranodal AnkG, we examined paranodal Palm2 clustering in *CNP*^*Cre*^*;Ank2*^*F/F*^*;Ank3*^*F/F*^ mice ^7^. In mice lacking both AnkG and AnkB in oligodendrocytes, we observed the complete loss of paranodal Palm2 compared to control mice (Fig. 4C, black arrowheads). Together, these results show that in both neurons and oligodendrocytes Palm2 clustering requires Ankyrin scaffolds.

**Figure 4.**
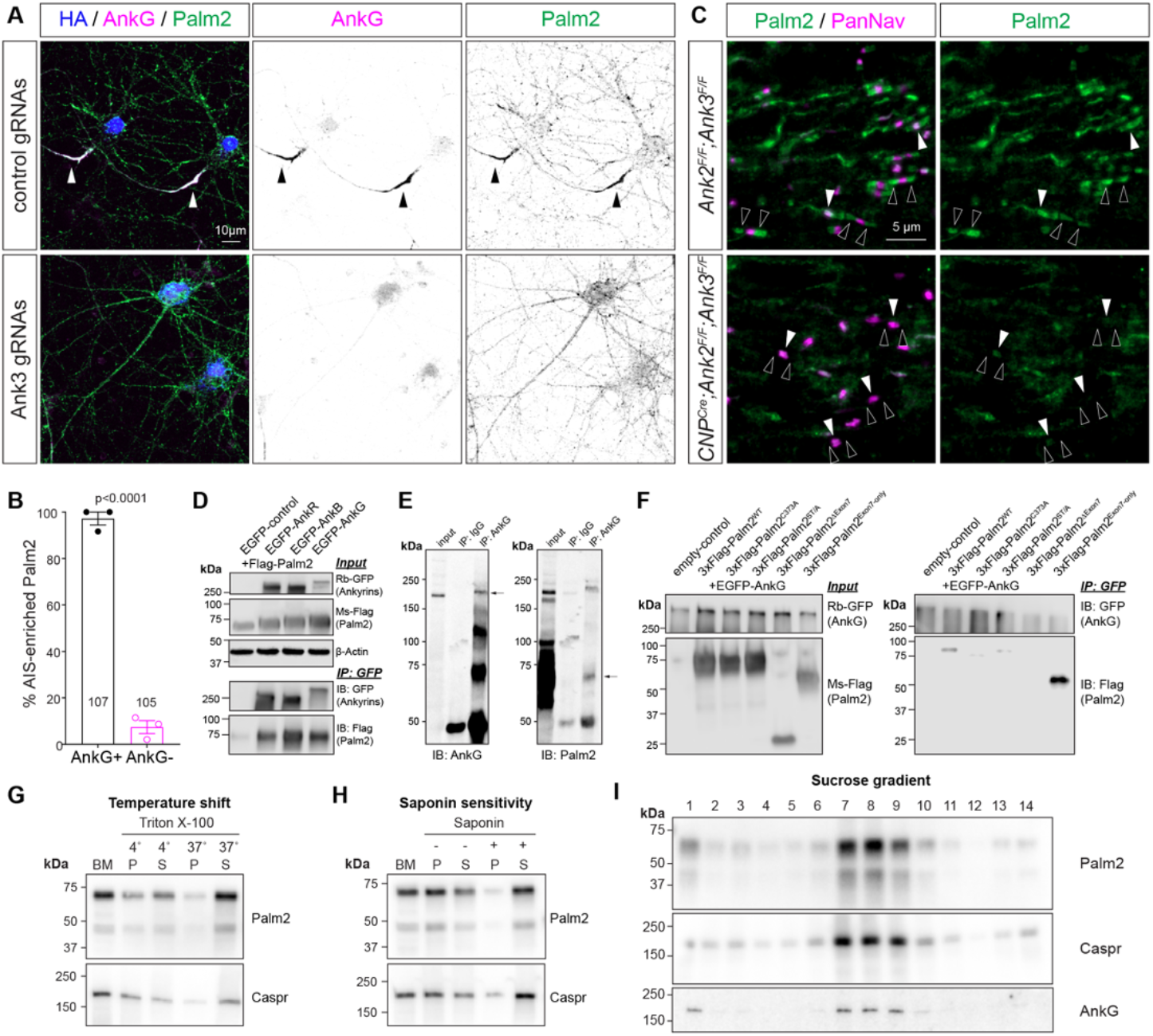
Palm2 interacts with AnkG through alternatively spliced exon 7 and is a lipid raft-associated protein. (A) Immunostaining of Palm2 (green) and AnkG (magenta) in HA-positive (blue) transduced neurons with either control or *Ank3* gRNAs. Arrowheads indicate AIS. (B) Percentage of cells with AIS-enriched Palm2 in AnkG-positive and AnkG-negative neurons. N = 3 independent cultures, with the number of neurons indicated within each bar. Error bars represent mean ± SEM. Welch’s t-test. (C) Immunostaining of Palm2 (green) and PanNav (magenta) in the optic nerves of *Ank2*^*F/F*^*;Ank3*^*F/F*^ (control) mice or *CNP*^*Cre*^*;Ank2*^*F/F*^*;Ank3*^*F/F*^ (AnkB/AnkG double conditional KO) mice at P17. White arrowheads indicate nodes of Ranvier. Black arrowheads indicate paranodes. (D) Co-immunoprecipitation of Flag-tagged Palm2 with EGFP-tagged Ankyrins (AnkR, AnkB, AnkG) in HEK293T cells. Ankyrins were immunoprecipitated with an anti-GFP antibody, followed by immunoblotting for GFP (Ankyrins) and Flag (Palm2). (E) Co-immunoprecipitation of Palm2 with AnkG from P21 mouse brain. Both AnkG and Palm2 were detected (arrows) following immunoprecipitation with an anti-AnkG antibody but not with an IgG isotype-matched control antibody. (F) Co-immunoprecipitation of Flag-tagged Palm2 mutants (C373A, ST/A, Δexon 7, or exon 7-only) with EGFP-tagged AnkG in HEK293T cells. AnkG was immunoprecipitated with an anti-GFP antibody, followed by immunoblotting for GFP (AnkG) and Flag (Palm2). (G) Immunoblotting of Palm2 and Caspr from P45 mouse brain membrane (BM) fractions treated with 1% Triton X-100 at either 4°C or 37°C. (P: pellet, S: supernatant). (H) Immunoblotting of Palm2 and Caspr from P45 mouse brain membrane (BM) fractions treated with 0.2% saponin at 4°C prior to extraction with Triton X-100. (P: pellet, S: supernatant). (I) Immunoblotting of Palm2, Caspr, and AnkG from P45 mouse brain membranes fractionated across a sucrose gradient. All three proteins cofractionate in fractions 7–9. Scale bars are shown in each panel.

### Palm2 interacts with ankyrins through its alternatively spliced exon 7

To determine if Palm2 interacts with Ankyrins we co-transfected HEK293T cells with Flag-tagged Palm2 and either GFP or GFP-tagged AnkR, AnkB, or AnkG. We found that Palm2 co-immunoprecipitates with all Ankyrins, although the affinity for AnkB and AnkG appears stronger (Fig. 4D). We also immunoprecipitated AnkG from 1 month old mouse brains and found that Palm2 could be co-immunoprecipitated by AnkG (Fig. 4E). Because deletion of exon 7 in Palm2 disrupts its AIS localization (Fig. 3G), we investigated if the corresponding protein region mediates the interaction between Palm2 and AnkG. To test this, we performed co-immunoprecipitation experiments between AnkG and truncations of Palm2. Since the C-terminal domain of Palm2 contains both prenylation and palmitoylation sites, and the molecular weight of Palm2 is larger than the predicted size, we reasoned lipid or other post-translational modifications might also regulate these interactions. Therefore, we generated point mutants lacking lipid or other post-translational modifications (PTMs). We found that whereas mutations affecting PTMs reduce the interaction between Palm2 and AnkG when co-transfected in HEK293T cells, the loss of exon 7 completely abolishes Palm2’s interaction with AnkG. Importantly, the protein region encoded by exon 7 alone is sufficient to strongly interact with AnkG in HEK293T cells (Fig. 4F). Together, these results show that AnkG and Palm2 interact through the protein domain encoded by exon 7.

### Palm2 is a lipid raft-associated protein

Since AnkG is a core component of the MPS ^26^, and Palm2 binds AnkG, we investigated whether Palm2 is also found in the MPS. We performed stochastic optical reconstruction microscopy (STORM), a form of single-molecule localization microscopy (SMLM), to examine the nanoscale organization of Palm2. Despite its interaction with AnkG, we did not find any periodicity in Palm2’s organization (Fig. S3G), and the average distance between Palm2 immunofluorescent puncta was ∼137 nm (Fig. S3H), below that of the periodic rings of actin that are spaced 190 nm apart^34^. To examine the core, insoluble AIS proteins, we extracted primary neurons for 5 min at room temperature using 0.5% Triton X-100 prior to fixation. This treatment did not remove AIS Palm2, did not result in a highly periodic organization, and only increased the peak-to-peak distance to 150 nm instead of 190 nm (Figs. S3G, H). These observations suggest that although Palm2 is associated with the cytoskeleton, it may also exist in an environment that is resistant to mild Triton X-100 treatment.

Since Palm2 is reported to be both prenylated and palmitoylated, we reasoned that Palm2 could be protected from Triton X-100 extraction by a unique lipid membrane environment. We previously showed that the major paranodal CAMs Caspr, Contactin, and NF155 have the biochemical properties of lipid raft-associated proteins ^35^.

Both AnkG and NF155 are palmitoylated ^36,37^, a reversible PTM that promotes the association and partitioning of proteins into highly ordered lipid rafts. AnkB, βII- and αII-spectrin were originally identified as paranodal proteins based on proteomics of lipid rafts isolated from myelinated axons ^38^. To determine if Palm2 is also a lipid-raft associated protein, we first showed that Palm2 is found in insoluble membrane fractions at 4°C, but after a temperature shift to 37°C, it becomes soluble in 1% Triton X-100 (Fig. 4G). Next, we found that treatment with saponin to deplete cholesterol from membranes also causes Palm2 to become soluble in 1% Triton X-100 at 4°C (Fig. 4H). Finally, we found that the 1% Triton X-100 detergent insoluble fraction containing Palm2 also floats on a sucrose density gradient in the same fractions as both Caspr and AnkG (Fig. 4I). Together, these results show that Palm2’s biochemical properties are consistent with proteins that are associated with lipid rafts, and consistent with previous studies showing AIS and paranodes and their associated proteins have properties of lipid rafts.

### Palm2 promotes cytoskeletal and membrane dynamics

To begin to determine the cellular function of Palm2, we performed gain-of-function experiments by expressing Flag-tagged Palm2 in primary cultured rat neurons, using GFP expression as a control. Flag-Palm2 expression induced prominent filopodial outgrowth and excessive membrane branching compared with control neurons (Fig. 5A). Notably, these morphological changes were abolished by deletion of exon 7, whereas expression of exon 7 alone was sufficient to recapitulate the phenotype. Quantitative analysis further confirmed that Palm2 over-expression significantly increased neurite branching and morphological complexity relative to GFP controls (Fig. 5B-C). Consistent with these qualitative observations, deletion of exon 7 eliminated this effect, whereas overexpression of exon 7 alone recapitulated the increase in morphological complexity induced by full-length Palm2 overexpression. Together, these findings identify exon 7 as both necessary and sufficient for Palm2-mediated morphological remodeling in neurons.

**Figure 5.**
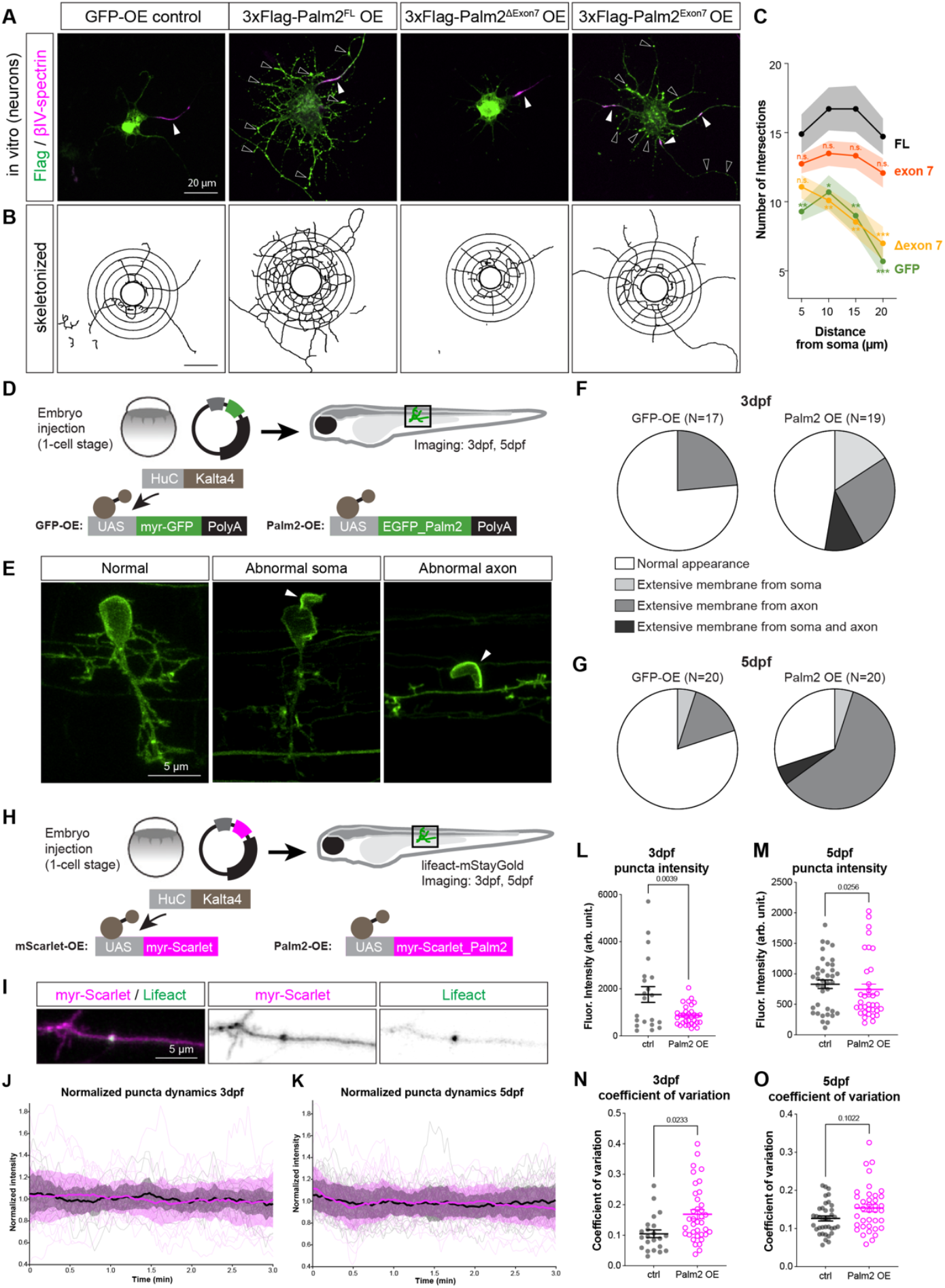
Palm2 regulates membrane and actin dynamics. (A) Immunostaining for GFP or Flag-Palm2 (green), and βIV-spectrin (magenta) in transfected neurons. White arrowheads indicate the AIS. Black arrowheads indicate protrusions and aggregates. (B) Skeletonized neurons showing intersections with concentric circles at radii of 5, 10, 15, and 25 μm from the soma to measure morphological complexity. (C) Sholl analysis showing the number of intersections at each radius in neurons expressing GFP (green), full-length Palm2-GFP (FL, black), Palm2-GFP without exon 7 (Δexon 7, yellow), or Palm2 exon 7-GFP alone (exon 7, orange). n = 10, 11, 12, and 11 transfected neurons. Two-way ANOVA with multiple comparisons against FL. *p<0.01, **p<0.001, ***p<0.0001, n.s. = not significant. (D) Illustration of overexpression of membrane-tethered myr-GFP or EGFP-Palm2 in the neurons of zebrafish larvae under the HuC:Kalta4/UAS system. (E) Representative examples of normal neurons, neurons with abnormal somatic protrusions, or neurons with abnormal axonal protrusions (arrow-heads) in zebrafish larvae. (F-G) Percentage of normal and abnormal neurons in myr-GFP-overexpressing or GFP-Palm2 expressing zebrafish at 3 days post-fertilization (dpf, F) or 5 dpf (G). (H) Illustration of co-expression of mStayGold-tagged Lifeact with membrane-tethered myr-mScarlet or mScarlet-Palm2 in zebrafish larvae. (I) Lifeact (green) in myr-mScarlet (magenta)-positive axons. (J-K) Normalized Lifeact fluorescence intensity in control cells (black) or mScarlet-Palm2 expressing neurons (magenta). The thick solid lines represent the mean, and the shaded areas indicate ± SEM. Each trace represents one punctum. (L-M) Lifeact puncta intensity in control cells (black) or mScarlet-Palm2 overexpressing cells (magenta) at 3 dpf (L) and 5 dpf (M). Each data point represents one punctum. Kolmogorov-Smirnov test. Error bars indicate mean ± SEM. For 3 dpf, N = 22 for control with 1 outlier excluded, N = 37 for Palm2 OE with 4 outliers excluded. For 5dpf, N = 38 for control, N = 39 for Palm2 OE with 4 outliers excluded. (N-O) Coefficient of variation of Lifeact intensity in control cells (black) or Palm2-overexpressing cells (magenta) at 3 dpf (N) and 5 dpf (O). Each data point represents one punctum. Kolmogorov-Smirnov test. Error bars indicate mean ± SEM. For 3 dpf, N = 22 for control, N = 37 for Palm2 OE. For 5dpf, N = 38 for control with 1 outlier excluded, N = 39 for Palm2 OE with 1 outlier excluded.

To extend these observations to neuronal morphology *in vivo*, we expressed GFP-tagged Palm2 or membrane-targeted myr-GFP (control) under the neuronal promoter *HuC*^39^ in zebrafish larvae at 3 and 5 days post fertilization (dpf) (Fig. 5D). Neurons were classified as exhibiting normal morphology, membrane protrusions from the soma, membrane protrusions from the axon, or both (Fig. 5E). Palm2 overexpression significantly increased the frequency of abnormal membrane protrusions at 3 dpf compared with controls (Fig. 5F), and these abnormalities became even more pronounced by 5 dpf (Fig. 5G). Together, these findings suggest that Palm2 overexpression drives excessive membrane growth *in vivo*.

In addition to the formation of membrane protrusions, Palm2 overexpression produced GFP-enriched puncta within discrete neuronal compartments. Because Palm2 has previously been reported to interact with Ezrin, a regulator of actin cytoskeletal dynamics^23^, we asked whether Palm2 influences actin remodeling *in vivo*. To address this, we co-expressed the filamentous actin (F-actin) reporter Lifeact-mStayGold ^40^ together with either myr-mScarlet (control) or mScarlet-Palm2 into zebrafish embryos at the one-cell stage and performed live imaging of double-positive neurons at 3 and 5 dpf (Figs. 5H-I). Lifeact accumulates at loci periodically located along axons and exhibits dynamic changes within a few minutes, indicating contant F-actin assembly and disassembly (Figs. 5I-J). We found that at 3dpf, these Lifeact loci showed reduced overall intensity in Palm2 overexpressing axons compared to controls (Fig. 5L,M). By measuring the Lifeact intensity over time, we found a significantly higher coefficient of variation in axons of Palm2-overexpressing neurons compared to controls at 3 dpf (Figs. 5L-O), indicating increased temporal fluctuation and enhanced F-actin assembly and disassembly dynamics. Together, these data suggest that elevated Palm2 expression in neurons promotes both excessive membrane protrusion and a faster turnover of actin filaments in neurons.

### Palm2 regulates AIS length

To determine the necessary roles of Palm2 in neurons, we performed CRISPR/Cas9-mediated knockout of Palm2 using 3x gRNA in primary cultured neurons from Cas9 transgenic mice. We first validated the knockout efficiency by immunostaining with independent antibodies against Palm2 (Fig. 6A). AAV-transduced neurons were identified as HA+ cells, and the AIS region was defined by AnkG staining. Both the rabbit polyclonal and mouse monoclonal Palm2 antibodies showed a significant reduction in Palm2 immunofluorescence intensity at the AIS, confirming efficient CRISPR/Cas9-mediated knockout (Fig. 6B-C). We next examined whether loss of Palm2 affects the organization of the AIS. At DIV11, Palm2 knockout neurons exhibited a modest but significant increase in AIS length, as measured by both βIV-spectrin and AnkG (Fig. 6D). Notably, despite the increase in AIS length (Fig. 6E-F), AIS formed normally and the overall organization of the AIS remained intact, with no detectable change in the mean fluorescence intensity of either AnkG or βIV-spectrin (Fig. 6G-H). These findings suggest that Palm2 is dispensable for AIS assembly but contributes to the regulation of AIS length during early development.

**Figure 6.**
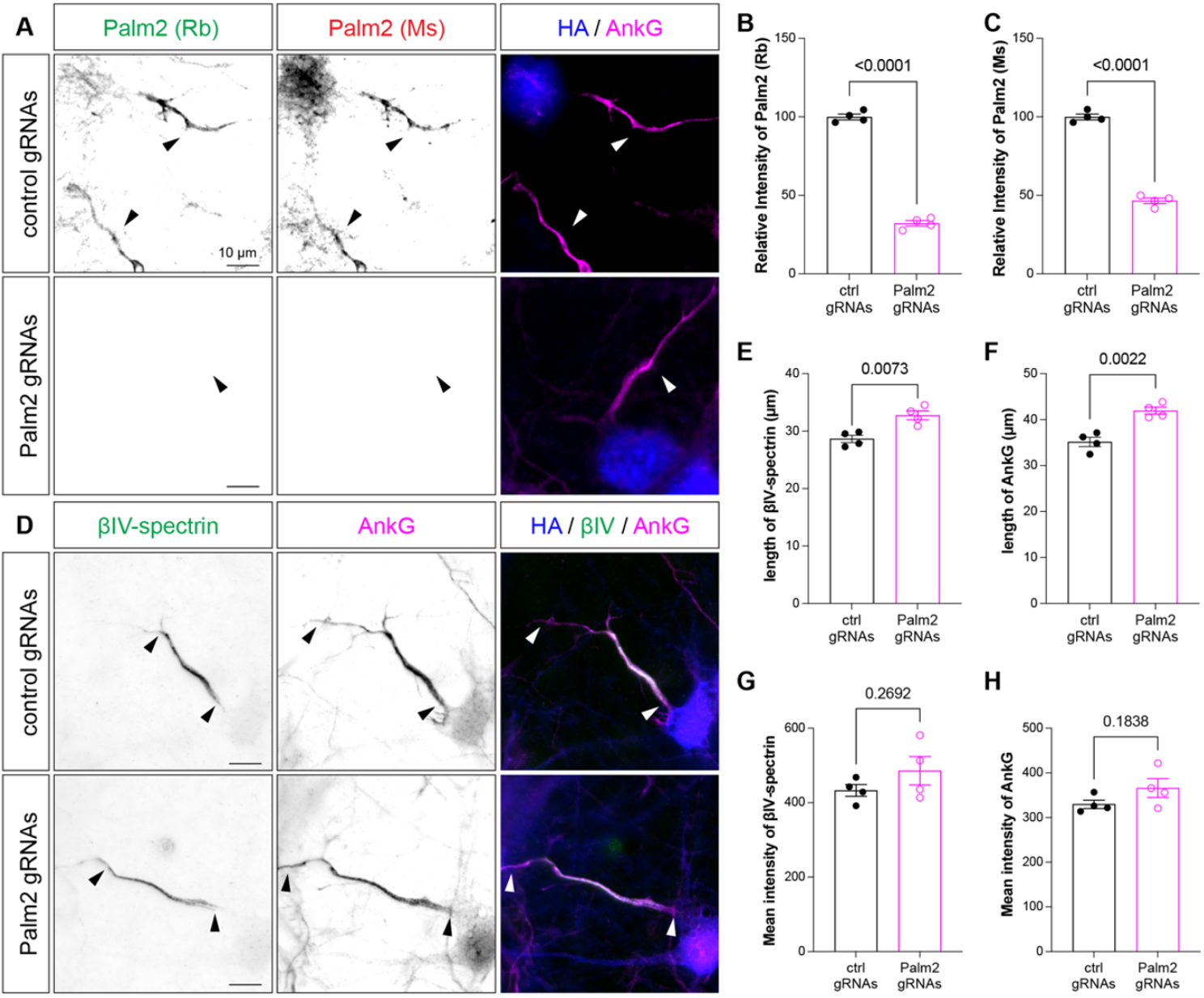
Palm2 regulates AIS length. (A) Immunostaining of Palm2 (green, rabbit polyclonal; red, mouse monoclonal) and AnkG (magenta) in HA-positive (blue) neurons transduced with control or Palm2 gRNAs. Arrowheads indicate the AIS. (B-C) Relative Palm2 immunofluorescence intensity in Cas9-expressing mouse neurons transduced with control or Palm2 gRNAs, using a rabbit polyclonal (B) and a mouse monoclonal (C) antibody against Palm2. Welch’s t-test. Error bars show mean ± SEM. Each data point is an independent culture. (D) Immunostaining for βIV-spectrin (green) and AnkG (magenta) in HA-positive (blue) neurons transduced with control or Palm2 gRNAs. Arrowheads indicate the boundaries of the AIS. (E-F) AIS length in Cas9-expressing mouse neurons transduced with control or Palm2 gRNAs, as defined by βIV-spectrin (E) or AnkG (F). Welch’s t-test. Error bars show mean ± SEM. Each data point is an independent culture. (G-H) Mean βIV-spectrin (G) and AnkG (H) fluorescence intensity in Cas9-expressing mouse neurons transduced with control or Palm2 gRNAs. Welch’s t-test. Error bars show mean ± SEM. Each data point is an independent culture.

### Loss of Palm2 has little effect on early myelination in zebrafish

Since Palm2 is also highly expressed in oligodendrocytes (Figs. S3A-C), we determined the requirement for Palm2 during early developmental myelination *in vivo*. To this end, we performed CRISPR/Cas9-mediated knockdown (KD) of Palm2 in zebrafish embryos. Cas9 protein together with five guide RNAs (gRNAs) targeting the first several exons of Palm2 was injected at the one-cell stage, and larvae were analyzed at 2, 3 and 5 dpf (Fig. S4A). Palm2 knockdown (KD) larvae were viable with very few deformed larvae (Fig. S4B), exhibited normal body length (Fig. S4C), and responded normally to external stimuli (Fig. S4D). At 3 dpf, Palm2 KD larvae showed no significant differences in the average sheath length per oligodendrocyte (Fig. S4E), sheath number per oligodendrocyte (Fig. S4F), or cumulative sheath length per oligodendrocyte (Fig. S4G). By 5 dpf, the average sheath length (Fig. S4H) and cumulative sheath length (Fig. S4J) were comparable between groups, whereas sheath number was modestly reduced (Fig. S4I). Collectively, these findings suggest that Palm2 is largely dispensable for early myelin development in zebrafish.

### Generation and validation of *Palm2*^*ΔEx4*^ mice

To investigate the function of Palm2 in mice using a constitutive genetic knockout model, we obtained *Palm2*^*ΔEx4*^ (Δex4) mice from the Knockout Mouse Project (KOMP). This allele was generated by CRISPR/Cas9-mediated deletion of exon 4 together with portions of the flanking intronic sequences, removing the 131-bp coding sequence encoding the paralemmin motif. This design also introduces a frameshift and multiple premature stop codons in exons 5, 6 and 7 (Fig. 7A). Palm2 wild type (WT), heterozygous (Δ4/+), and homozygous *Palm2*^*ΔEx4*^ (Δ4/Δ4) mice were born at the expected Mendelian ratio (Fig. S4K), indicating that loss of Palm2 does not affect viability. At P8, WT and *Palm2*^*ΔEx4*^ mice exhibited normal righting reflexes (Fig. S4L), suggesting grossly normal neurological development and general health. At two months of age, *Palm2*^*ΔEx4*^ mice also displayed normal locomotor behavior in both the open-field assay (Fig. S4M) and rotarod test (Fig. S4O), although they spent less time mobile in the center of the open field (Fig. S4N).

**Figure 7.**
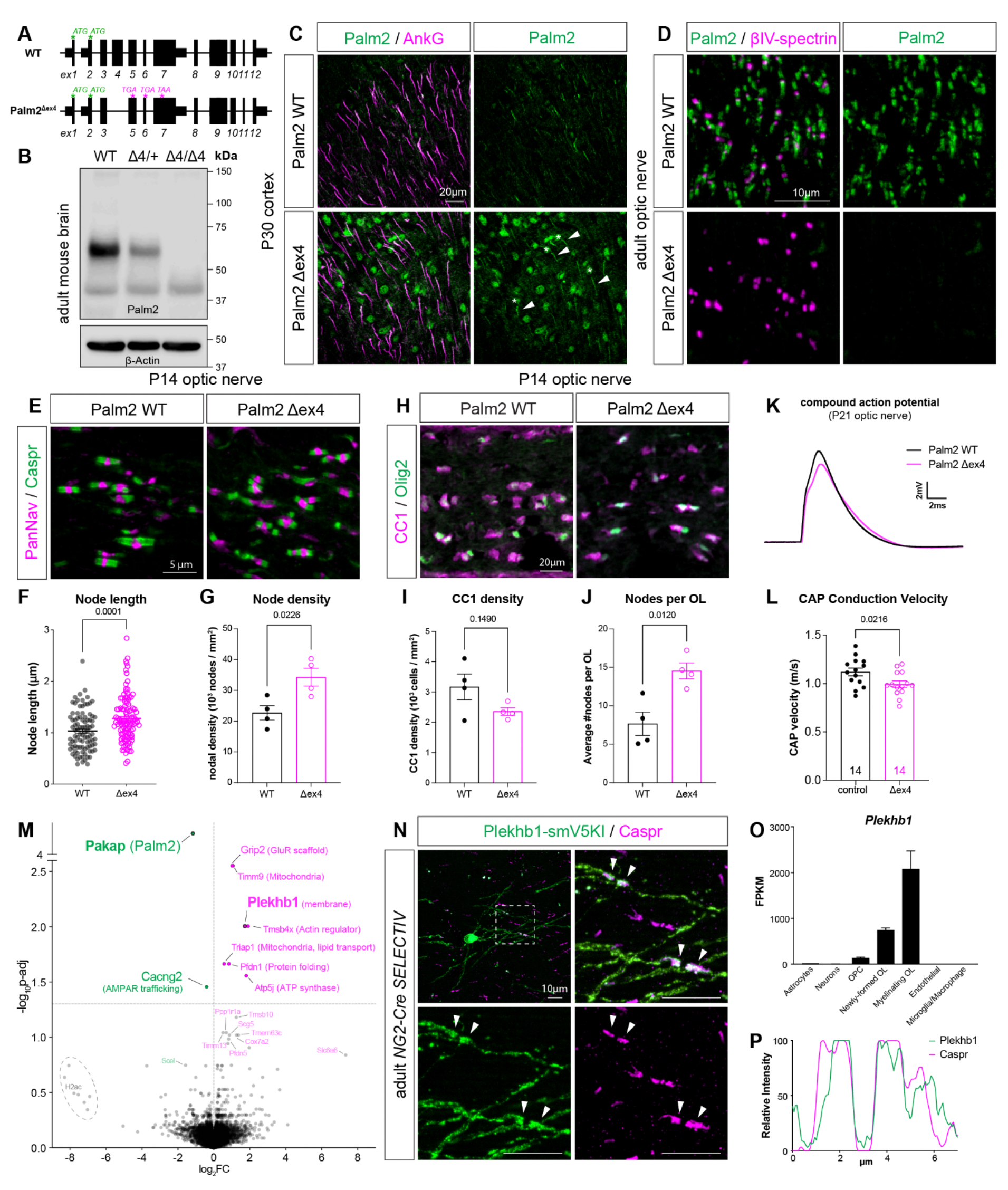
Oligodendroglial Palm2 knockout mice. (A) Schematic of the Pakap locus in WT and *Palm2*^*ΔEx4*^ mice. Exon 4 is deleted in the *Palm2*^*ΔEx4*^ mice. Start codons (ATG) are labeled in green, and premature stop codons are labeled in magenta. (B) Immunoblotting of Palm2 in adult mouse brain homogenates from Palm2 wild-type (WT), heterozygous (Δ4), and *Palm2*^*ΔEx4*^ (Δ4/Δ4) mice showing dose-dependent loss of the major Palm2-immunoreactive band ∼60 kDa. (C) Immunostaining of Palm2 (green) and AnkG (magenta) in P30 WT and *Palm2*^*ΔEx4*^ cortex, showing Palm2 immunoreactivity at the AIS (arrow-heads) in *Palm2*^*ΔEx4*^ mice. Asterisks indicate ectopic accumulation of Palm2 in the soma. (D) Immunostaining of Palm2 (green) and βIV-spectrin (magenta) in adult WT and *Palm2*^*ΔEx4*^ optic nerves, with complete loss of paranodal Palm2. (E) Immunostaining of Caspr (green) and PanNav (magenta) in P14 Palm2 WT and *Palm2*^*ΔEx4*^ optic nerves. (F) Node length defined by PanNav immunoreactivity in P14 Palm2 WT and *Palm2*^*ΔEx4*^ optic nerves. Each data point represents an individual node. N = 96 nodes in WT with 4 outliers excluded, and 109 nodes in *Palm2*^*ΔEx4*^ mice. Mann-Whitney U test. Error bars represent mean ± SEM. (G) Node density in P14 Palm2 WT and *Palm2*^*ΔEx4*^ optic nerves. Each data point represents an animal. N = 4 for both genotypes. Welch’s t-test. Error bars represent mean ± SEM. (H) Immunostaining of Olig2 (green) and CC1 (magenta) in P14 Palm2 WT and *Palm2*^*ΔEx4*^ optic nerves. (I) CC1 density in P14 Palm2 WT and *Palm2*^*ΔEx4*^ optic nerves. Each data point represents an animal. N = 4 for both genotypes. Welch’s t-test. Error bars represent mean ± SEM. (J) Node number per oligodendrocyte defined by node density divided by CC1 density in P14 Palm2 WT and *Palm2*^*Δ Ex4*^ optic nerves. Each data point represents an animal. N = 4 for both genotypes. Welch’s t-test. Error bars represent mean ± SEM. (K) Representative traces of compound action potential (CAP) recording from P21 Palm2 WT (black) and *Palm2*^*Δ*.*Ex4*^ (magenta) optic nerves. (L) CAP conduction velocity in P21 Palm2 WT and *Palm2*^*Δ*.*Ex4*^ optic nerves. Each data point represents an optic nerve. N = 14 optic nerves per genotype. Welch’s t-test. Error bars represent mean ± SEM. (M) Volcano plot showing log2 fold change (log2 FC) and -log10 adjusted p value (-log10 p-adj) for P45 brain membrane proteins in Palm2 WT and *Palm2*^*Δ*.*Ex4*^ mice detected by data-independent acquisition (DIA) mass spectrometry. N = 5 animals per genotype. The horizontal dashed line indicates the significance cutoff (p-adj < 0.05). Upregulated proteins are labeled in magenta and downregulated in green. (N) AAV/CRISPR-mediated endogenous protein tagging of Plekhb1 with smV5 (green) in adult oligodendrocytes using *NG2-Cre;SELECTIV* mice, co-labeled with Caspr (magenta). Arrowheads indicate colocalization at paranodes. (O) Cell-type specificity of Mus musculus Plekhb1 expression in the Brain-RNAseq database. (P) Representative line-scan profiles of Plekhb1 (green) and Caspr (magenta) fluorescence intensity across a pair of paranodes. Scale bars are indicated in each panel.

To confirm the loss of Palm2 protein, we performed immunoblotting using adult brain homogenates from WT, heterozygous, and *Palm2*^*ΔEx4*^ mice. The major Palm2-immunoreactive band located at ∼60-kDa was markedly reduced in HT mice and completely absent in *Palm2*^*ΔEx4*^ mice (Fig. 7B). To further confirm the loss of Palm2 at the subcellular level, we performed immunofluorescence labeling using three independent Palm2 antibodies in P30 cortex and adult optic nerve. Unexpectedly, residual Palm2 immunoreactivity was detected at the AIS in *Palm2*^*ΔEx4*^ mice, with additional ectopic accumulation in the soma (Fig. 7C). In stark contrast, paranodal Palm2 was completely absent from the *Palm2*^*ΔEx4*^ mice (Fig. 7D). Because all antibodies recognize exon 7, these results suggest that the deletion of exon 4 abolishes glial Palm2, but that neurons in *Palm2*^*ΔEx4*^ mice may process the mutant *Palm2* mRNA to produce a Palm2 variant lacking exon 4 but including exon 7.

To verify that exon 4 coding sequence was successfully deleted and that the predicted frameshift was introduced, we performed Sanger sequencing on cDNA from WT and *Palm2*^*ΔEx4*^ mouse brains. Electropherograms confirmed loss of exon 4, resulting in a +2 reading-frame shift and multiple premature termination codons (Fig. S5A). qRT-PCR further confirmed loss of exon 4-containing transcripts in *Palm2*^*ΔEx4*^ mice (Fig. S5B). However, exon 7 transcript levels remained comparable between WT and *Palm2*^*ΔEx4*^ brains (Fig. S5C), suggesting that nonsense-mediated decay is either inefficient or incomplete for the mutant transcripts in neurons.

To further characterize the transcript composition of the *Pakap* locus in the *Palm2*^*ΔEx4*^ brain, we performed nanopore sequencing of cDNA from P45 WT and *Palm2*^*ΔEx4*^ cortex. In addition to confirming deletion of exon 4 (Fig. S5D-S5E) and detecting all major annotated *Pakap* isoforms (Fig. 3B), we identified a previously unannotated transcript that was detected exclusively in *Palm2*^*ΔEx4*^ cortex (Fig. S5F). This transcript encodes a Palm2–Akap2 fusion protein containing most of exon 7, which is normally excluded from the WT *Palm2–Akap2* fusion transcript (Fig. 3B). Alignment to the mouse reference *Palm2* transcript predicted an in-frame fusion protein retaining the majority of exon 7, with only the last 14 amino acids absent (Fig. S5G). Together, these results show that the KOMP allele does not generate a complete functional null in all cell types. Instead, the *Palm2*^*ΔEx4*^ mouse is a constitutive allele that is functionally null in oligodendrocytes but hypomorphic in neurons. This unexpected allele therefore provides a unique genetic model to investigate the oligodendrocyte-specific functions of Palm2 *in vivo*.

### *Palm2*^*Δ*.*Ex4*^ mice have more nodes of Ranvier, but slower conduction velocity

To determine if loss of paranodal Palm2 affects paranode assembly in the CNS, we examined the developing optic nerve since it is one of the most homogeneous and heavily myelinated white matter tracts of the CNS. At P14, paranodes formed successfully in both WT and *Palm2*^*ΔEx4*^ mice, as indicated by Caspr immunoreactivity flanking clustered Nav channels (Fig. 7E). However, individual node length, measured using Pan-specific voltage-gated Na^+^ channel (PanNav) immunoreactivity, is approximately 20% longer in *Palm2*^*ΔEx4*^ mice compared to WT mice (1.21 μm vs. 0.97 μm; Fig. 7F). Increased node length suggests a less robust paranodal junction since the primary mechanism of Nav channel clustering in the CNS is by the paranodal axonal cytoskeleton ^41,42^. In addition, node density increased by approximately 1.5-fold in *Palm2*^*ΔEx4*^ mice (Fig. 7G) which could result from more mature oligodendrocytes. To test this, we quantified CC1-positive cells in optic nerve sections from the same animals (Fig. 7H) and observed no significant difference in the density of mature oligodendrocytes between WT and *Palm2*^*ΔEx4*^ mice (Fig. 7I). However, the average number of nodes per CC1-postive oligodendrocyte in *Palm2*^*ΔEx4*^ mice was nearly double the density found in WT mice (Fig. 7J). Alternatively, increased node density could indicate reduced individual myelin sheath length.

To determine if the longer nodes of Ranvier and higher nodal density observed in *Palm2*^*ΔEx4*^ mice have any physiological consequence, we examined compound action potentials recorded from P21 WT and *Palm2*^*ΔEx4*^ mouse optic nerves (Fig. 7K). We observed a modest but significant reduction in conduction velocity in *Palm2*^*ΔEx4*^ mice (Fig. 7L). These findings suggest that in addition to longer nodes, individual oligodendrocytes may generate a greater number of myelin sheaths during early development, resulting in shorter internodal distances, more nodes of Ranvier, and reduced conduction velocities.

### Loss of Palm2 induces proteome remodeling

To determine if compensatory mechanisms are activated after loss of Palm2, we isolated membrane proteins from whole brains of P45 *Palm2*^*ΔEx4*^ mice and performed quantitative proteomic analysis using data-independent acquisition (DIA) mass spectrometry (Figs. 7M, S5H, and Supplemental Data 2). Only two proteins were significantly downregulated: Palm2 itself and Cacng2 (Calcium Voltage-Gated Channel Auxiliary Subunit Gamma 2), also known as Stargazin, a key regulator of AMPA receptor trafficking and gating ^43^. Among the most significantly upregulated proteins was Grip2 (Glutamate Receptor Interacting Protein 2), a scaffold protein that in neurons anchors AMPA receptors at excitatory synapses ^44^. Other upregulated proteins included the cytoskeletal regulator Tmsb4x, which encodes the actin-sequestering protein thymosin β4; Pfdn1, which encodes a subunit of the prefoldin chaperone complex that facilitates folding of newly synthesized actin and tubulin; and the mitochondrial proteins Timm9, Triap1, and Atp5j.

Among the most upregulated proteins, we focused on Plekhb1 (Pleckstrin Homology Domain Containing B1), a poorly understood membrane-associated protein that is highly enriched in the oligodendrocyte lineage (Fig. 7O). Although previous transcriptomic studies identified its highly restricted cell-type-specific expression pattern, Plekhb1’s subcellular localization and function in oligodendrocytes remain unknown. To identify Plekhb1’s location in oligodendrocytes with single-cell resolution and to avoid unvalidated antibodies, we used OASIS, an AAV-mediated endogenous genome-editing platform for adult oligodendrocytes *in vivo* ^45^. This strategy tags endogenous Plekhb1 with the smV5 epitope in *NG2-Cre SELEC-TIV* mice via retro-orbital AAV delivery. Plekhb1-smV5 was specifically localized to the plasma membrane of myelinating oligodendrocytes and exhibited robust paranodal enrichment, wrapping around Caspr-positive paranodes (Figs. 7N and 7P). Together, these findings identify two downregulated and seven upregulated proteins that may underlie compensatory mechanisms following loss of paranodal Palm2. In addition, these experiments suggest Plekhb1 as a putative paranodal protein that may partially compensate for Palm2.

### Loss of Palm2 causes age-dependent deterioration of paranodes

Paranodes are highly stable structures but can undergo age-dependent disassembly in mice lacking key structural components of paranodes ^46,47^. To determine if loss of Palm2 disrupts paranode structure in aged mice, we performed immunofluorescence labeling of key paranodal proteins, including AnkG, Neurofascin, and Caspr in 16-month-old optic nerves of Palm2 WT and *Palm2*^*ΔEx4*^ mice. We found that paranodal AnkG was undetectable in aged *Palm2*^*ΔEx4*^ mice, while the nodal AnkG remained unaffected (Figs. 8A, and Fig. S6A). Quantitative analysis showed a significant decrease in the ratio of paranodal to nodal AnkG in aged *Palm2*^*ΔEx4*^ mice (Fig. 8B-C), suggesting an oligodendrocyte-specific deficit. Consequently, we also observed profound disruption of paranodal Neurofascin in the 16-month-old optic nerve of *Palm2*^*ΔEx4*^ mice (Fig. 8D, and S6B), phenocopying the previously described paranodal defects in aged *NG2*^*Cre*^*;Ank3*^*F/F*^ (AnkG cKO) mice ^46^. Quantitative analysis showed a significant increase in the frequency of abnormal paranodes in *Palm2*^*ΔEx4*^ mice compared to control mice (Fig. 8E).

**Figure 8.**
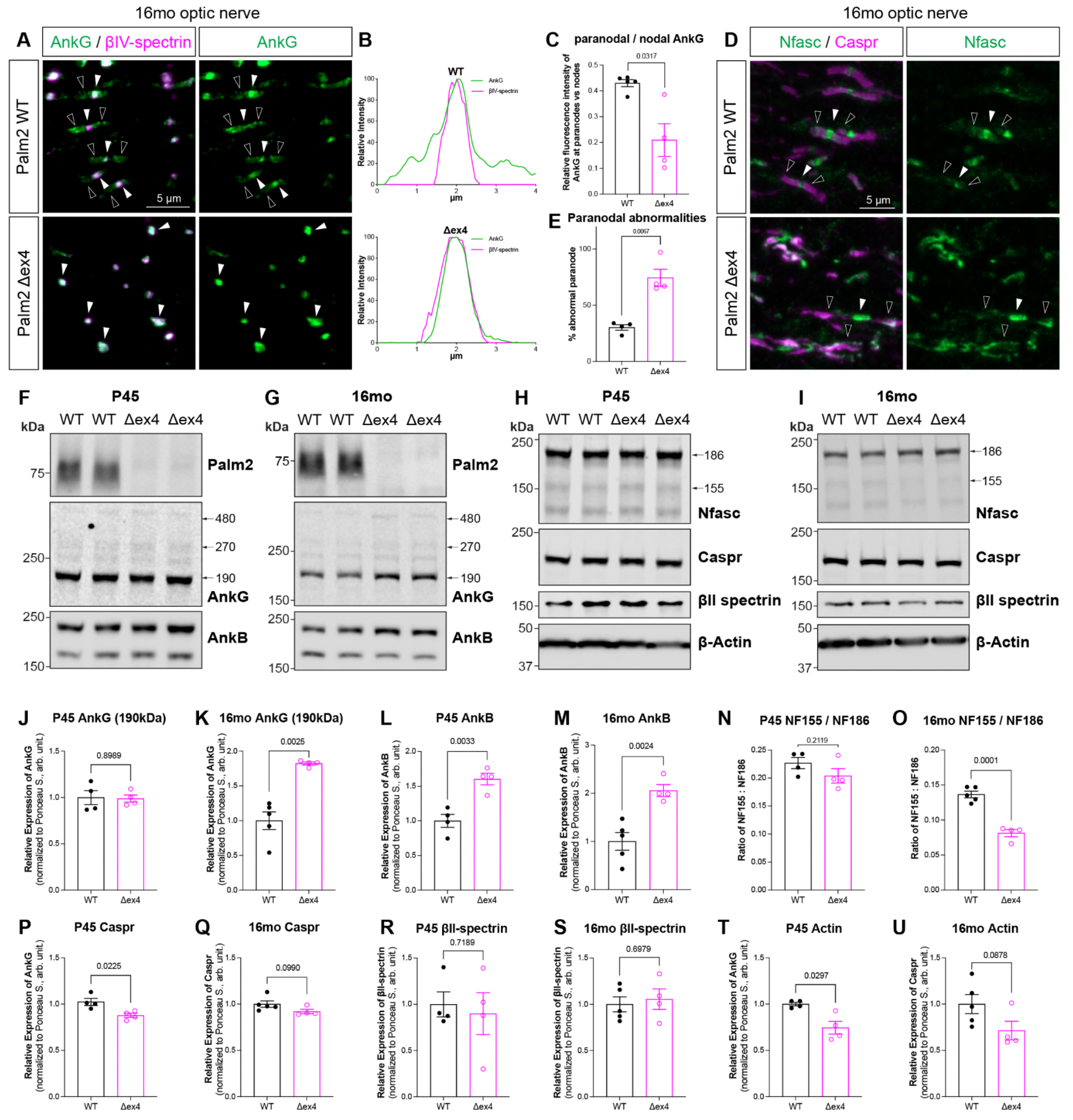
Palm2 preserves paranode integrity during aging. (A) Immunostaining of AnkG (green) and βIV-spectrin (magenta) in 16-month-old Palm2 WT and *Palm2*^*Δ*.*Ex4*^ optic nerves; paranodal AnkG immunoreactivity is lost in Palm2 *Palm2*^*Δ*.*Ex4*^ mice. White arrowheads indicate nodal AnkG; black arrowheads indicate paranodal AnkG. (B) Representative line-scan profiles of AnkG (green) and βIV-spectrin (magenta) immunofluorescence intensity across a node of Ranvier. (C) Ratio of paranodal to nodal AnkG immunofluorescence intensity in 16-month-old Palm2 WT and *Palm2*^*Δ*.*Ex4*^ optic nerves. N= 5 WT, 4 *Palm2*^*Δ*.*Ex4*^. Welch’s t-test. Error bars represent mean ± SEM. (D) Immunostaining of Neurofascin (Nfasc, green) and Caspr (magenta) in 16-month-old Palm2 WT and *Palm2*^*Δ*.*Ex4*^ optic nerves, showing disrupted paranode organization in *Palm2*^*Δ*.*Ex4*^ mice. White arrowheads indicate nodes of Ranvier; black arrowheads indicate paranodes. (E) Percentage of paranodes that appeared abnormal as defined by disrupted Nfasc/Caspr organization in 16-month-old Palm2 WT and *Palm2*^*Δ*.*Ex4*^ optic nerves. N= 4 mice for each genotype. Welch’s t-test. Error bars represent mean ± SEM. (F-I) Immunoblots of brain membrane proteins from P45 (F, H) and 16-month-old (G, I) WT and *Palm2*^*Δ*.*Ex4*^ mice. Major paranodal proteins were examined, including AnkG and AnkB (F-G), Nfasc, Caspr, βII-spectrin, and β-Actin (H-I). (J-U) Relative expression levels of paranodal proteins, including the 190-kDa AnkG isoform (J, P45; K, 16mo), AnkB (L, P45; M, 16mo), the ratio of NF155/NF186 (N, P45; O, 16mo), Caspr (P, P45; Q, 16mo), βII-spectrin (R, P45; S, 16mo), and β-Actin (T, P45; U, 16mo). N= 4 mice for each genotype. Welch’s t-test. Error bars represent mean ± SEM. Scale bars: 1 μm.

To examine whether the paranodal protein complexes display age-dependent alterations in *Palm2*^*ΔEx4*^ and WT mice, we performed immunoblotting of whole brain membrane homogenates from either P45 (Fig. 8F, H) or 16-month-old (16mo; Fig. 8G, I) mice. We examined known key components of the paranodal junctions including AnkG, AnkB, Caspr, Nfasc, and βII-spectrin (Fig. 8F-I). We found that the paranode-specific isoform of the scaffolding protein AnkG (190 kDa) was unchanged at P45 (Fig. 8J), but showed a 2-fold increase in the 16-month-old mice (Fig. 8K). In contrast, AnkB, a known paranodal protein that compensates for the loss of paranodal AnkG^7^, is consistently upregulated in both P45 (Fig. 8L) and 16-month-old brains (Fig. 8M). The dramatic increase in total Ankyrins accompanied by complete loss of the paranodal immunoreactivity of AnkG in *Palm2*^*ΔEx4*^ mice suggests that Palm2 may regulate the proper localization and maintenance of AnkG at the paranode rather than affecting its overall protein synthesis or turnover within the cell.

Consistently, the ratio between paranodal and nodal Neurofascin (NF155:NF186) also showed an age-dependent decrease of around 2-fold at 16 months of age (Fig. 8N-O), while the total amount of Caspr remained unchanged in 16-month-old *Palm2*^*ΔEx4*^ mice (Fig. 8Q) despite a slight decrease at P45 (Fig. 8P), suggesting an oligodendrocyte-specific response to the loss of Palm2 during aging. Interestingly, although the cytoskeletal protein βII-spectrin remained unchanged (Fig. 8R-S), the amount of actin was reduced in P45 (Fig. 8T) but not aged *Palm2*^*ΔEx4*^ mice (Fig. 8U). Together, these findings suggest that Palm2 is required for the long-term retention of AnkG at paranodes, rather than AnkG’s initial recruitment, and that failure to retain AnkG at paranodes eventually results in reduced paranodal NF155 and derangement of paranodal junctions.

## Discussion

Axons and oligodendrocytes have distinct, specialized membrane domains that play essential roles in action potential initiation and propagation. These domains share a common molecular organization centered on the master scaffolding protein AnkyrinG ^48^. However, the composition of this core protein complex remains incompletely understood. Here, we reported a genetic mouse model that permits the unbiased identification of endogenous AnkG proximity proteomes. We used this mouse to reveal Paralemmin 2 (Palm2) as a previously unrecognized component of the AIS, nodes of Ranvier, and paranodes. A detailed analysis of Palm2 expression shows that unlike other AIS and node-enriched proteins, Palm2 is found only transiently at these axonal domains in developing neurons; in oligodendrocytes Palm2 is found at paranodes throughout life. This cell type- and developmental stage-specific localization is regulated by alternative splicing of Palm2’s terminal coding exon, which is both necessary and sufficient for its AnkG interaction. Furthermore, this alternatively spliced exon confers lipid raft association through palmitoylation and prenylation, potentially coupling membrane lipid organization to cytoskeletal dynamics. Although Palm2 is largely dispensable for the initial assembly of these membrane domains, it is essential to maintain paranodes during aging. Thus, Palm2 is an alternatively spliced, AnkG-interacting organizer of polarized membrane domains that links maintenance of membrane lipid organization to cytoskeletal architecture in both neurons and oligodendrocytes.

An important observation here is the identification of altenative splicing as a recurring mechanism to diversify polarized membrane domains in axons and oligodendrocytes. Previous studies show that neurons use the larger isoforms of AnkG and Neurofascin to organize AIS and nodes of Ranvier, while oligodendrocytes use smaller isoforms at paranodes ^6,7,49,50^. Palm2 is a third example of this intriguing phenomenon. Such diversification through splicing may be an efficient evolutionary strategy to selectively regulate membrane targeting and protein-protein interactions in different cell types using a minimal set of genes. Whether similar mechanisms extend to additional components of polarized membrane domains and how these splicing decisions are regulated remain important questions for future investigation. Furthermore, the unexpected phenomenon we observed where neurons and oligodendrocytes responded differently to the same genetic disruption of the Palm2 locus emphasizes differences in gene regulation between these cell types. While oligodendroglial Palm2 was efficiently eliminated from paranodes in *Palm2*^*ΔEx4*^ mice, neuronal Palm2 expression persisted at the AIS. Our discovery of an unrecognized mutant transcript in the *Palm2*^*ΔEx4*^ cortex, which was absent from WT mice, suggests that neurons may circumvent disruptions through alternative transcript processing and be more tolerant to deviations from normal gene regulatory programs. Consistent with this notion, previous studies reported that both alternative splicing and translational readthrough events occur more frequently in neurons than oligodendrocytes ^33,51^, suggesting that neuronal gene regulation exhibits greater flexibility in response to genetic perturbations.

In addition to its AnkG-interacting properties, Palm2 is also a lipid raft-associated protein due to both dual palmitoylation at cysteines found in the critical AnkG-binding domain encoded by exon 7, and prenylation at its C-terminal CAAX motif; these three lipid anchors cause Palm2 to strongly associate with the cell membrane. In comparison, Neurofascin is singly palmitoylated ^36^. We previously reported that NF155 partitions into lipid rafts at paranodes in oligodendrocytes, but that in paranodal mutant mice the association of NF155 with lipid rafts is disrupted ^35^. AnkG is also singly palmitoylated, and this palmitoylation was previously shown to be essential for its localization and function at the AIS ^52^. The multiple lipid modifications in Palm2 suggest it may be even more strongly associated with rafts than Neurofascin or AnkG. In contrast to paranodes, mature AIS *in vivo* do not need Palm2 for their maintenance. Nevertheless, the mild effect on AIS in primary cultured neurons after Palm2 knockout suggests that Palm2 may participate in the initial maturation of the AIS. We speculate that once the AIS is fully assembled with a robust cytoskeleton, clustered CAMs and ion channels, and a fully assembled AIS extracellular matrix, the AIS is less dependent on its lipid membrane for its structural and molecular maintenance. In contrast, we suggest that protein-protein interactions alone are unlikely to account for the remarkable stability of the paranodal junction throughout life. A highly specialized paranodal lipid raft may function not only as a signaling platform, but also provide an additional layer of structural stability by enriching paranodal proteins with lipid modifications that preferentially partition into paranodal lipid rafts. We suggest that Palm2 is a key regulator of paranodal lipid raft assembly and its loss results in destabilization of the paranodal junction with increasing age.

The relatively mild developmental phenotypes together with the pronounced age-dependent disruption of the paranodes in *Palm2*^*ΔEx4*^ mice therefore suggest that Palm2 functions primarily to stabilize rather than instruct the assembly of the paranodal junction. Developmental assembly and lifelong maintenance are often studied in parallel, yet these processes likely rely on only partially overlapping mechanisms. While developing membrane domains are highly dynamic and may tolerate substantial perturbations by activating redundant pathways and developmental plasticity, a mature membrane domain must preserve its organization for decades or throughout life, despite continuous protein turnover and membrane remodeling, as well as chronic or acute mechanical damage and stress. Consistent with this notion, we identified proteomic remodeling in young *Palm2*^*ΔEx4*^ mice, especially upregulation of several proteins involved in membrane organization or cytoskeletal regulation, including paranode-enriched Plekhb1. Thus, activation of compensatory pathways may be sufficient to preserve nervous system function during development but become insufficient during aging. This interpretation is also consistent with an age-dependent effect of the paranode instability in oligodendrocyte-specific AnkG KO mice ^46^. More broadly, these findings highlight distinctions between mechanisms governing developmental assembly and those required for long-term maintenance of membrane domains.

In aged *Palm2*^*ΔEx4*^ mice we observed a seemingly contradictory phenomenon where the total AnkG protein increased by almost two-fold despite it being completely absent from the paranodes. These results indicate that Palm2 is responsible primarily for AnkG’s proper localization and maintenance at paranodes rather than its total expression levels. We speculate that Palm2-associated lipid rafts stabilize AnkG by promoting its partitioning into paranodal lipid rafts, and that loss of Palm2 permits the diffusion of AnkG away from paranodes. The increase in total AnkG protein abundance may represent a failed compensatory attempt by oligodendrocytes to restore proper paranode scaffolding. The excessive accumulation of oligodendroglial AnkG in membrane homogenates from aged *Palm2*^*ΔEx4*^ mice suggests that mislocalized AnkG-190 may be passively incorporated into myelin membranes. More broadly, our findings highlight that the abundance and localization of a protein can be uncoupled in highly polarized cells like neurons and glia. Thus, quantitative proteomic or immunoblot analyses alone may underestimate or misinterpret the functional outcome when the primary consequence is disrupted protein localization rather than protein expression. Future advanced proteomic techniques with spatial and temporal resolution may further accelerate real biological discoveries.

Finally, our experiments demonstrate the power of endogenous proximity proteomics as an unbiased strategy for molecular discovery through interrogation of endogenous protein interactomes *in vivo*. Despite the functional significance of the paranodal junction, Palm2 is the first paranodal protein to be newly identified and functionally characterized in nearly two decades. The first molecular description of paranodes and axoglial interactions included the landmark discoveries of Caspr, Contactin, and Neurofascin ^49,53-56^. Later, several paranodal cytoskeletal proteins were also identified ^38,57^. These CAMs and cytoskeletal proteins are now recognized as necessary components of paranodes, and more recent studies revealed human pathogenic variants and autoimmune diseases targeting the paranodal CAMs ^58,59^. However, the pace of discovery stalled, creating the impression that the molecular composition of the largest intercellular adhesive junction in the vertebrate nervous system had been largely defined. Instead, this apparent slowdown primarily reflects the technical challenges associated with conventional biochemical purification, as both the detergent insolubility of the paranode and extensive cytoskeletal interactions hinder traditional proteomic analyses. TurboID-based proximity proteomics overcomes these challenges, and our findings demonstrate that substantial new biology remains to be discovered at paranodes and other AnkG-enriched membrane domains. Beyond Palm2, the AnkG-TurboID proximity proteomes uncovered numerous candidate proteins with unknown functions at AIS, nodes, paranodes, and the PJZ. The proximity proteomes reported here are a rich resource and roadmap for future systematic characterization and functional interrogation of candidates. These studies may reveal new principles governing the molecular organization of specialized AnkG-enriched membrane domains in neurons, oligodendrocytes, and PJZs. The *Ank3*^*LSL-TurboID*^ mouse should also be combined with other Cre-lines to define additional AnkG-enriched membrane domains and protein complexes in diverse tissues and cells including heart and epithelial cells, respectively ^60,61^. More broadly, endogenous proximity biotinylation provides a versatile platform that can be readily extended to other subcellular domains, enabling unbiased identification of previously unrecognized mechanisms for functional organization of the nervous system. Importantly, the AAV/CRISPR-mediated endogenous genome editing approach in both neurons and oligodendrocytes that we used here, also provides a rapid orthogonal validation platform independent of protein specific antibodies, ensuring reproducibility while accelerating mechanistic studies of nervous system development, maintenance, and function in health and disease.

Together, the results presented here not only describe new biology important to understand the assembly and maintenance of neuronal and oligodendroglial membrane domains, but also illustrate a framework for discovering previously unrecognized components of polarized membrane domains – from discovery, to validation, to investigation of function. Conceptually, the results illustrate how shared molecular toolkits can be diversified through alternative splicing to generate unique neuronal and glial membrane domains.

## ACKNOWLEDGEMENTS

This work was supported in part by the National Institutes of Health / National Eye Institute (K99EY037780 to X.D.), the National Multiple Sclerosis Society (NMSS-TA-2403-42997 to X.D.), the National Institutes of Health / National Institute of Mental Health (R01MH121544 to M.N.R.), the National Institutes of Health / National Institute of Neurological Disorders and Stroke (R35NS122073 to M.N.R. and F31NS139435 to V.L.P.), the National Institutes of Health / National Institute of Arthritis and Musculoskeletal and Skin Diseases (R01AR074988 to M.N.R.), the Adelson Medical Research Foundation (to A.L.B. and M.N.R.), the McNair Medical Institute at The Robert and Janice McNair Foundation (to Y.G.), and Baylor College of Medicine seed funding (to Y.G.). This project was in part supported by the Genomic and RNA Profiling Core at Baylor College of Medicine with funding from the CPRIT (RP250580). The Baylor College of Medicine Mass Spectrometry Proteomics Core is supported by the Dan L. Duncan Comprehensive Cancer Center Award (P30CA125123), CPRIT Core Facility Award (RP210227), Intellectual Developmental Disabilities Research Center Award (P50HD103555). We acknowledge the joint participation of the Diana Helis Medical Research Foundation and the Adrienne Helis Malvin Medical Research Foundation and Baylor College of Medicine in support of this research (Bruker timsTOF Ultra2). We thank Chris and Trudy Cunningham and the Cunningham family for support. We thank Drs. Alexander J King and Garret Anderson (UC Riverside) for consulting and providing metadata for exploratory splicing analysis; We also thank all members of the Rasband laboratory for helpful discussions.

## AUTHOR CONTRIBUTIONS

Conceptualization, X.D. and M.N.R.; Investigation, X.D., Y.O., V.L.P., W.Z., A.P.A., L.G.–L., S.G.H., Y.X., D.V.M.N., J.R.C., D.J.C., A.J., J.A.O.–P., T.T.T., and J.L.; data curation, Z.L., Z.Y., J.A.O.–P., andA.B.S.; formal analysis, X.D., Y.O., W.Z., A.P.A., L.G.–L., Z.L., Z.Y., Y.X., J.R.C., Y.W., A.B.S., and J.L.; visualization, X.D., Y.O., V.L.P., W.Z., Z.L., Y.X., and J.L.; resources, M.T.; writing – original draft, X.D. and M.N.R.; writing – review & editing, X.D. and M.N.R.; funding acquisition, X.D., V.L.P., Y.G., A.L.B., and M.N.R.; supervision, D.C.K., A.L.B., A.M., L.S., P.L., J.L., Y.G., and M.N.R.

## Declaration of generative ai and ai-assisted technologies in the writing process

During the preparation of this work, the authors used ChatGPT to check for grammar errors. After using this tool or service, the authors reviewed and edited the content as needed and take full responsibility for the content of the publication.

## SUPPLEMENTAL INFORMATION

Figures S1–S6

## MATERIALS AND METHODS

### Generation of transgenic mice

Mice were group housed at Baylor College of Medicine under a 12 h/12 h light/dark cycle at 20–22°C and 40–60% humidity, with ad libitum access to food and water.

*Ank3*^*LSL-TurboID*^ knock-in (KI) mice were generated by TransViragen; *Palm2*^*Δ*.*Ex4*^ mice (RRID:MMRRC_042398-JAX), Cas9 knock-in mice (RRID:IMSR_JAX:027650), SELECTIV mice (RRID:IMSR_JAX:037553), and Vglut2^Cre^ mice (RRID:IMSR_EM:04587) were obtained from The Jackson Laboratory; NG2^Cre^ mice (RRID:IMSR_JAX:008533) were generously provided by Dr. Jonah Chan (University of California, San Francisco), and Myo^Cre^ mice were generously provided by Dr. Eric Olson (UT Southwestern Medical Center). Both male and female mice were used for experiments. Only female NG2^Cre^ mice and male Myo^Cre^ mice were used for breeding.

#### Ank3^LSL-TurboID^knock-in

Ank3LSL-TurboID knock-in (KI) mice were generated by TransViragen through targeted insertion of a loxP-STOP-loxP cassette with a duplicated Ank3 exon 42 and TurboID coding sequence into the endogenous Ank3 locus. Specifically, two guide RNAs targeting the intron 41 and intron 42 were selected for genome editing: Ank3-5g93T: 5’-ATGGGGATCGTTAACGGTT-3’ Ank3-3g76B: 5’-GCCGTAGGTACCTAAAATG-3’

The donor vector was designed to produce an Ank3 allele with 3’UTR from exon 43, producing the wild-type AnkG protein in the absence of Cre recombinase. In the presence of Cre, the exon 42-43 fusion and SV40pA stop cassettes were removed, allowing splicing to the exon 42-TurboID fusion then to the native exon 43, producing a TurboID-tagged AnkG protein.

The donor vector plasmid was cloned with the following key elements:

An 1000 bp 5’ homology arm corresponding to intron 41 sequences immediately before the 5’ gRNA cut site, a loxP site inserted at the 5’ gRNA cut site, a 327 bp 3’ end of intron 41 including native splice acceptor and 3693 bp fusion of exons 42 and 43, 3x SV40pA stop cassette, FRT and loxP, duplicate copy of 327 bp 3’ end of intron 41 and exon 42 with 3xHA-tagged TurboID coding sequence inserted upstream of the native stop codon, 844 bp intron 42 5’ sequence followed by a C to A mutation that destroys the PAM site for the Ank3-3g76B guide RNA, and an 1000 bp 3’ homology arm corresponding to the 1000 bp downstream of the PAM mutation (Fig. 1A).

#### Palm2^*ΔEx4*^ *mice*

*Palm2*^***ΔEx4***^ mice were obtained from KOMP and validated by Sanger sequencing (C57BL/6NJ-Pakapem1(IMPC)J/Mmjax, RRID:MMRRC_042398-JAX). Specifically, four guide RNAs targeting the genomic region surrounding exon 4 were selected for genome editing:

5’-CATGGCTTTCGGTATCTGCT-3’

5’-AGACAACAGATGACACCTTT-3’

5’-CGTGCTGCGACTTTTCCCTG-3’

5’-TCCTAAATAGATACTGGTGT-3’

The resulting allele contains a 292-bp deletion encompassing the 131-bp exon 4 and portions of the flanking introns, leading to a frameshift after amino acid 42 and a premature stop codon 10 amino acids later.

#### Genotyping

Genotyping for mice was performed using standard PCR with the following primers:

Ank3^LSL-TurboID^ -KI-F: 5’-GCTCCAATCTGTAGAAGTT-GGTGC-3’

Ank3^LSL-TurboID^ -KI-R: 5’-CACAGATGACCTTGGCTTT-GAC-3’

Cas9-WT-F: 5’-AGTGGGACTGCTTTTTCCAG-3’

Cas9-WT-R: 5’-GATCTGGGGCCATAAATGC-3’

Cas9-mut-F: 5’-GGGCAACGTGCTGGTTATTG-3’

Cas9-mut-R: 5’-CCAGGCCGATGCTGTACTTC-3’

Cre-F: 5’-TGCTGTTTCACTGGTTATGCGG-3’

Cre-R: 5’-TTGCCCCTGTTTCACTATCCAG-3’

*Palm2*^*ΔEx4*^ -F: 5’-GACAGATTTGATGGAGAAGTCC-3’

*Palm2*^*ΔEx4*^ -R: 5’-ACTTACAGGAGAGCATTTGGG-3’

SELECTIV-mut-F: 5’-GGCAAACACCTTTGAAGTCC-3’

SELECTIV-mut-R: 5’-ACCTTCAGCTTGGCGGTCT-3’

SELECTIV-ctrl-F: 5’-AGTGGGACTGCTTTTTCCAG-3’

SELECTIV-ctrl-R: 5’-GATCTGGGGCCATAAATGC-3’

### Transient gene expression in Zebrafish

Fertilized zebrafish embryos were microinjected at the one-cell stage with approximately 1 nL of injection solution containing 20 ng/µL plasmid DNA, 25 ng/µL Tol2 transposase mRNA, 0.02% phenol red, and 0.2 M KCl. Injected F0 embryos were either used for transient-expression experiments or raised to adulthood.

### CRISPR/Cas9-mediated disruption of zebrafish palm2

Single-guide RNAs (sgRNAs) targeting zebrafish *palm2* were designed using CHOPCHOP and CRISPOR. DNA templates containing a T7 promoter and the sgRNA sequence were generated by PCR, and sgRNAs were synthesized by *in vitro* transcription using the MEGAshortscript T7 Transcription Kit. Following transcription, sgRNAs were purified and combined with recombinant Cas9 protein to generate ribonucleoprotein complexes.

Approximately 1 nl of the sgRNA-Cas9 mixture was injected into one-cell-stage embryos. A non-targeting sgRNA was injected with Cas9 protein as a control. sgRNA totals at 100ng/ul and Cas9 at 1mg/ul. The following sgRNA sequences were used:

Control sgRNA: 5’-GCGAGGTATTCGGCTCCGCG-3’

*palm2* exon 3 sgRNA 1: 5’-AGACGAAGACCGCAG-GAAAC-3’

*palm2* exon 4 sgRNA 2: 5’-GGTTCTCCGA-GAGCGACTGC-3’

*palm2* exon 6 sgRNA 3: 5’-GGTGACGGGTTT-GTCAGCGG-3’

*palm2* exon 6 sgRNA 4: 5’-GAAATCTCTGCGGGA-GAAGA-3’

*palm2* exon 6 sgRNA 5: 5’-GTCTGCTGCATT-GCCGTCGA-3’

Injected F0 larvae were used for morphological, behavioral, and myelin analyses at the indicated developmental stages.

### Mounting and live imaging of zebrafish larvae

For live imaging, zebrafish larvae were anesthetized with 0.16 mg/mL tricaine, equivalent to approximately 600 µM, in embryo medium. Larvae were mounted laterally in 0.8% low-melting-point agarose (Sigma-Aldrich, A9414) on a coverslip.

Live imaging was performed using an upright Zeiss LSM 990 confocal microscope equipped with Airyscan 2 and operated in 4Y fast-acquisition mode. A ×20/1.0-NA water-immersion objective was used. EGFP and mStayGold were excited with the 488-nm laser, and mScarlet was excited with the 561-nm laser. Imaging parameters were held constant among experimental groups within each experiment.

### Primary neuronal culture

For primary rat neurons, cerebral cortices were dissected in ice-cold HBSS, minced, and digested in 0.25% trypsin supplemented with DNase I (1 mg/mL) for 20 min at 37°C. Dissociated neurons were plated onto poly-D-lysine (1 mg/mL)- and laminin (10 μg/mL)-coated glass coverslips at a density of 2.2 × 10^4^ cells/cm^2^ for immunostaining experiments. Neurons were maintained in Neurobasal medium supplemented with 2% B27, 1% GlutaMAX, and 1% penicillin-streptomycin (P/S). Three hours after plating, the medium was replaced with fresh culture medium. One-third of the culture medium was replaced every 6 days. Neurons were fixed or harvested at the indicated days *in vitro* (DIV).

For primary mouse neurons, Briefly, neonatal Cas9 KI mice were euthanized by decapitation, and forebrain tissues containing both cortices and hippocampi were rapidly isolated and dissociated with papain (Worthington Cat# LS003120). Cells were seeded at a density of ∼2.5 × 10^5^ cells/cm^2^ on poly-L-lysine (Sigma # P2636) coated coverslips and maintained in the Neurobasal Plus culture system (ThermoFisher Cat# A3653401); for mouse cultures we used gentamycin at 12 ug/mL instead of 1% P/S as was used in primary rat neurons.

For expression analysis during development, primary rat neurons were fixed at DIV4, DIV7, and DIV21 in either 4% PFA or Glyoxal + MeOH. For transient overexpression using pcDNA-CAG-Flag-Palm2, primary rat cortical neurons were transfected with 0.8 μg of plasmids using Lipofectamine 2000 in Neurobasal media at DIV 7. Media was changed after 3 hours, and the cells were harvested for immunofluorescence labeling and imaging at DIV 9.

For AAV/CRISPR-mediated endogenous KI, primary rat neurons were transduced with viral vectors at DIV 0, and media was changed after 2 days. Neurons were fixed at DIV 14 for immunofluorescence labeling and imaging.

For AAV-CamKII-V5-GOI mediated recombinant expression, neurons were transduced with viral vector at DIV 10, and media was changed after 2 days. Neurons were fixed at DIV 14 for immunofluorescence labeling and imaging.

For AAV/CRISPR-mediated KO, primary mouse Cas9 neurons were transduced with pAAV-S3G-HA at DIV1, and media was not changed. Neurons were fixed at DIV 11 for immunofluorescence labeling and imaging.

### AAV production

#### For in vitro use

HEK293T cells were seeded at the density of 3 × 10^5^ cells/well in a 12-well plate and transfected with 0.36 µg PHP.eB, 0.49 µg pHelper, and 0.22 μg of pAAV-KI vector or 0.27 μg of pAAV-KO or pAAV-OE vector using PEI MAX in Opti-MEM. Media was changed 12 hours after transfection. 48 h later, AAV-containing supernatant was collected and filtered through a Costar Spin-X centrifuge tube filter (0.45 µm cellulose acetate, Corning Cat# 8162). The AAV-containing media was directly applied to the culture.

#### For in vivo use

HEK293T cells were seeded at the density of 1.2 × 10^7^ cells/dish in two 15 cm^2^ dishes and transfected with 14 µg PHP.eB, 19 µg pHelper, and 8.6 μg of pAAV-KI vector or 11 μg of pAAV-KO or pAAV-OE vector per dish using PEI MAX in Opti-MEM. Media was changed 12 hours after transfection. Both AAV-containing media and AAV-producing cells were collected 72 h later to extract and purify AAVs. AAVs in the supernatant were mixed with 25% volume of 40% PEG8000 and via end-to-end rotation at 4°C overnight, followed by centrifugation at 5000 × g at 4°C for 10 min. AAVs in the cells were released by lysing the cells with 6 mL Citrate buffer incubation at 37°C for 5 min, vortexed and centrifuged at 3000 × g at 4°C for 15 min. Supernatant was collected and neutralized with 600 μL 2M Tris-HCl (pH 9.5) and stored at 4°C until AAVs collected from supernatants are ready for processing. AAVs from both supernatant and cells are combined and filtered through a Millex PVDF syringe filter (0.45 μm, Millipore Cat# SLHVR33RS), followed by four washes through an Amicon ultrafiltration system (100 kDa MWCO, Sigma Cat# UFC910024) by centrifugation at 5000 × g at 4°C for 40 min each until reaching a final volume of ∼100μl. AAV titration was performed using the Takara AAVPro kit following manufacture’s direction. The titer was between 10^12^ ∼10^13^ vg/mL.

To systemically deliver AAVs in adult *NG2*^*Cre*^ *SELECTIV* mice, AAVs were diluted in dPBS for a final total volume of 100 μL per mouse via retro-orbital injection.

### Plasmid construction

For AAV-CRISPR/Cas9-mediated endogenous protein tagging using NHEJ (Fig. 1K), the gene-of-interest (GOI) specific gRNA was cloned into the pAAV-backbone under a U6 promoter by ligation at the BbsI site, followed by a U6 promoter driving a gRNA targeting two *“donor recognition sites”* (DRS) flanking the spaghetti monster-V5 (smFP-V5) tag sequence, which would be released and inserted into the 5’ of the GOI. The GOI-specific gRNAs were designed using CRISPOR as previously described. Specifically, the following gRNAs were used:

rPalm2-smV5-N-KI-gRNA-1: 5’-CAATTCCGCCTCTGCCATCC-3’

rPalm2-smV5-N-KI-gRNA-2: 5’-CCGTGTGCCCTTCTCCAGGA-3’

rPalm2-smV5-N-KI-gRNA-3: 5’-GGCAGAGGCGGAATT-GCACA-3’

rPalm2-smV5-C-KI-gRNA-1: 5’-GGTAC-CTGGGGCCGGGTCTG-3’

rPalm2-smV5-C-KI-gRNA-2: 5’-AGA-GAGCCTGGCCACAGACC-3’

For AAV-CRISPR/Cas9-mediated KO of GOI (Fig. 2D), three independent gRNAs targeting different exon regions of the same GOI were cloned into the pAAV-KO vector under three U6 promoters as previously described. A smFP-HA under an hSyn1 promoter was included to facilitate identification of the transduced cells. The gRNAs were also designed by CRISPOR. Specifically, the following gRNAs were used:

control-gRNA-1: 5’-CTGTCTTCATCATGGCCGAC-3’

control-gRNA-2: 5’-GTTCGCATTATCCGAACCAT-3’

control-gRNA-3: 5’-TAAGCGTCGCAAGAAGA-3’

mPalm2-gRNA-1: 5’-GACCCCCTCTGGGCCAATGG-3’

mPalm2-gRNA-2: 5’-CGGTACCAAAGTAGTGTATG-3’

mPalm2-gRNA-3: 5’-TATGGGCTACCAAAACATCG-3’

rAnk3-gRNA-1: 5′-CTGCTCGAGAACGACACGAA-3′

rAnk3-gRNA-2: 5′-CGCTCGGTTTAACAGCAACG-3′

rAnk3-gRNA-3: 5′-CTTCACGCCGCTGTATATGG-3′

For AAV-mediated recombinant expression, the V5-tagged cDNA sequences were cloned into a pAAV-backbone under the CamKII promoter. For transient recombinant overexpression, the Flag-tagged cDNA sequences were cloned into pcDNA3 under the CAG promoter. Truncations were generated by PCR and infusion cloning. For mutation of all Ser/Thr sites in exon 7, DNA fragments were synthesized by Twist Bioscience and cloned into the same backbone by infusion cloning.

For zebrafish experiments, plasmids were assembled using NEBuilder HiFi DNA Assembly (New England Biolabs). To generate 4xUAS:Lifeact–mStayGold, the expression-vector backbone was PCR-amplified from an existing UAS:PSD-95-GFP plasmid ^62^. DNA fragments encoding Lifeact and mStayGold were PCR-amplified, purified, and assembled into the backbone. For PALM2 over-expression experiments, mouse *Palm2* cDNA was fused in-frame to either EGFP or mScarlet and inserted downstream of a 10xUAS promoter to generate 10xUAS:EGFP–PALM2 and 10xUAS:mScarlet–PALM2. Additional plasmids used in this study included 10xUAS:myrGFP, 10xUAS:myrmScarlet, and elavl3/HuC:Kalta4. All newly generated constructs were verified by sequencing.

### Immunohistochemistry

#### In cultured neurons

Neurons grown on coverslips were fixed either with 4% PFA for 15 min at 4°C, or with a glyoxal solution (3% glyoxal, 0.8% acetic acid, 20% methanol, pH 5.0) for 10 min at 4°C with post-fixation in prechilled methanol at −20°C for 10 min for Palm2 labeling. Cells were washed three times with PBS, followed by permeabilization by 0.3% Triton in PBS (PBST), blocking with 10% goat serum in PBST (blocking buffer) for 1 h at room temperature. Primary antibodies diluted in blocking solution were incubated at 4°C overnight, followed by three washes with PBS, incubation with secondary antibodies for 1 h at room temperature, three washes with PBS, and mounted onto a glass slide with VECTASHIELD HardSet Antifade Mounting Medium (Vector Laboratories Cat# H-1400-10). Cells were imaged using a Zeiss Apotome.2 microscope.

The following primary antibodies were used: rabbit polyclonal antibodies against βIV-spectrin (1:1000, M.N. Rasband, Baylor College of Medicine, Cat# βIV SD, RRID:AB_2315634), Palm2 (1:1000, Proteintech Cat# 18889-1-AP, RRID:AB_3085569); a guinea pig polyclonal antibody against AnkG (1:500, Synaptic Systems Cat# 386 004, RRID:AB_2725774); a chicken polyclonal antibody against MAP2 (1:1000, EnCor Biotechnology Cat# CPCA-MAP2, RRID:AB_2138173); mouse monoclonal antibodies against HA (1:1000, BioLegend Cat# 901505, RRID:AB_2565023), Palm2 (1:300, Abnova Cat# H00114299-M09, RRID:AB_566057), V5 (1:1000, Innovative Research Cat# R960CUS, RRID:AB_159298); a rat monoclonal antibody against HA (1:1000, Roche Cat# 11867423001, RRID:AB_390918). All Alexa-fluorophore-conjugated secondary antibodies were from Thermo Fisher Scientific and used at 1:500 dilution.

#### In mouse tissues

Mice were anesthetized using isoflurane and followed by transcardial perfusion with ice-cold PBS and different fixatives. For staining for Palm2 at the AIS, animals were perfused with a Glyoxal solution (Glyox+MeOH; 3% glyoxal, 0.8% acetic acid, 20% methanol) and post-fixed in the same solution at 4°C overnight. Tissues were dehydrated through serial sucrose gradient from 20% to 30% over two nights. OCT was used to embed the cryo-protected samples and sections were made at 10 µm thickness and directly attached onto the SuperFrost PLUS slides. Tissues sections were stored at -80°C until ready for staining. Briefly, slides were washed 3 times, 5 min each in PBS to remove OCT and rehydrate, then permeabilized with PBST for 3 times, 5 min each in a vertical slide mailer. A hydrophobic barrier was created using VECTASHIELD Immege Pen and slides were blocked with blocking buffer for 1 h at room temperature (RT). The subsequent steps were identical with *in vitro* staining.

The following primary antibodies were used: rabbit polyclonal antibodies against NF186 (1:500, Cell Signaling Technology Cat# 15034, RRID:AB_2773024), βIV-spectrin (1:500, M.N. Rasband, Baylor College of Medicine, Cat# βIV SD, RRID:AB_2315634), Palm2 (1:500, Proteintech Cat# 18889-1-AP, RRID:AB_3085569), Palm2 (1:200, Atlas Antibodies Cat# HPA074153, RRID:AB_2686669), Caspr (1:1000, Abcam Cat# ab34151, RRID:AB_869934), Olig2 (1:200, Millipore Cat# AB9610, RRID:AB_570666), and V5 (1:1000, Sigma-Aldrich Cat# V8137, RRID:AB_261889); a guinea pig polyclonal antibody against AnkG (1:500, Synaptic Systems Cat# 386 004, RRID:AB_2725774); a chicken polyclonal antibody against Neurofascin (1:500, R and D Systems Cat# AF3235, RRID:AB_10890736); mouse monoclonal antibodies against Palm2 (1:500, Abnova Cat# H00114299-M09, RRID:AB_566057), CC1 (1:500, Millipore Cat# OP80, RRID:AB_2057371), and PanNav (1:500, James Trimmer, University of California, Davis Cat# K58/35, RRID:AB_3073602). All secondary antibodies, Streptavidin, and α-bungarotoxin conjugated with Alexa-fluorophores were from Thermo Fisher Scientific and used at 1:500 dilution.

#### In human tissues

The formalin-fixed paraffin-embedded (FFPE) sample was sectioned at 6 µm thickness, deparaffinized in xylene three times for 5 min each, and rehydrated through graded ethanol solutions of 100%, 95%, 70%, and 50% ethanol for 10 min each, twice per concentration. Sections were then rinsed twice in deionized water for 5 min each. Antigen retrieval was performed in citrate buffer containing 10 mM sodium citrate and 0.05% Tween-20, pH 6.0, maintained at sub-boiling temperature for 15 min using a microwave. Slides were cooled slowly to RT while submerged in citrate buffer for 45 min, then washed twice in distilled water and once in 1 × PBS. A hydrophobic barrier was drawn around the tissue using an ImmEdge pen. Sections were permeabilized with 0.5% Triton X-100 in 1 × PBS for 15 min at RT and blocked for 1 h at RT in blocking buffer containing 0.3% Triton X-100 and 10% goat serum in 1 × PBS. Primary antibodies were diluted 1:80 in blocking buffer and incubated overnight at 4°C in a dark, sealed humidified chamber. The following primary antibodies were used: A rabbit polyclonal anti-βIV-spectrin antibody (M.N. Rasband, Baylor College of Medicine, Cat# βIV SD, RRID:AB_2315634), a mouse monoclonal anti-Palm2 antibody (Abnova Cat# H00114299-M09, RRID:AB_566057), and a mouse monoclonal anti-Caspr antibody (UC Davis/NIH NeuroMab Facility Cat# K65/35, RRID:AB_2877274).

After primary antibody incubation, slides were washed three times with 0.3% Tween-20 in 1 × PBS for 10 min each at RT. Alexa Fluor-conjugated secondary antibodies were diluted 1:100 in blocking buffer and incubated for 2 h at RT in a dark, sealed humidified chamber. The following secondary antibodies were used: A goat anti-mouse IgG2a Alexa Fluor 488 cross-adsorbed secondary antibody (Thermo Fisher Scientific Cat# A-21131, RRID:AB_2535771), a goat anti-mouse IgG1 Alexa Fluor 594 cross-adsorbed secondary antibody (Thermo Fisher Scientific Cat# A-21125, RRID:AB_2535767), and a goat anti-rabbit IgG Alexa Fluor 680 antibody (Molecular Probes Cat# A-21076, RRID:AB_2535736).

After secondary antibody incubation, slides were washed twice with 0.3% Tween-20 in 1 × PBS for 10 min each at RT, followed by three washes in 1 × PBS for 5 min each. To reduce autofluorescence, TrueBlack Lipofuscin Autofluorescence Quencher (Biotium Cat# 23007) was applied at 1:20 dilution in 70% ethanol for 2 min in the dark at RT. Slides were washed twice with 1 × PBS, mounted with VECTASHIELD HardSet Antifade Mounting Medium (Vector Laboratories Cat# H-1400-10), and imaged using a Zeiss Apotome.2 microscope.

### Stochastic Optical Reconstruction Microscopy (STORM)

Rat hippocampal neurons were cultured on #1.5H coverslips (Neuvitro, Cat# GG-18-1.5H-PRE) at a density of 1.2 × 10^4^ cells/cm^2^. At DIV 10, neurons were either directly fixed with 4% PFA in PEM buffer (80 mM PIPES pH 6.8, 5 mM EGTA, 2 mM MgCl2) for 10 min at RT, or detergent extracted with 0.5% Triton X-100 in PEM buffer for 5 min at RT before fixation. After fixation, coverslips were washed three times with 0.1 M phosphate buffer (0.1 M PB pH 7.4: 81 mM NaH_2_PO_4_ with 19 mM Na_2_HPO_4_) and blocked for 1 hour in blocking buffer (0.1 M PB containing 0.22% bovine gelatin and 0.1% Triton X-100). They were then incubated with rabbit anti-Palm2 primary antibody (1:150–1:200, Atlas Antibodies Cat# HPA074153, RRID:AB_2686669) diluted in blocking buffer at 4ºC overnight. The coverslips were washed three times with blocking buffer, incubated with secondary antibody diluted in blocking buffer (1:300, goat anti-rabbit Alexa Fluor 647) for 1 hour at RT, washed twice with blocking buffer, followed by sequential washes with 0.1 M PB and 0.05 M PB.

STORM imaging was performed on an N-STORM microscope (Nikon Instruments). Coverslips were mounted in a closed adhesive silicone chamber (CoverWell chamber, EMS Diasum Cat# 70334A) filled with STORM buffer (Smart Kit Buffer; Abbelight), and 40,000 images (256 × 256, ∼10 ms exposure time) were acquired. Nikon Elements N-STORM software was used to compute single-molecule locations. Image reconstruction (16 nm/pixel) was performed in Fiji-ImageJ using the ThunderSTORM and ChriSTORM plugins ^26,63^, and an 8-nm Gaussian blur was applied.

### In vivo proximity labeling

Biotin solution (2.5 mg/mL) was freshly prepared by dissolving biotin (Sigma-Aldrich, B4501) in Kolliphor EL, DMSO, and saline at a ratio of 1:1:18. The solution was heated to 95°C for 10 min, vortexed until completely dissolved, and cooled to room temperature before intraperitoneal (IP) injection (200 μL per mouse). Twenty-four hours after biotin administration, mice were euthanized by isoflurane overdose. For brain collection, mice were transcardially perfused with ice-cold Dulbecco’s phosphate-buffered saline (dPBS) before tissue dissection. Cerebra (olfactory bulbs and cerebellum removed), optic nerves, or the tibialis anterior muscles were rapidly dissected, snap-frozen in liquid nitrogen, and stored at −80°C until further processing.

#### Protein purification and affinity capture

Frozen tissues were homogenized in lysis buffer containing 50 mM Tris-HCl (pH 7.6), 1% SDS, 0.5% sodium deoxycholate, and 1% Triton X-100 supplemented with cOmplete protease inhibitor cocktail (Roche, 11836153001). Each cerebrum was homogenized in 2 mL lysis buffer on ice using a glass Dounce homogenizer (20 strokes each with loose and tight pestles). Optic nerves from 16-20 mice were pooled to generate one biological replicate and homogenized in 1 mL lysis buffer on ice using 100 strokes each with loose and tight pestles. Skeletal muscles were homogenized using a Bullet Blender (Next Advance). Homogenates were sonicated on ice in short pulses for a total of 4 min and centrifuged at 5,000 × g for 10 min at 4°C and the supernatant was collected. Proteins were precipitated by adding eight volumes of ice-cold methanol and incubating overnight at −80°C. Protein pellets were collected by centrifugation at 5,000 × g for 5 min at 4°C, washed in ice-cold methanol and incubated again at −80°C for 1 h, and centrifuged again. Residual methanol was allowed to evaporate completely before protein pellets were resuspended in 0.5% SDS. Protein concentrations were measured using the BCA assay and normalized before affinity purification.

Streptavidin-conjugated agarose beads were equilibrated by washing twice with dPBS and incubated with protein lysates (200 μL bead slurry per sample) for 3 h at room temperature with end-over-end rotation. Supernatant was removed by centrifugation at 3,000 × g for 2 min. Beads were washed twice with 2% SDS in dPBS, twice with 8 M urea in dPBS, twice with 2 M NaCl in dPBS, and four times with 100 mM triethylammonium bicarbonate (TEAB; Sigma, T7408). After the final wash, beads were stored at −80°C until further processing for mass spectrometry analysis.

### Mass spectrometry of biotinylated proteins

#### On-bead digestion

Sample-incubated streptavidin-conjugated agarose beads were resuspended in 5 mM dithiothreitol (DTT) in 100 mM NH_4_HCO_3_ and incubated for 30 min at room temperature. Iodoacetamide was then added to a final concentration of 7.5 mM, and samples were incubated for an additional 30 min in the dark at room temperature. 0.5 μg sequencing-grade trypsin (Promega) was added to each sample, and proteins were digested overnight at 37°C. The supernatants were collected, and the beads were subjected to a second digestion with an additional 0.5 μg trypsin in 100 mM NH_4_HCO_3_ for 2 h. Peptides from both consecutive digestions were combined, purified by solid-phase extraction using C18 ZipTips (Millipore Cat# ZTC18S096), and resuspended in 0.1% formic acid for liquid chromatography-tandem mass spectrometry (LC-MS/MS) analysis.

Trypsin digested peptides were analyzed using an Orbitrap Fusion Lumos mass spectrometer (Thermo Scientific) connected to a NanoAcquity Ultra Performance UPLC system (Waters). A 15-cm EasySpray C18 column (Thermo Scientific) was used to resolve peptides (90 min gradient with 0.1% formic acid in water as mobile phase A and 0.1% formic acid in acetonitrile as mobile phase B). The mass spectrometer was operated in data-dependent acquisition mode, automatically switching between MS and MS/MS scans. MS spectra were acquired over an m/z range of 375–1500 at a resolution of 120,000. For each full MS scan, multiply charged ions exceeding the threshold of 2 × 10^4^ were selected for MS/MS in cycles of 3 s with an isolation window of 0.7 m/z. Precursor ions were fragmented by higher-energy collisional dissociation (HCD). MS/MS spectra were acquired in centroid mode at a resolution of 60,000 with a lower limit of m/z 110. A dynamic exclusion window was applied which prevented the same m/z (mass tolerance 30 ppm) from being selected for 30 s after its acquisition.

#### Data analysis

Peak lists were generated using PAVA software (cite) and searched against the mouse subset of the SwissProt database (SwissProt release 2017.11.01) using Protein Prospector (cite). Database searches were performed with the following parameters: a precursor mass tolerance of 10 ppm, a fragment mass tolerance of 30 ppm, Carbamidomethylation of cysteine was specified as a fixed modification, acetylation of the N terminus of the protein, pyroglutamate formation from N-terminal glutamine, and oxidation of methionine as variable modifications. All spectra identified as peptide matches for a given protein were reported, and the total number of spectra (Peptide Spectral Matches, PSMs) was used for label-free quantification of protein abundance.

### Brain membrane extraction

Rat and mouse brain membrane fractions were purified following previously described method. Briefly, animals were euthanized by isoflurane overdose, and the brains were immediately dissected out and snap-frozen in liquid nitrogen and stored at -80°C. Brain was homogenized with a Dounce homogenizer (20 × loose + 20 × tight) on ice using a sucrose homogenization buffer (320 mM sucrose, 5 mM Sodium phosphate (NaPi), 1 mM Sodium fluoride, 1 mM Sodium orthovanadate, supplemented with cOmplete protease inhibitor cocktail). Post-nuclear supernatant was obtained by centrifugation at 700 × g for 10 min. Membrane-enriched fraction was further pelleted from the supernatant by centrifugation at 27,200 × g for 90 min at 4°C. The pellet was resuspended in homogenization buffer, and the concentration was measured using BCA assay and adjusted. The brain membrane fractions were stored at -80°C for regular immunoblotting. The samples were analyzed by label-free Data-Independent Acquisition (DIA) Mass Spectrometry or immunoblotting.

### Data Independence Acquisition (DIA) mass spectrometry

#### Sample processing

The protein lysate was diluted and denatured in final 5% SDS buffer. The protein sample was reduced in 5 mM DTT at 90°C, 5 min followed by alkylation in 10 mM IAA for 15 min in dark at room temperature. The protein lysate was concentrated, processed on S-Trap™ Micro Spin Column (CO2-micro, ProtiFi, NY) as per manufacturer protocol. The protein digestion was carried out using trypsin and LysC enzyme mixture at 37°C overnight. The peptide concentration was measured using the Pierce™ Quantitative Colorimetric Peptide Assay (Thermo Scientific Cat# 23275). The peptides were dried in a speed vac and dissolved in MS loading buffer (0.015% DDM prepared in 0.1% formic acid) to make a 100 ng/μl concentration.

LC-MS/MS analysis was conducted on a nanoElute2 UHPLC system (Bruker Daltonics, Germany) coupled online to timsTOF Ultra2 mass spectrometer (Bruker Daltonics, Germany) equipped with a CaptiveSpray nanoelectrospray ion source. The data acquisition was performed in diaPASEF mode. The peptide samples were loaded onto a 25 cm × 75 µm × 1.5 µm PepSep column (Bruker) with trap-and-elute method and separated using a 28 min active gradient time. The diaPASEF acquisition scheme included m/z range (400 to 1000 m/z) with a mobility range (0.64 to 1.37 1/K0). The MS/MS method employed 24 DIA windows, resulting in a cycle time of 0.96 s. The diaPASEF raw data was processed using DIA-NN (v 2.2.0) in a library-free search mode. The FASTA file was downloaded from GENCODE database (Release M32). The precursor ions were generated from FASTA in silico digest with deep learning-based spectra, retention times (RTs), and ion mobilities (IMs) prediction. The proteolytic enzyme Trypsin/P was selected, allowing for up to one missed cleavage. Variable modification of oxidation (M) and protein N-term acetylation was allowed. Cysteine carbamidomethylation was enabled as a fixed modification. Parameter specifics include peptide lengths ranging from 7 to 30, precursor charges from 1 to 4, precursor m/z from 300 to 1800, and fragment ion m/z from 200 to 1800. The MBR was enabled and precursor identification was set at 1% FDR. The DIA-NN precursor-level report file was processed through gpGrouper algorithm ^64^to obtain protein level quantification using the iBAQ approach. The Ms1.Normalised values were used for iBAQ calculation.

#### Data analysis

After median normalization of each sample, differential analysis was performed on log transformed values using a variance moderated t-test as implemented in limma with both robust and trend variance estimation settings enabled ^65^. Missing value imputation was performed with a random draw from a normal distribution N(1.8*mu_observed, 0.35*sd_observed). Multiple-hypothesis testing correction was controlled at the False Discovery Rate (FDR) with the Benjamini–Hochberg procedure.

### Immunoblotting

Protein samples were boiling in SDS reducing sample buffer at 95°C for 10 min, followed by centrifugation at 2,500 × g for 5 min before loading on 6% or 14% polyacrylamide gels with SDS (SDS-PAGE) to resolve proteins of interest. 20 µg of reduced total protein was resolved on SDS-PAGE by electrophoresis and transferred onto 0.45 µm nitrocellulose membrane using the Bio-Rad Trans-blot system. Ponceau S was briefly added to the membrane to evaluate the quality of transfer and for normalization by total protein, then washed away using TBST (TBS with 0.1% Tween-20). The membrane was blocked with Blotto solution (5% skim milk in TBST) for 1h at RT. Primary antibodies against proteins of interest were diluted in Blotto solution and incubated with the membrane at 4°C overnight. The membrane was then washed with TBST for three times, 10 min each, and then incubated with secondary antibodies in Blotto solution for 1 h at RT. The membrane was then washed with TBST three times and PBS twice before imaging with an Azure 600 imager or a Licor Odyssey Fc imager. Chemiluminescence signal was developed using SuperSignal West Pico PLUS (Pierce Cat# 34580). Restore Western Blot Stripping Buffer (Thermo Scientific Cat# 21059) was used following manufacturer’s instruction to re-incubate the same membrane with different primary antibodies.

The following primary antibodies were used: rabbit polyclonal antibodies against Palm2 (1:4000, Proteintech Cat# 18889-1-AP, RRID:AB_3085569), Palm2 (1:1000, Atlas Antibodies Cat# HPA074153, RRID:AB_2686669), AnkB (1:2000, Rasband Lab, E1588), Caspr (1:2000, Abcam Cat# ab34151, RRID:AB_869934), and GFP (1:4000, Proteintech Cat# 50430-2-AP, RRID:AB_11042881); a guinea pig polyclonal antibody against AnkG (1:2000, Synaptic Systems Cat# 386 004, RRID:AB_2725774); a chicken polyclonal antibody against Neurofascin (1:2000, R and D Systems Cat# AF3235, RRID:AB_10890736); mouse monoclonal antibodies against Palm2 (1:2000, Abnova Cat# H00114299-M09, RRID:AB_566057), βII-spectrin (BD Biosciences Cat# 612563, RRID:AB_399854), and Flag (1:4000, Sigma-Aldrich Cat# F3165, RRID:AB_259529); and an HRP-conjugated mouse monoclonal antibody against β-Actin (1:2000, Santa Cruz Biotechnology Cat# sc-47778, RRID:AB_626632).

HRP-conjugated secondary antibodies were from Jackson ImmunoResearch, Alexa-fluorophore-conjugated secondary antibodies were from Thermo Fisher Scientific, and IRdye800-conjugated donkey anti-guinea pig IgG (LICORbio Cat# 926-32411, RRID:AB_1850024) was from Licor. All secondary antibodies were used at 1:4000 dilution.

### Lipid raft assays

#### Detergent solubilization

Brain membrane homogenates from P45 mice were diluted to 1 mg/mL in lysis buffer (20 mM Tris-HCl pH 8.0, 10 mM EDTA, 150 mM NaCl, 10 mM iodoacetamide supplemented with cOmplete protease inhibitor cocktail) containing 1% Triton X-100, and incubated for 1 h via end-to-end rotation at 4°C or 37°C. Lysates were centrifuged at 13,000 × g for 30 min at 4°C to separate the detergent-soluble and detergent-insoluble fractions. The insoluble fraction was resuspended in lysis buffer to the original starting volume. Equal volumes of the detergent-soluble fraction and the resuspended insoluble fraction were mixed with 2 × reducing sample buffer (125 mM Tris-HCl pH 6.8, 20% glycerol, 0.04% bromophenol blue, 4% SDS, 200 mM dithiothreitol) and incubated for 5 min at 95°C. 5 µg brain homogenate or the equivalent of 5 µg protein split between the soluble and insoluble fractions for each condition was used for immunoblotting. Immunoblotting was performed as described above. Samples were run on 4-20% gradient Tris-glycine polyacrylamide gels and transferred to 0.45 µm nitrocellulose membranes (BioRad Cat# 1620115).

The following primary antibodies were used: mouse anti-Palm2 (1:500, Abnova Cat# H00114299-M09, RRID:AB_566057), rabbit anti-Caspr (1:1000, Abcam Cat# ab34151, RRID:AB_869934), and guinea pig anti-Ankyrin G (1:1000, Synaptic Systems Cat# 386 004, RRID:AB_2725774). HRP-conjugated secondary antibodies were used at 1:1000 dilution. Blots were developed with chemiluminescent substrate and visualized with an Azure 600 imaging system.

#### Cholesterol extraction

Brain membrane homogenates from P45 mice were diluted to 1 mg/mL protein in lysis buffer without 1% Triton X-100, with or without 0.2% (w/v) saponin treatment for 30 min at 4°C via end-to-end rotation. Samples were centrifuged at 13,000 × g for 10 min at 4°C. The saponin-treated and untreated pellets were resuspended in 1 mL of lysis buffer with 1% Triton X-100 and rotated for 1 h at 4°C. After centrifugation at 13,000 × g for 30 min at 4°C, the detergent-soluble and detergent-insoluble fractions were separated. The insoluble fractions were stored at - 80°C until ready for processing. The soluble fractions were precipitated with nine volumes of chilled ethanol and incubated overnight at -80°C. After centrifugation at 12,500 × g for 30 min at 4°C, the supernatant was decanted and the pellet containing precipitated soluble fractions was allowed to air-dry. The pellets from both soluble and insoluble fractions were resuspended in equal volumes of 2 × reducing sample buffer and incubated at 95°C for 5 min. 10 µg brain homogenate and 10 µL of the soluble and insoluble fractions from the saponin-treated or untreated homogenates were loaded on 4-20% gradient Tris-glycine gels. Immunoblotting was performed as described above.

#### Sucrose gradient centrifugation

The sucrose gradient preparation was adapted from Schafer et al. ^35^. Sucrose solutions were prepared in lysis buffer without Triton X-100. Detergent-resistant membrane fractions were isolated by solubilizing brain membrane homogenates from P45 mice in lysis buffer containing 1% Triton X-100 at a concentration of 1 mg/mL protein for 1 hr at 4°C via end-to-end rotation. After centrifugation at 13,000 × g for 30 min at 4°C, the pellet was resuspended in 2.25 mL lysis buffer without Triton X-100 and transferred to an Ultra-Clear centrifuge tube (Beckman Coulter Cat# 344060), mixed with 2.25 mL 2 M sucrose, overlaid with 4.5 mL 1 M sucrose, and then overlaid with 3.6 mL 0.5 M sucrose. Samples were centrifuged at 40,000 RPM for 19 h at 4°C in an Optima XPN-100 Ultracentrifuge (Beckman Coulter) with a SW 40 Ti swinging-bucket rotor. 0.9 mL fractions were collected starting from the top of the tube, then ethanol-precipitated as described for the cholesterol extraction experiments. The precipitated pellets were resuspended in 2 × reducing sample buffer and incubated at 95°C for 5 min. 5 µL of each fraction was loaded on a 4-20% gradient Tris-glycine gel. Immunoblotting was performed as described for the temperature sensitivity assay.

### Co-immunoprecipitation

#### In vitro

For *in vitro* CoIP experiments, HEK293T cells were seeded at the density of 1.6 × 10^6^ cells/well in 6-well plates and transfected the following day with 2.5 µg total DNA per well using Lipofectamine 3000 (L3000001; Thermo Fisher Scientific) in Opti-MEM, followed by media change the next day. 48 h after transfection, cells were first washed with ice-cold dPBS and lysed directly in the well with Co-IP buffer (20 mM Tris-HCl pH 8.0, 150 mM NaCl, 1% Triton X-100, 10 mM EDTA pH 8.0, supplemented with cOmplete protease inhibitor cocktail and PMSF). After 15 min incubation on ice, the lysed cells were scrapped off and collected into an Eppendorf tube and further incubated at 4°C for 10 min with end-to-end rotation. Supernatant were collected following centrifugation at 20,000 × g for 10 min at 4°C. Protein concentration was measured by BCA and adjusted with Co-IP buffer. 10% of the supernatant was kept as input. For the remaining IP samples, 4 μl of rabbit anti-GFP antibody (Proteintech Cat# 50430-2-AP, RRID:AB_11042881) was added and the lysate antibody mixture was rotated end-to-end at 4°C overnight. Antibodies were then immunoprecipitated using Protein A (Cat# 28951378; Cytiva) magnetic sepharose beads equilibrated and pretreated with 0.1% BSA 1 h before being added to the lysate antibody mixture and rotated end-to-end at 4°C for 1 h. The samples were placed on a magnetic stand, and the supernatant was aspirated to remove unbound proteins. Protein-bound beads were washed with the supplemented Co-IP buffer for seven times, each with end-to-end rotation for 5 min at 4°C. After the final wash, the supernatant was removed and sample buffer was added to the beads and boiled at 95°C for 5 min. Immunoblotting was performed as described above.

The following primary antibodies were used for immunoblotting: a rabbit polyclonal antibody against GFP (1:2000, Proteintech Cat# 50430-2-AP, RRID:AB_11042881), and a mouse monoclonal antibody against Flag (1:2000, Sigma-Aldrich Cat# F3165, RRID:AB_259529).

#### In vivo

For CoIP of endogenous proteins from mouse brain, P21 mice were euthanized via isoflurane overdose. Freshly dissected brains from three mice were pooled together as a biological replicate and immediately homogenized (20 strokes each with loose and tight pestles) in a Dounce homogenizer on ice in 7 mL lysis buffer: 20 mM Tris HCl (pH 8.0), 137 mM NaCl, 1% NP-40, 2 mM EDTA, supplemented with cOmplete protease inhibitor cocktail. The homogenates further incubated at 4°C with end-to-end rotation for 30 min before centrifugation at 18,000 × g for 15min to collect the supernatant. Protein concentration was measured using BCA and adjusted to 2 mg/mL. 10% of the sample was kept as input. The remaining samples for IP were pre-cleared with protein A/G agarose beads by end-to-end rotation for another 15 min at 4°C, followed by centrifugation at 2,500 × g for 2 min at 4°C. 1 mL of the cleared lysate was used for each IP reaction. 4 µL primary antibodies were added and incubated at 4°C overnight by end-to-end rotation, including guinea pig anti-AnkG (Synaptic Systems Cat# 386 004, RRID:AB_2725774), or an isotype-matched IgG control antibody against Trim46 (Synaptic Systems Cat# 377 308, RRID:AB_2924929).

Protein A/G agarose beads were equilibrated by washing twice with dPBS and incubated with protein lysates (20 μL bead slurry per sample) for 2 h at 4°C with end-over-end rotation. Beads were Pelleted at 2,500 × g for 2 min at 4°C and supernatant was discarded. Protein-bound beads were then washed for 6 times in the following wash buffer: 10 mM Tris (pH 7.4), 1mM EDTA, 1 mM EGTA (pH 8.0), 150 mM NaCl, 0.5% NP-40, 0.2 mM sodium orthovanadate, supplemented with cOmplete protease inhibitor cocktail. Each wash was carried out through end-to-end rotation at 4°C for 5 min followed by centrifugation at 2,500 × g at 4°C for 2 min. After the final wash, 4 × SDS-sample buffer was added, and the samples were boiled at 95°C for 5-10min and ran on a 6% or 12% SDS-PAGE as described above in immunoblotting.

The following primary antibodies were used for immunoblotting: a guinea pig polyclonal antibody against AnkG (1:2000, Synaptic Systems Cat# 386 004, RRID:AB_2725774), and a rabbit polyclonal antibody against Palm2 (1:2000, Proteintech Cat# 18889-1-AP, RRID:AB_3085569).

### RNA purification and cDNA synthesis

Total RNA was isolated using the Qiagen RNeasy Mini kit (Qiagen Cat# 74104) following the manufacturer’s instructions. The RNA quality was checked using the NanoDrop spectrophotometer (Thermo Fisher Scientific) and Agilent Bioanalyzer 2100 or Tape Station. cDNA was synthesized by reverse transcription of the total RNA using SuperScript IV VILO MasterMix with genomic DNA removal using the ezDNase Enzyme (Invitrogen Cat# 11766050), following manufacturer’s instruction.

#### qRT-PCR

Quantitative real-time PCR (qRT-PCR) was performed on the StepOnePlus Real-Time PCR System with the Bio-Rad iTaq Universal SYBR Green Supermix. Melting curve plots were examined for amplicon specificity. The 2^-ΔΔCT^ quantification method was used to examine the relative expression of genes-of-interest, with *Gapdh* serving as the internal control. The following primers were used:

*mPalm2-exon 3* Forward: AGAAAGAGGCAGACGGAAA-TAG

*mPalm2-exon 3-4* Reverse: GAAGCACTTTGGACTTGGAATG

*mPalm2-exon 7* Forward: TGACGGTACCAAAGTAGTG-TATG

*mPalm2-exon 7* Reverse: CTGTCCCGCCTTCTGAATTA

*mGapdh* Forward: ATGACATCAAGAAGGTGGTG

*mGapdh* Reverse: CATACCAGGAAATGAGCTTG

#### Sanger sequencing

Sanger sequencing of the Palm2 WT and *Palm2*^*ΔEx4*^ samples were performed on the PCR-amplified sequence from synthesized cDNA. Briefly, a pair of primers were designed to amplify the coding sequence of Palm2 in both *Palm2*^*ΔEx4*^ mice and WT control mice. After gel electrophoresis and gel purification, the following primer was used for sanger sequencing:

*mPalm2-seq*: ATGGCAGAGGCGGAATT

### Library prep and long-read Nanopore sequencing

Briefly, full-length cDNA libraries were generated from 10 ng of total RNA using the Oxford Nanopore Technologies cDNA-PCR barcoding workflow (SQK-PCB114.24) and sequenced on a PromethION platform (FLO-PRO114M flow cells) according to the manufacturer’s instructions. Libraries were quality controlled using Qubit and Agilent Bioanalyzer/TapeStation instruments prior to equimolar pooling and sequencing of four cDNA libraries per flow cell.

The cDNA-PCR sequencing workflow employs a strand-switching approach together with a barcoding kit to generate full-length cDNA from total RNA, enabling users to pool up to 24 samples in a single sequencing run. During the strand-switching step, a unique molecular identifier (UMI) is incorporated. The resulting cDNA is then converted into double-stranded form and amplified by PCR using primers that contain 5′ tags. Library preparation was performed in four steps: (1) reverse transcription with strand-switching to generate full-length cDNA from total RNA, incorporating UMIs. (2) PCR amplification and enrichment of full-length cDNA using rapid barcode attachment primers. (3) rapid ligation of sequencing adapters to amplified products; and (4) flow cell priming and loading at 50 fmol equimolar pooled barcoded cDNA for sequencing. Pod5, FastQ files and QC reports were generated using MinKNOW sequencing software which controls the device, data acquisition and real-time basecalling.

### Transcript isoform analysis

Raw sequencing reads from individual sequencing runs were combined for each biological sample prior to downstream analysis. Read quality was assessed using FastQC (Babraham Bioinformatics). Sequencing reads were aligned to the mouse reference genome (GRCm38/mm10) using minimap2 (RRID:SCR_018550) (v2.31) ^66^ with -ax splice -s 40 -G 350k -t 25 –MD. Transcript isoforms were reconstructed using the Full-Length Alternative Isoform Analysis of RNA (FLAIR, v2.0) ^67^. Splice junction correction was performed against the GENCODE mouse annotation (release M25) (RRID:SCR_014966), followed by transcript collapsing using corrected read alignments from all samples to generate a unified high-confidence transcriptome using the default flair -correct and flair -collapse options. Isoform abundance was quantified using the FLAIR quantification module, and isoform-level read counts were generated for each sample and used for downstream analyses of transcript expression and isoform usage. Gene-specific transcript analyses were performed by extracting all transcript isoforms mapping to the *Palm2*-*Pakap* locus from the collapsed transcript annotation. Isoform structures were visualized using the Integrative Genomics Viewer (RRID:SCR_011793). Alternative splicing events and exon usage were further examined using IGV Sashimi plots to visualize exon-exon junction support and exon inclusion levels across genotypes. Isoform usage was quantified as the proportion of read counts assigned to each isoform relative to the total number of counts mapped to the corresponding gene within each biological sample. Transcript annotations, including transcript length, predicted coding sequence length, exon composition, and inclusion or exclusion of exons of interest, were integrated with isoform abundance to compare transcript diversity and alternative splicing events in both wild-type and *Palm2*^*ΔEx4*^ tissues.

### Electrophysiology

The compound action potential (CAP) recording was performed as previously described. Briefly, optic nerves from the P21 Palm2 WT or *Palm2*^*ΔEx4*^ mice were freshly dissected out between the optic nerve head immediately behind the eyeball and the optic chiasm. The dissected optic nerves were immediately submerged in oxygenated Locke’s solution bath at 25°C (154 mM NaCl, 2 mM CaCl_2_, mM KCl, 10 mM HEPES, pH adjusted to 7.4 with NaOH) supplemented with 1 mg/mL glucose. The unmyelinated segment at the optic nerve head was trimmed away and the optic chiasm was also removed before sealed into the suction electrodes. The optic nerve was stimulated on the anterior end and recorded on the posterior end.

Increasing currents were applied using Molecular Devices: MultiClamp 700B Microelectrode Amplifier (RRID:SCR_018455) to reach the supra-maximal threshold, where the CAP was subsequently recorded. Data visualization and analysis was performed in Clampex 10.7 and Clampfit 10.7. The CAP conduction velocity was calculated by dividing the length of the optic nerve between the two electrodes by the latency to maximal peak from onset of stimulation.

### Behavioral analysis

#### Touch-evoked response

Zebrafish larvae injected with control or *palm2*-targeting sgRNAs were examined at 3 and 5 dpf for gross morphological abnormalities. Touch-evoked escape responses were elicited by applying a blunt probe to the head and tail. Responses were recorded and scored using predefined criteria.

#### Righting reflex

P8 mice were picked up and gently placed on a warm pad on their back. The latency before then successfully land on all four limbs were recorded and quantified to analyze general health, motor, and neurological function.

#### Open field

The open field test was carried out to assess general locomotor function in adult mice. Briefly, mice were placed in the center of an open Plexiglass arena (40 × 40 cm, height 20 cm) and allowed to explore freely for 10 minutes. The inner zone was defined as the center 20 × 20 cm area. A camera was mounted above the arena for automatic tracking and data analysis using the AnyMaze software. The parameters reported here are total distance traveled and time spent mobile in the inner zone. Both male and female mice were included for each genotype. No sex differences were identified.

#### Rotarod

The rotarod test was conducted as previously described, with the following modifications. 2-month-old adult mice were placed on an accelerating rotarod to examine their motor function and motor learning. Animals were acclimated to walking on the rod at a constant speed of 4 rpm for 3 min before returning to home cage for a 30 min break and beginning of the 7 trials across three days. On day 1 and day 2, mice, three trials were conducted (with the rotation speed increasing from 4 rpm to 40 rpm during a 5 min test period), with each trial separated by an 1 h rest for the mice. On day 3, the final trial was conducted. The latency to fall from the rotarod was recorded for all seven trials. The latency to fall was triggered by the lever or manually recorded if the animal passively clutched onto the rod for two full rotations.

## QUANTIFICATION AND STATISTICAL ANALYSIS

### Image analysis

Polarity index of Palm2 was defined by the ratio of correct mean intensity at the AIS to correct mean intensity at proximal dendrites. AIS was defined by endogenous AnkG or βIV-spectrin labeling. Correct mean intensity (CMF) was obtained using mean intensity in target regions extracted by the background of the same image. At least three independent experiments were performed.

AIS length was defined by AnkG or βIV spectrin signal and measured in FIJI/ImageJ using the segmented line function. Fluorescence intensity was measured using Rolling Ball Background Subtraction (radius = 50 pixels) in FIJI/ImageJ.

Node number was measured in FIJI/ImageJ using autothresholding and the particle analysis function. Node length was measured by determining the longitudinal extent of individual PanNav-positive nodes. Manual quantification was performed at randomly selected regions-of-interest using the cell-counter function.

Sholl analysis was performed by quantifying the number of intersections between the skeleton and concentric circles positioned at fixed distances from the soma boundary. Overlay images were generated for visual verification of soma annotation, skeletonization, and Sholl circle placement. Specifically, GFP-channel of neuron images were used as program input. Soma location and size were manually selected by an interactive graphical interface. Neurons with overlapping processes were not selected for analysis. Neuronal processes were enhanced with a Sato ridge filter, thresholded using a percentile-based cutoff, cleaned to reduce small objects and artifacts, and skeletonized to one-pixel-wide structures. The soma region was removed prior to skeletonization and only the skeleton connected to soma was retained for analysis. A Python-based pipeline was used, and the code is available on: https://github.com/Xiaoye-the-tubulin/Sholl-analysis-Xiaoyun-zoomed-image.git.

Time-lapse image analysis for Lifeact dynamics in zebrafish was performed in Fiji/ImageJ. Lifeact-positive puncta located on mScarlet-positive axons were identified. Regions of interest were manually placed around individual punctum, and fluorescence intensity was measured throughout the 3-min imaging period. Local background fluorescence was subtracted from each measurement, and changes in Lifeact fluorescence over time were quantified for each region of interest.

Myelin sheath morphology in zebrafish was analyzed from sparsely labeled oligodendrocytes in the spinal cord. Individual sheaths were traced through the complete confocal z-stack using the segmented-line tool in Fiji/ImageJ, and the corresponding regions of interest were saved for analysis. Custom ImageJ macros were used to export sheath-length measurements, which were subsequently processed using custom MATLAB scripts. For each oligodendrocyte, the analysis quantified the number of myelin sheaths, the length of each individual sheath, the mean sheath length, and the total sheath length. Oligodendrocytes from segment 15 were chosen for analysis.

Densitometry analysis of immunoblots were performed in FIJI/ImageJ as previously described. Due to a potential effect of Palm2 on cytoskeleton dynamics, and a significant and systemic change in β-Actin observed in the *Palm2*^*ΔEx4*^ mice, normalization for total protein loaded was performed against Ponceau S staining instead.

### Statistical analysis

Statistical analyses were performed using GraphPad Prism 11 (RRID:SCR_002798), with additional data processing performed in Microsoft Excel, Python (RRID:SCR_008394), and R. No statistical methods were used to predetermine sample size, but sample sizes were comparable to those reported in previous studies. For *in vivo* analyses, at least three pairs of age- and sex-matched littermates were analyzed per genotype unless otherwise indicated, and the exact sample size for each experiment is provided in the figure legends. For *in vitro* analyses, at least three independent cultures were analyzed unless otherwise indicated. For electrophysiological recordings, traces from 14 control and 14 *Palm2*^*ΔEx4*^ optic nerves were analyzed.

Data normality was checked using the Shapiro–Wilk test. Homogeneity of variance was assessed using the F-test for two-group comparisons or the Brown–Forsythe test for comparisons among three or more groups. Proper data transformations (e.g., log transformation) were applied to meet normality assumptions when appropriate; otherwise, corresponding non-parametric methods were used. Outliers were identified using the default ROUT method (Q = 1%) in GraphPad Prism 11 only when indicated in the figure legends; identified outliers were excluded from subsequent analysis, and results for excluded outliers are also reported in the Source Data file. Individual data points are shown whenever possible, and data are presented as mean ± SEM.

For comparisons between two groups, unpaired Student’s t-tests were used when data met assumptions of normality and equal variance and the N was equal to or larger than 6; Welch’s t-tests were used when variances were unequal or the N was smaller than 6; and Mann–Whitney U tests were used when data did not meet the assumption of normality. For comparisons among three or more groups, one-way ANOVA with pairwise comparisons and Tukey’s correction was used for normally distributed data; when normality was not met, the Kruskal–Wallis rank test with Dunn’s multiple-comparison correction was applied. Two-way ANOVA with Bonferroni’s multiple-comparison correction was used for experiments with two independent variables. Chi-square tests were used for categorical data such as the survival analysis by genotype. Unless otherwise stated, all statistical tests were two-tailed at a significance level of 0.05. Both unadjusted P values and, where applicable, estimated false discovery rates adjusting for multiple endpoints are reported. The specific statistical test used for each experiment is indicated in the corresponding figure legend.

## Supplemental Figures

**Figure S1.**
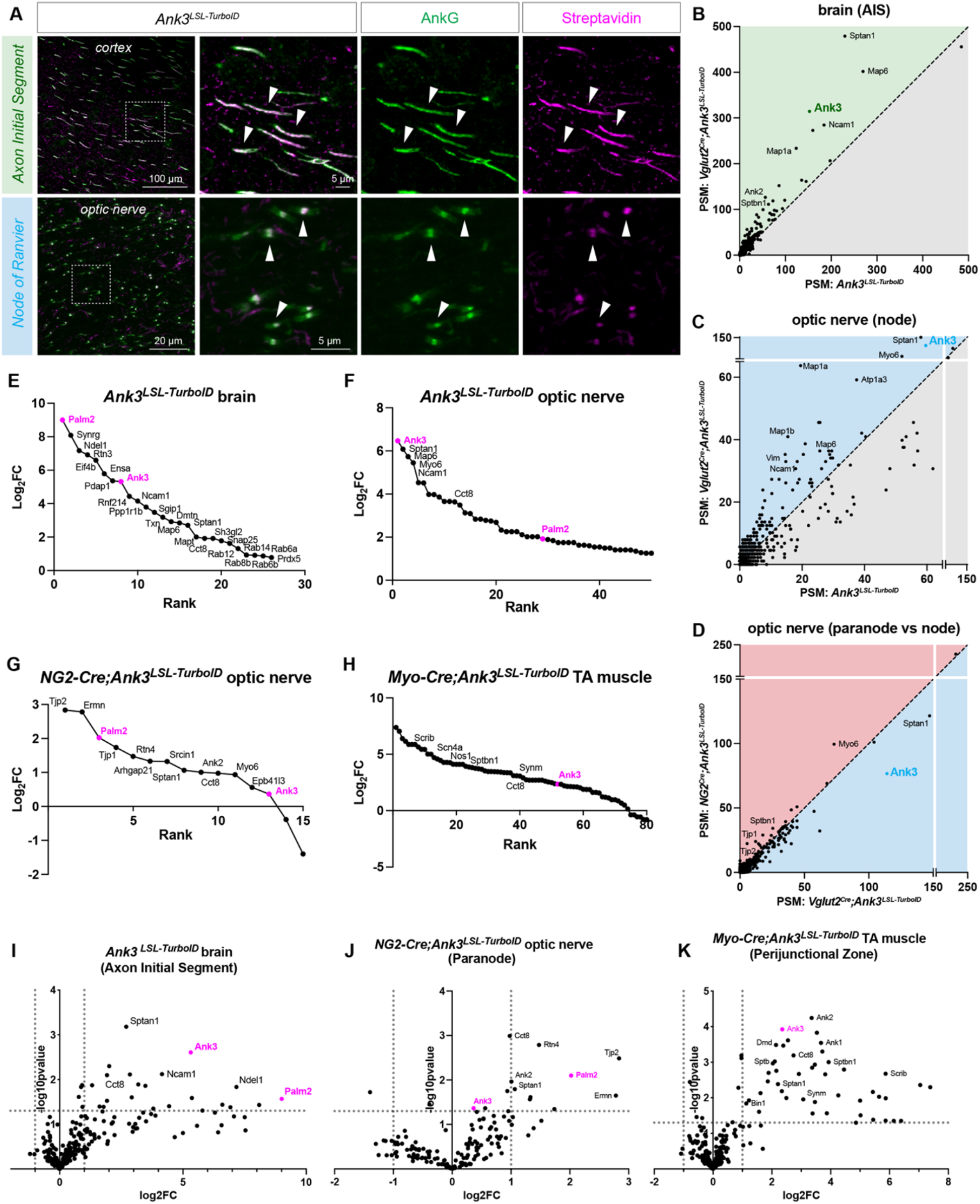
AnkG-proximity proteomics at the axon initial segments, nodes of Ranvier, paranodes, and perijunctional zone. (A) Immunostaining of adult cortex showing baseline biotinylation of some AIS and nodes of Ranvier in *Ank3*^*LSL-TurboID*^ mice. Arrowhead indicates colocalization of streptavidin (magenta) with AnkG (green) at the AIS. (B-D) PSMs of candidate AIS proteins (B) and nodal proteins (C) in *Vglut2*^*Cre*^*;Ank3*^*LSL-TurboID*^ versus *Ank3*^*LSL-TurboID*^ mouse brain; and PSM of candidate paranodal in *NG2*^*Cre*^*;Ank3*^*LSL-TurboID*^ versus candidate nodal proteins in *Vglut2*^*Cre*^ *Ank3*^*LSL-TurboID*^ mice (D). (E-H) Top candidate proteins ranked by log_2_ FC at the AIS (E), node (F), paranodes (G), and perijunctional zone (H). Volcano plot showing top candidate AIS proteins in the *Ank3*^*LSL-TurboID*^ mouse brain. (I-K) Volcano plot showing top candidate proteins enriched in *Ank3*^*LSL-TurboID*^ mouse brains, *NG2*^*Cre*^;*Ank3*^*LSL-TurboID*^ mouse optic nerves, and *Myo-* ^*Cre*^*;Ank3*^*LSL-TurboID*^ mouse muscles. Scale bars are indicated in each panel.

**Figure S2.**
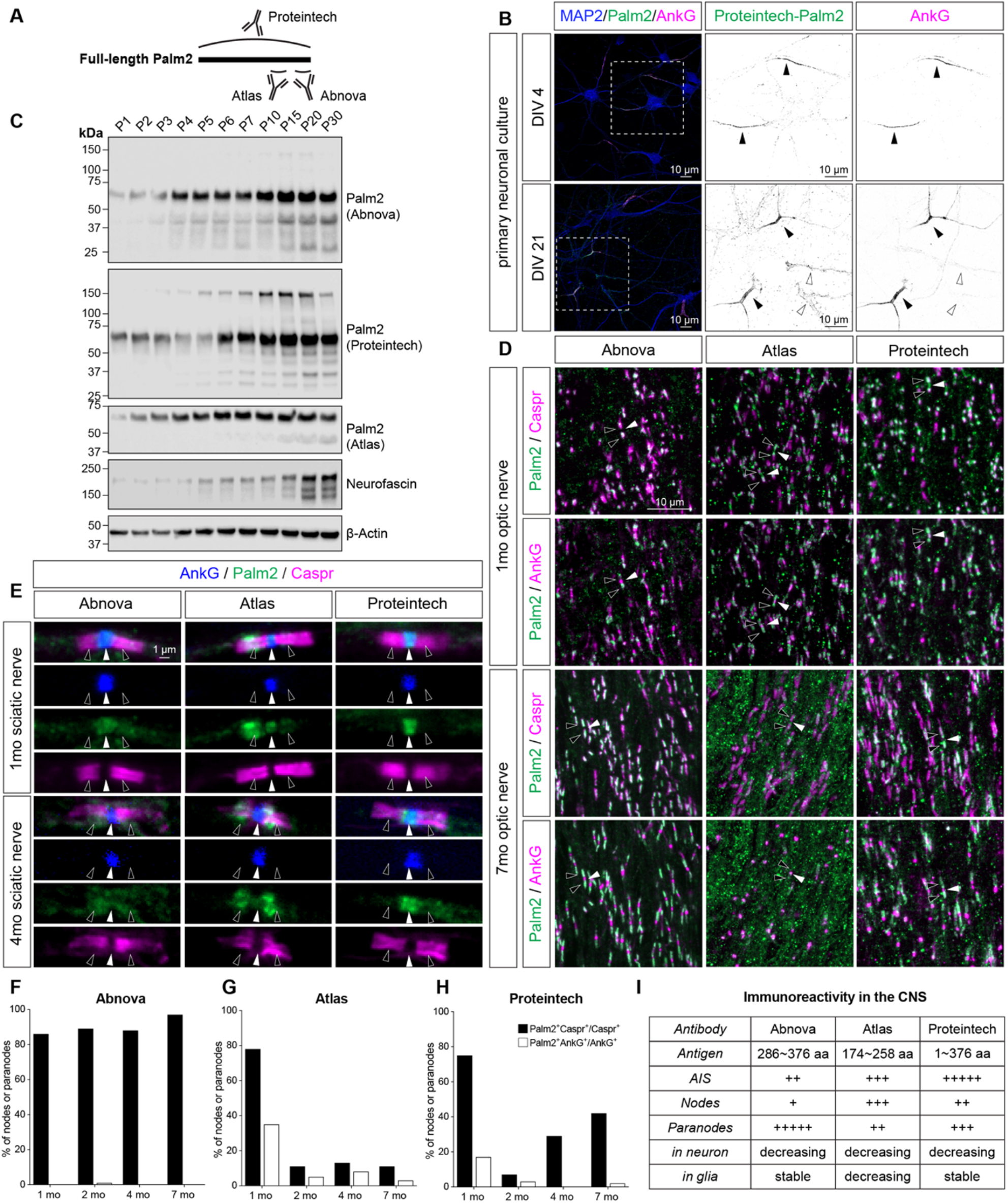
Validation of Palm2 expression with three independent antibodies across different developmental stages and tissue types. (A) Schematic of the regions in Palm2 that were used to generate the three commercial Palm2 antibodies used here. (B) Immunostaining of endogenous Palm2 shows colocalization of Palm2 (green) with AnkG (magenta) at DIV 4 and 21. AIS are indicated by black arrowheads. White arrowheads show low levels of Palm2 outside the AIS. (C) Immunoblotting of Palm2 in developing mouse brain homogenates using all three antibodies, Neurofascin, and b-actin. The increase in Neurofascin protein reflects the development of nodes and paranodes with developmental myelination. (D) Immunostaining of Palm2 (green) and Caspr (magenta) in 1- and 7-month-old mouse optic nerve using the three commercial antibodies. Black arrowheads indicate paranodes, white arrowheads indicate nodes. (E) Immunostaining of Palm2 (green), Caspr (magenta), and AnkG (blue) in 1-month-old and 4-month-old mouse sciatic nerves using the three commercial Palm2 antibodies. (F-H) Percentage of nodes (white bars) or paranodes (black bars) in the optic nerve labeled using each Palm2 commercial antibody at different ages from 1 month to 7 months of age. (I) Summary of Palm2 antibodies with their antigen region and sensitivity at detecting Palm2 at AIS, nodes of Ranvier, and paranodes. Scale bars are indicated in each panel.

**Figure S3.**
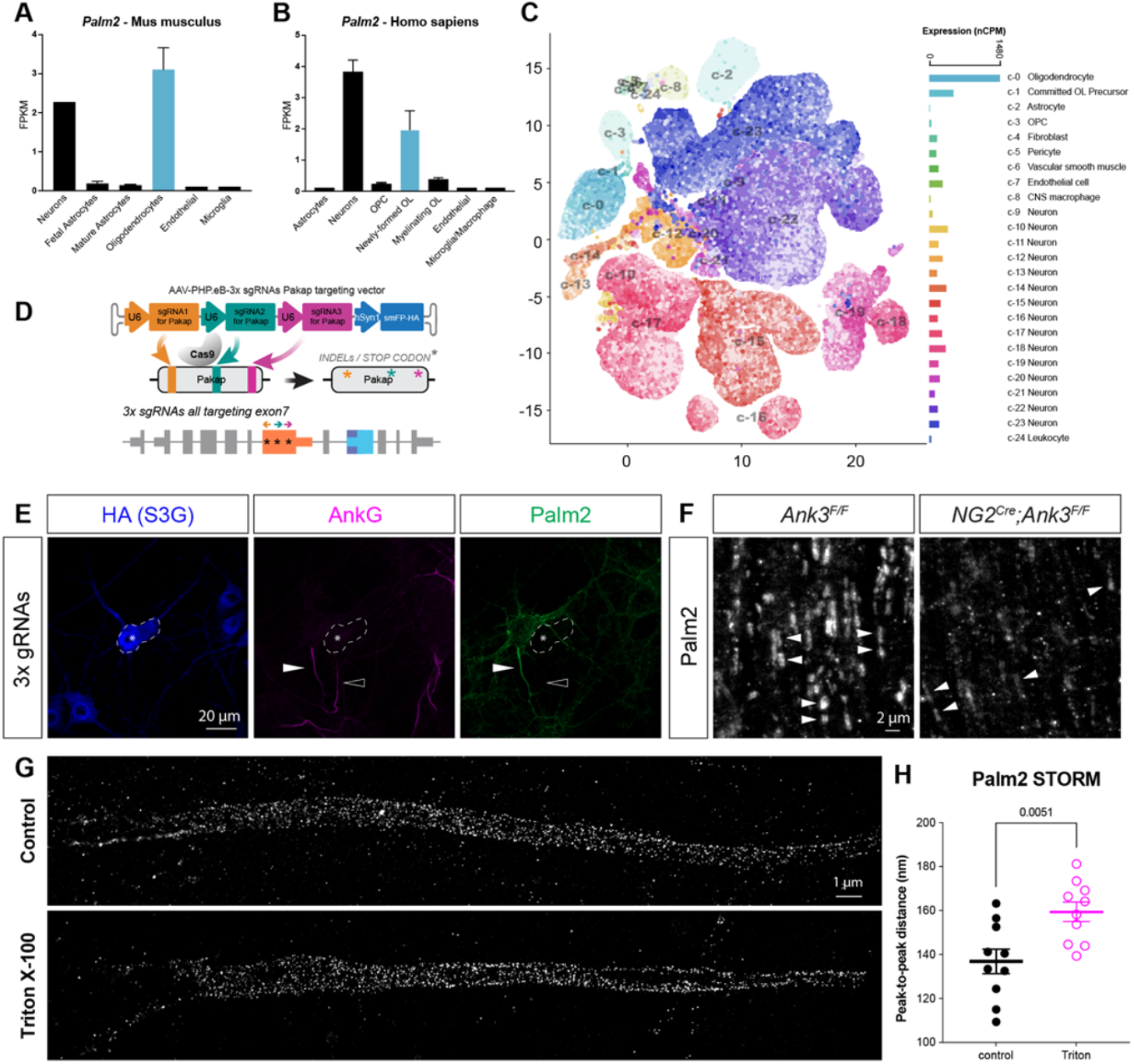
Cell-type specificity of Palm2 expression and dependence of exon 7 for AIS localization. (A-B) Cell-type specificity of *Palm2* expression in mouse (A) and human (B). Data are from the Brain-RNAseq database. Error bars represent mean ± SEM. (C) Cell-type specific expression of Palm2 from the brain single-cell atlas. (D) Schematic of AAV and triple (3x) gRNA-mediated disruption of exon 7 in *Palm2*. (E) Immunostaining of Palm2 (green) and AnkG (magenta) in HA-positive (blue) neurons transduced with AAV to express the 3x gRNA against exon 7 of *Palm2*. White arrowheads indicate the AIS of a non-transduced neuron with Palm2 colocalizing with AnkG. Black arrowheads indicate a transduced neuron with AnkG and no Palm2. Scale bar: 20 μm. (F) Immunostaining of Palm2 in *Ank3*^*F/F*^ and *NG2*^*Cre*^*;Ank3*^*F/F*^ optic nerves. Arrowheads indicate paranodes. Scale bar: 2 μm. (G) Super-resolution imaging of the nanoscale organization of Palm2 by stochastic optical reconstruction microscopy (STORM) in cultured neurons at DIV 10. Live neurons were treated with 0.5% Triton X-100 for 5 min at room temperature to extract detergent soluble pool of Palm2. Scale bar: 1 μm. (H) Peak-to-peak distance of fluorescence intensity across Palm2 immunofluorescent puncta measured by line-scan analysis along the AIS. Individual data point represents an AIS. N = 10 for both groups. Unpaired Student’s t-tests. Error bars represent mean ± SEM.

**Figure S4.**
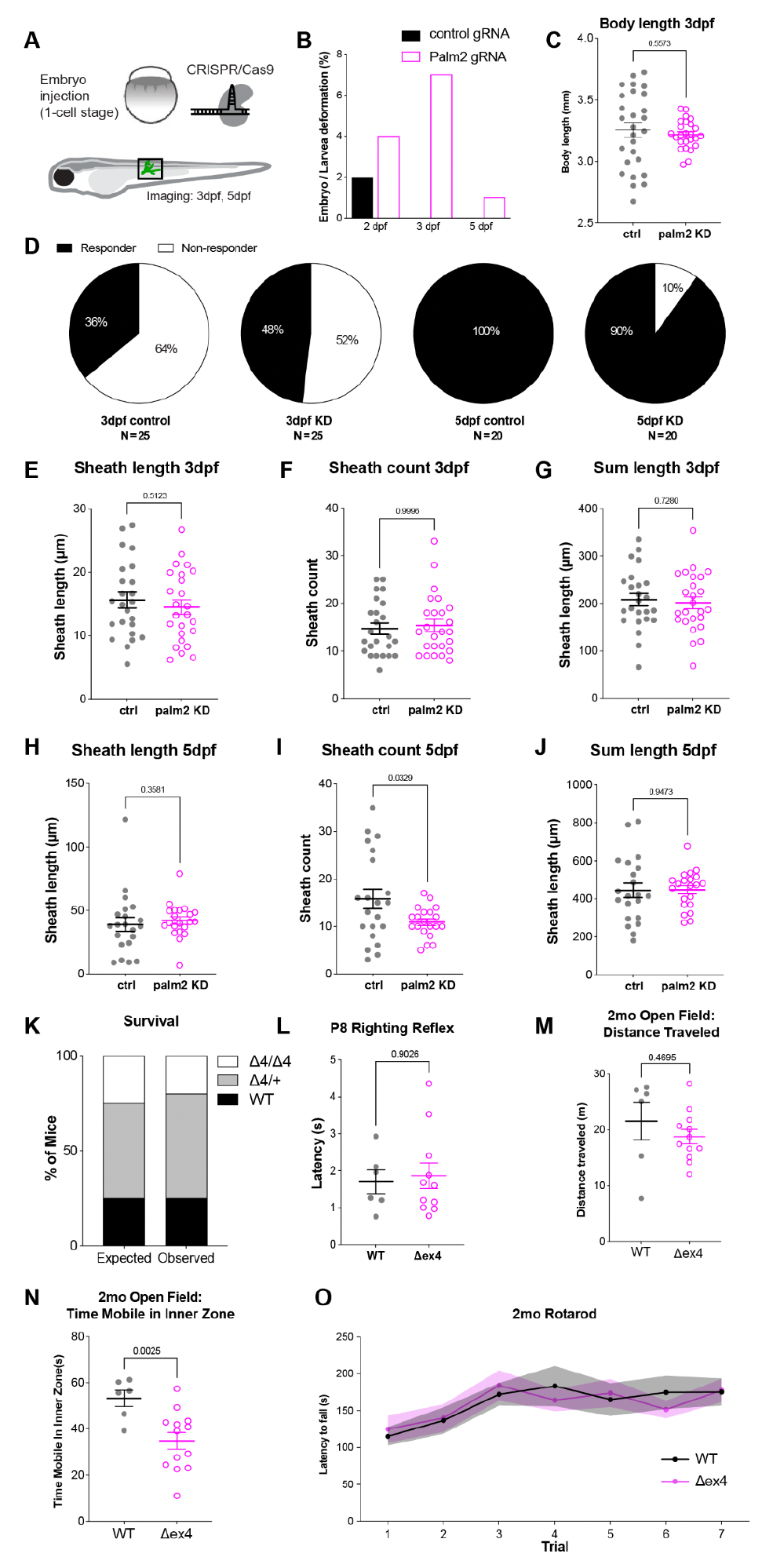
Characterization of survival, health, and early myelin development in Palm2 KD zebrafish larvae and survival and health in Palm2 KO mice. (A) Illustration of CRISPR/Cas9-mediated *Palm2* knock-down (KD) in zebrafish larvae. (B) Deformation rate in embryo/larvae at 2, 3, and 5 dpf following injection of control gRNAs (black) or Palm2 gRNAs (white). N = ∼100 larvae. (C) Body length at 3 dpf in control and Palm2 KD fish. Each data point represents one animal. N = 27 control and 25 KD. Mann Whitney U test. Error bars represent mean ± SEM. (D) Percentage of responders (black) and nonresponders (white) to gentle poke in control and Palm2 KD zebrafish larvae at 3 and 5 dpf. N= 25 and 20 fish at 3 and 5 dpf, respectively. (E-G) Myelin sheath length per cell (E), myelin sheath count per cell (F), and sum of sheath length per cell (G) in control and Palm2 KD groups at 3 dpf. Each data point represents one cell. N = 24 for control and 25 for KD. E: unpaired t-test. F: Kolmogorov-Smirnov test. G: unpaired t-test. (H-J) Myelin sheath length per cell (H), myelin sheath count per cell (I), and sum of sheath length per cell (J) in control and Palm2 KD groups at 5 dpf. Each data point represents one cell. N = 21 for each group, with 1 outlier excluded for KD group in J. I: Kolmogorov-Smirnov test. J: Welch’s t-test. K: Welch’s t-test. Error bars represent mean ± SEM. (K) Percentage of mice born with each genotype (WT: black, HT: gray, KO: white) as expected or observed. N = 69 mice. Chi-square analysis, χ^2^ = 0.7937, df = 2, p = 0.6725. (L) Latency to turn in the righting reflex test in P8 WT and KO mice. Each data point represents one animal. N = 6 for WT and 12 for KO with 1 outlier excluded. Mann Whitney U test. Error bars represent mean ± SEM. (M-N) Total distance traveled (M) in the open field apparatus and time spent mobile in the inner zone (N) by adult WT and KO mice. Each data point represents one animal. N = 6 for WT and 13 for KO with 1 outlier excluded for total distance traveled. No outlier was excluded for mobility in the inner zone. Welch’s t-test. Error bars represent mean ± SEM. (O) Latency to fall by adult WT and KO mice on the rotarod. N = 8 for WT and 16 for KO. Two-way ANOVA with repeated measurement and multiple comparison. Solid lines represent the mean and shaded area indicate ± SEM.

**Figure S5.**
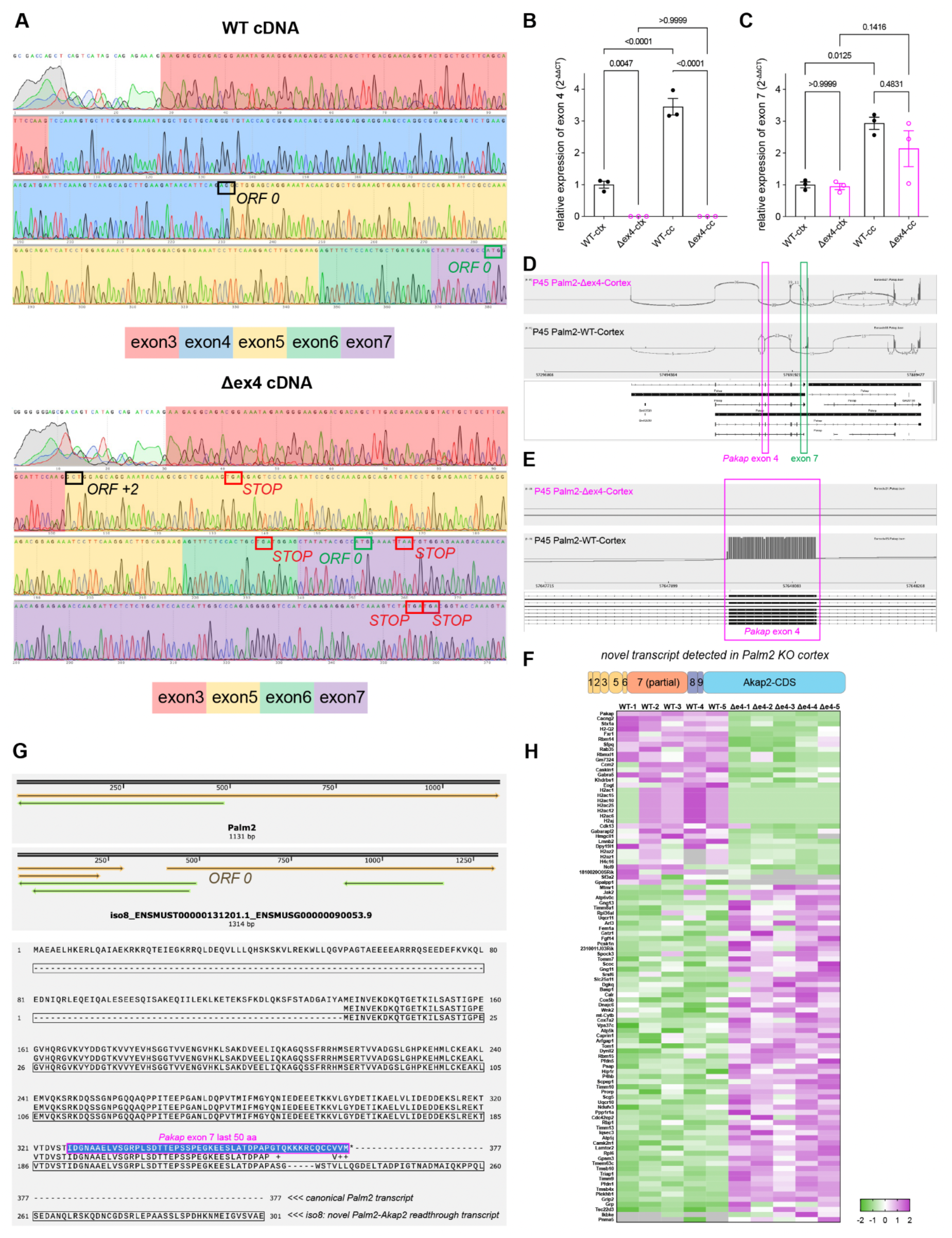
Validation of Palm2 KO mice and differential proteomes. (A) Sanger sequencing of cDNAs from adult Palm2 WT and KO mouse brains. (B-C) qRT-PCR of cDNAs from P45 Palm2 WT and KO mouse brains, detecting exons 4 (B) and 7 (C). Each data point represents an independent animal. N = 3 for both genotypes. Two-way ANOVA with multiple comparison. Error bars represent mean ± SEM. (D) Sashimi plots showing nanopore sequencing results of cDNA libraries from P45 Palm2 WT and KO mouse cortices. The region corresponding to exon 4 is labeled in magenta and exon 7 in green. (F) Zoomed-in view of (D), showing deletion of exon 4 in the Palm2 KO mice. (G) Schematic of a novel Palm2-Akap2 fusion transcript containing most of exon 7, detected only in Palm2 KO cortex. (H) Alignment of reference Palm2 cDNA and the novel isoform, showing inclusion of the majority of exon 7 except the coding sequence for the last 14 amino acids of Palm2. (I) Heatmap of top differentially expressed genes identified by DIA-mass spec using P45 brain membrane fractions from 5 WT mice and 5 KO mice. Color scale represents log_2_ iBAQ z-score, with green representing downregulated and magenta representing upregulated genes.

**Figure S6.**
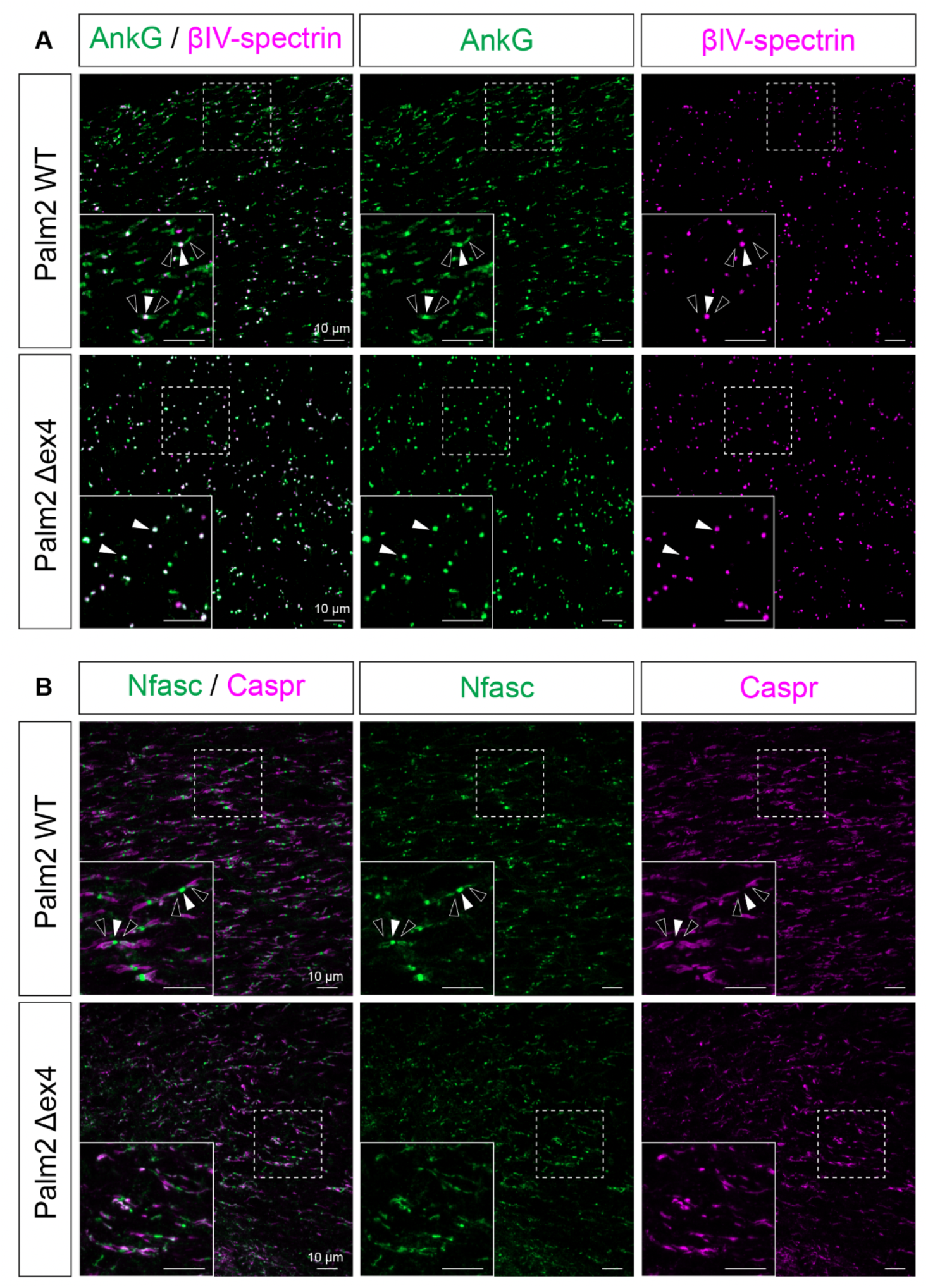
Disrupted paranodal AnkG and Nfasc in aged Palm2 KO optic nerves. (A) A low-magnification view of Fig. 8A, showing immunostaining for AnkG (green) and βIV-spectrin (magenta) in the optic nerves of 16-monthold Palm2 WT and KO mice. The dashed line square corresponds to the inset. Black arrowheads indicate paranodes and the white arrowhead indicates the node. (B) A low-magnification view of Fig. 8D, showing immunostaining for Nfasc (green) and Caspr (magenta) in the optic nerves of 16-month-old Palm2 WT and KO mice. The dashed line square corresponds to the inset. Scale bars: 10 μm.

## References

1. Jenkins, P.M., and Bender, K.J. (2024). Axon Initial Segment Structure and Function in Health and Disease. Physiol Rev. 10.1152/physrev.00030.2024.

2. Leterrier, C. (2018). The Axon Initial Segment: An Updated Viewpoint. J Neurosci 38, 2135–2145. 10.1523/JNEUROSCI.1922-17.2018.

3. Rasband, M.N., and Peles, E. (2021). Mechanisms of node of Ranvier assembly. Nat Rev Neurosci 22, 7–20. 10.1038/s41583-020-00406-8.

4. Eshed-Eisenbach, Y., Brophy, P.J., and Peles, E. (2023). Nodes of Ranvier in health and disease. J Peripher Nerv Syst 28 Suppl 3, S3–S11. 10.1111/jns.12568.

5. Buffington, S.A., and Rasband, M.N. (2011). The axon initial segment in nervous system disease and injury. Eur J Neurosci 34, 1609–1619.

6. Jenkins, P.M., Kim, N., Jones, S.L., Tseng, W.C., Svitkina, T.M., Yin, H.H., and Bennett, V. (2015). Giant ankyrin-G: a critical innovation in vertebrate evolution of fast and integrated neuronal signaling. Proc Natl Acad Sci U S A 112, 957–964. 10.1073/pnas.1416544112.

7. Chang, K.J., Zollinger, D.R., Susuki, K., Sherman, D.L., Makara, M.A., Brophy, P.J., Cooper, E.C., Bennett, V., Mohler, P.J., and Rasband, M.N. (2014). Glial ankyrins facilitate paranodal axoglial junction assembly. Nat Neurosci 17, 1673–1681. 10.1038/nn.3858.

8. Pillai, A.M., Thaxton, C., Pribisko, A.L., Cheng, J.G., Dupree, J.L., and Bhat, M.A. (2009). Spatiotemporal ablation of myelinating glia-specific neurofascin (Nfasc NF155) in mice reveals gradual loss of paranodal axoglial junctions and concomitant disorganization of axonal domains. J Neurosci Res 87, 1773–1793.

9. Zonta, B., Tait, S., Melrose, S., Anderson, H., Harroch, S., Higginson, J., Sherman, D.L., and Brophy, P.J. (2008). Glial and neuronal isoforms of Neurofascin have distinct roles in the assembly of nodes of Ranvier in the central nervous system. J Cell Biol 181, 1169–1177.

10. Sherman, D.L., Tait, S., Melrose, S., Johnson, R., Zonta, B., Court, F.A., Macklin, W.B., Meek, S., Smith, A.J., Cottrell, D.F., and Brophy, P.J. (2005). Neurofascins are required to establish axonal domains for saltatory conduction. Neuron 48, 737–742.

11. Hamdan, H., Lim, B.C., Torii, T., Joshi, A., Konning, M., Smith, C., Palmer, D.J., Ng, P., Leterrier, C., Oses-Prieto, J.A., et al. (2020). Mapping axon initial segment structure and function by multiplexed proximity biotinylation. Nature communications 11, 100. 10.1038/s41467-019-13658-5.

12. Gao, Y., Shonai, D., Trn, M., Zhao, J., Soderblom, E.J., Garcia Moreno, S.A., Gersbach, C.A., Wetsel, W.C., Dawson, G., Velmeshev, D., et al. (2024). Proximity analysis of native proteomes reveals phenotypic modifiers in a mouse model of autism and related neurodevelopmental conditions. Nature communications 15, 6801. 10.1038/s41467-024-51037-x.

13. Zhang, W., Fu, Y., Peng, L., Ogawa, Y., Ding, X., Rasband, A., Zhou, X., Shelly, M., Rasband, M.N., and Zou, P. (2023). Immunoproximity biotinylation reveals the axon initial segment proteome. Nature communications 14, 8201. 10.1038/s41467-023-44015-2.

14. Zhang, W., Palfini, V.L., Wu, Y., Ding, X., Melton, A.J., Gao, Y., Ogawa, Y., and Rasband, M.N. (2025). A hierarchy of PDZ domain scaffolding proteins clusters the Kv1 K(+) channel protein complex at the axon initial segment. Sci Adv 11, eadv1281. 10.1126/sciadv.adv1281.

15. Anderson, A.P., Kim, S., Melton, A.J., Ding, X., Zhang, W., Saltzman, A.B., Malovannaya, A., Rasband, M.N., and Gao, Y. (2025). A distinct PP2A subunit regulates local protein phosphorylation at the axon initial segment. Nature communications 16, 10850. 10.1038/s41467-025-66120-0.

16. Zhang, C., Joshi, A., Liu, Y., Sert, O., Haddix, S.G., Teliska, L.H., Rasband, A., Rodney, G.G., and Rasband, M.N. (2021). Ankyrin-dependent Na(+) channel clustering prevents neuromuscular synapse fatigue. Curr Biol 31, 3810–3819 e3814. 10.1016/j.cub.2021.06.052.

17. Zhou, D., Lambert, S., Malen, P.L., Carpenter, S., Boland, L.M., and Bennett, V. (1998). AnkyrinG is required for clustering of voltage-gated Na channels at axon initial segments and for normal action potential firing. J Cell Biol 143, 1295–1304.

18. Ho, T.S., Zollinger, D.R., Chang, K.J., Xu, M., Cooper, E.C., Stankewich, M.C., Bennett, V., and Rasband, M.N. (2014). A hierarchy of ankyrin-spectrin complexes clusters sodium channels at nodes of Ranvier. Nat Neurosci 17, 1664–1672. 10.1038/nn.3859.

19. Nakashiba, T., Sugiyama, T., Iwano, S., Iwama, M., Yoshiki, A., Miyawaki, A., and Abe, K. (2026). Assessing leaky expression of Cre-dependent DNA constructs in the mouse genome using sensitive bioluminescent reporters. Cell Rep Methods 6, 101369. 10.1016/j.crmeth.2026.101369.

20. Yang, Z., Zhang, Y., Fang, Y., Zhang, Y., Du, J., Shen, X., Zhang, K., Zou, P., and Chen, Z. (2025). Spatial barcoding reveals reaction radii and contact-dependent mechanism of proximity labeling. Nat Chem Biol. 10.1038/s41589-025-02086-w.

21. Haddix, S.G., Zhang, C., Liu, Y., Oses-Prieto, J., Burlingame, A.L., and Rasband, M.N. (2026). The perijunctional zone is a molecularly distinct muscle subdomain altered in Duchenne muscular dystrophy. Nature communications 17. 10.1038/s41467-026-73525-y.

22. Hultqvist, G., Ocampo Daza, D., Larhammar, D., and Kilimann, M.W. (2012). Evolution of the vertebrate paralemmin gene family: ancient origin of gene duplicates suggests distinct functions. PLoS One 7, e41850. 10.1371/journal.pone.0041850.

23. Deng, D.X., Li, C.Y., Zheng, Z.Y., Wen, B., Liao, L.D., Zhang, X.J., Li, E.M., and Xu, L.Y. (2023). Prenylated PALM2 Promotes the Migration of Esophageal Squamous Cell Carcinoma Cells Through Activating Ezrin. Molecular & cellular proteomics : MCP 22, 100593. 10.1016/j.mcpro.2023.100593.

24. Macarron-Palacios, V., Hubrich, J., do Rego Barros Fernandes Lima, M.A., Metzendorf, N.G., Kneilmann, S., Trapp, M., Acuna, C., Patrizi, A., D’Este, E., and Kilimann, M.W. (2025). Paralemmin-1 controls the nanoarchitecture of the neuronal submembrane cytoskeleton. Sci Adv 11, eadt3724. 10.1126/sciadv.adt3724.

25. Petropavlovskiy, A.A., Kogut, J.A., Leekha, A., Townsend, C.A., and Sanders, S.S. (2021). A sticky situation: regulation and function of protein palmitoylation with a spotlight on the axon and axon initial segment. Neuronal Signal 5, NS20210005. 10.1042/NS20210005.

26. Leterrier, C., Potier, J., Caillol, G., Debarnot, C., Rueda Boroni, F., and Dargent, B. (2015). Nanoscale Architecture of the Axon Initial Segment Reveals an Organized and Robust Scaffold. Cell Rep 13, 2781–2793. 10.1016/j.celrep.2015.11.051.

27. Gao, Y., Hisey, E., Bradshaw, T.W.A., Erata, E., Brown, W.E., Courtland, J.L., Uezu, A., Xiang, Y., Diao, Y., and Soderling, S.H. (2019). Plug-and-Play Protein Modification Using Homology-Independent Universal Genome Engineering. Neuron 103, 583–597e588. 10.1016/j.neuron.2019.05.047.

28. Dowd, G., and Ogawa, Y. (2026). Protocol for one step design of a triple sgRNA-based CRISPR/Cas9 construct for neuronal gene knockout in the mouse brain. STAR Protoc 7, 104715. 10.1016/j.xpro.2026.104715.

29. Zhang, Y., Chen, K., Sloan, S.A., Bennett, M.L., Scholze, A.R., O’Keeffe, S., Phatnani, H.P., Guarnieri, P., Caneda, C., Ruderisch, N., et al. (2014). An RNA-sequencing transcriptome and splicing database of glia, neurons, and vascular cells of the cerebral cortex. J Neurosci 34, 11929–11947. 10.1523/JNEUROSCI.1860-14.2014.

30. Zhang, Y., Sloan, S.A., Clarke, L.E., Caneda, C., Plaza, C.A., Blumenthal, P.D., Vogel, H., Steinberg, G.K., Edwards, M.S., Li, G., et al. (2016). Purification and Characterization of Progenitor and Mature Human Astrocytes Reveals Transcriptional and Functional Differences with Mouse. Neuron 89, 37–53. 10.1016/j.neuron.2015.11.013.

31. Hu, B., Copeland, N.G., Gilbert, D.J., Jenkins, N.A., and Kilimann, M.W. (2001). The paralemmin protein family: identification of paralemmin-2, an isoform differentially spliced to AKAP2/AKAP-KL, and of palmdelphin, a more distant cytosolic relative. Biochem Biophys Res Commun 285, 1369–1376. 10.1006/bbrc.2001.5329.

32. Weissbach, S., Milkovits, J., Pastore, S., Heine, M., Gerber, S., and Todorov, H. (2024). Cortexa: a comprehensive resource for studying gene expression and alternative splicing in the murine brain. BMC Bioinformatics 25, 293. 10.1186/s12859-024-05919-y.

33. Weyn-Vanhentenryck, S.M., Feng, H., Ustianenko, D., Duffie, R., Yan, Q., Jacko, M., Martinez, J.C., Goodwin, M., Zhang, X., Hengst, U., et al. (2018). Precise temporal regulation of alternative splicing during neural development. Nature communications 9, 2189. 10.1038/s41467-018-04559-0.

34. Xu, K., Zhong, G., and Zhuang, X. (2013). Actin, Spectrin, and Associated Proteins Form a Periodic Cytoskeletal Structure in Axons. Science 339, 30495–30501.

35. Schafer, D.P., Bansal, R., Hedstrom, K.L., Pfeiffer, S.E., and Rasband, M.N. (2004). Does paranode formation and maintenance require partitioning of neurofascin 155 into lipid rafts? J Neurosci 24, 3176–3185.

36. Ren, Q., and Bennett, V. (1998). Palmitoylation of neurofascin at a site in the membrane-spanning domain highly conserved among the L1 family of cell adhesion molecules. J Neurochem 70, 1839–1849.

37. He, M., Abdi, K.M., and Bennett, V. (2014). Ankyrin-G palmitoylation and betaII-spectrin binding to phosphoinositide lipids drive lateral membrane assembly. J Cell Biol 206, 273–288. 10.1083/jcb.201401016.

38. Ogawa, Y., Schafer, D.P., Horresh, I., Bar, V., Hales, K., Yang, Y., Susuki, K., Peles, E., Stankewich, M.C., and Rasband, M.N. (2006). Spectrins and ankyrinB constitute a specialized paranodal cytoskeleton. J Neurosci 26, 5230–5239.

39. Park, H.C., Kim, C.H., Bae, Y.K., Yeo, S.Y., Kim, S.H., Hong, S.K., Shin, J., Yoo, K.W., Hibi, M., Hirano, T., et al. (2000). Analysis of upstream elements in the HuC promoter leads to the establishment of transgenic zebrafish with fluorescent neurons. Dev Biol 227, 279–293. 10.1006/dbio.2000.9898.

40. Riedl, J., Crevenna, A.H., Kessenbrock, K., Yu, J.H., Neukirchen, D., Bista, M., Bradke, F., Jenne, D., Holak, T.A., Werb, Z., et al. (2008). Lifeact: a versatile marker to visualize F-actin. Nature methods 5, 605–607. 10.1038/nmeth.1220.

41. Susuki, K., Chang, K.J., Zollinger, D.R., Liu, Y., Ogawa, Y., Eshed-Eisenbach, Y., Dours-Zimmermann, M.T., Oses-Prieto, J.A., Burlingame, A.L., Seidenbecher, C.I., et al. (2013). Three mechanisms assemble central nervous system nodes of Ranvier. Neuron 78, 469–482.

42. Amor, V., Zhang, C., Vainshtein, A., Zhang, A., Zollinger, D.R., Eshed-Eisenbach, Y., Brophy, P.J., Rasband, M.N., and Peles, E. (2017). The paranodal cytoskeleton clusters Na(+) channels at nodes of Ranvier. eLife 6. 10.7554/eLife.21392.

43. Bissen, D., Foss, F., and Acker-Palmer, A. (2019). AMPA receptors and their minions: auxiliary proteins in AMPA receptor trafficking. Cell Mol Life Sci 76, 2133–2169. 10.1007/s00018-019-03068-7.

44. Anggono, V., and Huganir, R.L. (2012). Regulation of AMPA receptor trafficking and synaptic plasticity. Curr Opin Neurobiol 22, 461–469. 10.1016/j.conb.2011.12.006.

45. Ding, X., Curtis, J.R., Xing, Y., Wu, Y., Peles, E., and Rasband, M.N. (2025). OASIS: in vivo AAV-mediated transduction and genome editing of adult oligodendrocytes. bioRxiv. 10.1101/2025.10.20.683551.

46. Ding, X., Wu, Y., Vainshtein, A., Rodriguez, V., Ricco, E., Okoh, J.T., Liu, Y., Kraushaar, D.C., Peles, E., and Rasband, M.N. (2024). Age-dependent regulation of axoglial interactions and behavior by oligodendrocyte AnkyrinG. Nature communications 15, 10865. 10.1038/s41467-024-55209-7.

47. Susuki, K., Zollinger, D.R., Chang, K.J., Zhang, C., Huang, C.Y., Tsai, C.R., Galiano, M.R., Liu, Y., Benusa, S.D., Yermakov, L.M., et al. (2018). Glial betaII Spectrin Contributes to Paranode Formation and Maintenance. J Neurosci 38, 6063–6075. 10.1523/JNEUROSCI.3647-17.2018.

48. Stevens, S.R., and Rasband, M.N. (2022). Pleiotropic Ankyrins: Scaffolds for Ion Channels and Transporters. Channels (Austin) 16, 216–229. 10.1080/19336950.2022.2120467.

49. Tait, S., Gunn-Moore, F., Collinson, J.M., Huang, J., Lubetzki, C., Pedraza, L., Sherman, D.L., Colman, D.R., and Brophy, P.J. (2000). An oligodendrocyte cell adhesion molecule at the site of assembly of the paranodal axo-glial junction. J Cell Biol 150, 657–666.

50. Eshed, Y., Feinberg, K., Poliak, S., Sabanay, H., Sarig-Nadir, O., Spiegel, I., Bermingham, J.R., Jr., and Peles, E. (2005). Gliomedin mediates schwann cell-axon interaction and the molecular assembly of the nodes of ranvier. Neuron 47, 215–229.

51. Brilkova, M., Nigri, M., Kumar, H.S., Moore, J., Mantovani, M., Keller, C., Grimm, A., Eckert, A., Shcherbakov, D., Akbergenov, R., et al. (2022). Error-prone protein synthesis recapitulates early symptoms of Alzheimer disease in aging mice. Cell Rep 40, 111433. 10.1016/j.celrep.2022.111433.

52. He, M., Jenkins, P., and Bennett, V. (2012). Cysteine 70 of ankyrin-G is S-palmitoylated and is required for function of ankyrin-G in membrane domain assembly. J Biol Chem 287, 43995–44005. 10.1074/jbc.M112.417501.

53. Peles, E., Nativ, M., Lustig, M., Grumet, M., Schilling, J., Martinez, R., Plowman, G.D., and Schlessinger, J. (1997). Identification of a novel contactin-associated transmembrane receptor with multiple domains implicated in protein-protein interactions. EMBO J 16, 978–988.

54. Berglund, E.O., Boyle, M.E.T., Murai, K.K., Peles, E., Weber, L., and Ranscht, B. (2000). Contactin regulates axon Schwann cell interactions at the paranode in myelinated peripheral nerve. Society for Neuroscience Abstracts, 407.409.

55. Bhat, M.A., Rios, J.C., Lu, Y., Garcia-Fresco, G.P., Ching, W., St Martin, M., Li, J., Einheber, S., Chesler, M., Rosenbluth, J., et al. (2001). Axon-glia interactions and the domain organization of myelinated axons requires neurexin IV/Caspr/Paranodin. Neuron 30, 369–383.

56. Menegoz, M., Gaspar, P., Le Bert, M., Galvez, T., Burgaya, F., Palfrey, C., Ezan, P., Arnos, F., and Girault, J.A. (1997). Paranodin, a glycoprotein of neuronal paranodal membranes. Neuron 19, 319–331.

57. Ohno, N., Terada, N., Yamakawa, H., Komada, M., Ohara, O., Trapp, B.D., and Ohno, S. (2006). Expression of protein 4.1G in Schwann cells of the peripheral nervous system. J Neurosci Res.

58. Hengel, H., Magee, A., Mahanjah, M., Vallat, J.M., Ouvrier, R., Abu-Rashid, M., Mahamid, J., Schule, R., Schulze, M., Krageloh-Mann, I., et al. (2017). CNTNAP1 mutations cause CNS hypomyelination and neuropathy with or without arthrogryposis. Neurol Genet 3, e144. 10.1212/NXG.0000000000000144.

59. Martin-Aguilar, L., Lleixa, C., and Pascual-Goni, E. (2022). Autoimmune nodopathies, an emerging diagnostic category. Curr Opin Neurol 35, 579–585. 10.1097/WCO.0000000000001107.

60. Hashemi, S.M., Hund, T.J., and Mohler, P.J. (2009). Cardiac ankyrins in health and disease. J Mol Cell Cardiol 47, 203–209. 10.1016/j.yjmcc.2009.04.010.

61. Kizhatil, K., and Bennett, V. (2004). Lateral membrane biogenesis in human bronchial epithelial cells requires 190-kDa ankyrin-G. J Biol Chem 279, 16706–16714.

62. Li, J., Miramontes, T.G., Czopka, T., and Monk, K.R. (2024). Synaptic input and Ca(2+) activity in zebrafish oligodendrocyte precursor cells contribute to myelin sheath formation. Nat Neurosci 27, 219–231. 10.1038/s41593-023-01553-8.

63. Ovesny, M., Krizek, P., Borkovec, J., Svindrych, Z., and Hagen, G.M. (2014). ThunderSTORM: a comprehensive ImageJ plug-in for PALM and STORM data analysis and super-resolution imaging. Bioinformatics 30, 2389–2390. 10.1093/bioinformatics/btu202.

64. Saltzman, A.B., Leng, M., Bhatt, B., Singh, P., Chan, D.W., Dobrolecki, L., Chandrasekaran, H., Choi, J.M., Jain, A., Jung, S.Y., et al. (2018). gpGrouper: A Peptide Grouping Algorithm for Gene-Centric Inference and Quantitation of Bottom-Up Proteomics Data. Molecular & cellular proteomics : MCP 17, 2270–2283. 10.1074/mcp.TIR118.000850.

65. Ritchie, M.E., Phipson, B., Wu, D., Hu, Y., Law, C.W., Shi, W., and Smyth, G.K. (2015). limma powers differential expression analyses for RNA-sequencing and microarray studies. Nucleic Acids Res 43, e47. 10.1093/nar/gkv007.

66. Li, H. (2021). New strategies to improve minimap2 alignment accuracy. Bioinformatics 37, 4572–4574. 10.1093/bioinformatics/btab705.

67. Tang, A.D., Felton, C., Hrabeta-Robinson, E., Volden, R., Vollmers, C., and Brooks, A.N. (2024). Detecting haplotype-specific transcript variation in long reads with FLAIR2. Genome Biol 25, 173. 10.1186/s13059-024-03301-y.

